# Env-antibody coevolution identifies B cell priming as the principal bottleneck to HIV-1 V2 apex broadly neutralizing antibody development

**DOI:** 10.1101/2025.05.03.652068

**Authors:** Rumi Habib, Ryan S. Roark, Hui Li, Andrew Jesse Connell, Michael P. Hogarty, Kshitij Wagh, Shuyi Wang, Lorie Marchitto, Ashwin N. Skelly, John W. Carey, Kirsten J. Sowers, Kasirajan Ayyanathan, Samantha J. Plante, Frederic Bibollet-Ruche, Younghoon Park, Colby J. Agostino, Ajay Singh, Christian L. Martella, Emily Lewis, Juliette M. Rando, Neha Chohan, Jinery Lora, Wenge Ding, Mary S. Campion, Chengyan Zhao, Weimin Liu, Yingying Li, Xuduo Li, Bo Liang, Rohan Roy Chowdhury, Khaled Amereh, Elizabeth Van Itallie, Zizhang Sheng, Amrit R. Ghosh, Katharine J. Bar, Wilton B. Williams, Kevin Wiehe, Kevin O. Saunders, Robert J. Edwards, Derek W. Cain, Mark Lewis, Facundo D. Batista, Dennis R. Burton, Raiees Andrabi, Daniel W. Kulp, Barton F. Haynes, Bette Korber, Lawrence Shapiro, Peter D. Kwong, Beatrice H. Hahn, George M. Shaw

**Affiliations:** Departments of Medicine and Microbiology, Perelman School of Medicine, University of Pennsylvania, Philadelphia, PA 19104, USA; Vaccine and Immunotherapy Center, The Wistar Institute, Philadelphia, PA 19104, USA; Aaron Diamond AIDS Research Center, Vagelos College of Physicians and Surgeons, Columbia University, New York, NY 10032, USA; Department of Biochemistry and Molecular Biophysics, Columbia University, New York, NY 10027, USA; Duke Human Vaccine Institute, Duke University School of Medicine, Durham, NC 27710, USA; Departments of Microbiology and Immunology, Harvard Medical School; The Ragon Institute of MGH, MIT and Harvard, Cambridge, MA 02139, USA; Department of Biology, Massachusetts Institute of Technology, Cambridge, MA 02139, USA; Bioqual, Inc., Rockville, MD 20850, USA; Department of Immunology and Microbiology, The Scripps Research Institute, La Jolla, CA 92037, USA; New Mexico Consortium, Los Alamos, NM 87545, USA

## Abstract

Broadly neutralizing antibodies (bNAbs) are rarely elicited during HIV-1 infection. To identify obstacles to bNAb development, we longitudinally studied 122 rhesus macaques infected by one of 16 different simian-human immunodeficiency viruses (SHIVs). We identified V2 apex as the most common bNAb target and a subset of Envs that preferentially elicited these antibodies. In 10 macaques, we delineated Env-antibody coevolution from B cell priming to bNAb development. Antibody phylogenies revealed permissive developmental pathways guided by evolving Envs that contained few mutations in or near the V2 apex C-strand, which were a sensitive indicator of apex-targeted responses. The absence of such mutations reflected a failure in bNAb priming. These results indicate that efficiency of B cell priming, and not complexities in Env-guided affinity maturation, is the primary obstacle to V2 apex bNAb elicitation in SHIV-infected macaques and identify specific HIV-1 Envs to advance as novel vaccine platforms.

**One sentence summary:** B cell priming is the primary bottleneck to HIV-1 V2 apex bNAb elicitation.

## INTRODUCTION

A primary objective of HIV-1 vaccine design is the reproducible elicitation of broadly neutralizing antibodies (bNAbs) to the HIV-1 envelope (Env) glycoprotein. This is a challenging goal as potent bNAbs are rarely elicited during HIV-1 infection and generally require years to develop (*1–7*). A major question in the fields of HIV-1 immunology and vaccinology is why some individuals develop bNAbs and others do not. While several correlates of bNAb induction have been identified, including viral load, CD4 T cell count, superinfection, viral diversity, length of infection, integrity of the Env glycan shield, and Env genealogy (*6–12*), the mechanistic bottlenecks underlying the stochasticity in bNAb development during HIV-1 infection remain unknown. Here we provide evidence that in the case of V2 apex bNAbs, priming of germline B cells capable of bNAb development is the rate-limiting step.

HIV-1 bNAbs in humans target one of six canonical epitope sites that are generally conserved across the global diversity of HIV-1 strains and subtypes. These include the V2 apex, V3-glycan high mannose patch, CD4-binding site (CD4bs), silent face (SF), gp120/gp41 interface region including the fusion peptide (FP), and the membrane-proximal external region (MPER). Surface exposed glycans targeted by Fab-dimerized antibodies represent an additional set of bNAb epitopes (*13–16*). bNAbs targeting each of these sites exhibit properties that reflect the unique obstacles to antibody access inherent to the heavily glycosylated viral Env trimer as well as underlying features of the human B cell immunoglobulin gene repertoire (*13–15*). The most common bNAb epitope specificity in people living with HIV-1 is the V3-glycan high mannose patch, followed by V2 apex (*4, 6, 17*). In rhesus macaques (RMs) infected with chimeric simian-human immunodeficiency viruses (SHIVs) expressing HIV-1 Env ectodomains (*18, 19*), the relative frequency of different bNAb epitope specificities is unknown.

A mechanistic understanding of bNAb elicitation in people living with HIV-1 and in SHIV-infected monkeys can guide vaccine design in multiple ways. First, the inference of authentic bNAb unmutated common ancestors (UCAs) by B cell lineage tracing can provide important antibody sequence information, structures, and molecular reagents that are essential for immunogen design as they comprise representative examples of B cell receptors that effective priming immunogens must engage (*15, 20*). UCAs inferred with high confidence from deep longitudinal immunoglobulin sequence datasets represent authentic bNAb precursors, whereas germline inferences that revert only gene-templated antibody regions (referred to as reverted or inferred germline receptors, or iGLs) are invariably confounded by uncertainty at non-templated V-D and D-J junctions. Second, tracing viral evolution can identify Env immunotypes that bind bNAb UCAs and lineage intermediates and then guide antibody affinity maturation along desired pathways (*21–26*). Third, the identification of shared features among different UCAs of a bNAb class and their evolved progeny can inform both immunogen design and clinical trial assessment by defining success criteria for priming and boosting B cell lineages that possess bNAb potential. Finally, studies of Env-antibody coevolution can point to obstacles or rate-limiting steps in bNAb development during the course of infection, and by inference, following vaccination (*20–24*). To address these issues, we analyzed V2 apex bNAb development in a prospective cohort of 122 SHIV-infected RMs to understand why such antibodies are so rarely elicited during infection, and how they might be more efficiently induced by vaccination.

SHIV-infected RMs are a favorable model to study bNAb development. The model allows for regular and controlled sampling of blood and lymphoid tissues as well as an analysis and comparison of Env-antibody coevolution that results from the same Env replicating in multiple outbred animals, some of which develop bNAbs and others that do not (*27*). We can thus look for shared patterns of B cell priming and antibody evolution leading to breadth, and at the same time, identify patterns of Env evolution driving that breadth. We previously showed that V2 apex bNAbs from SHIV-infected RMs exhibit striking structural homology to prototypic human V2 apex bNAbs (*27, 28*). In the present report, we use next-generation sequencing (NGS) of B cells and single genome sequencing (SGS) plus NGS of plasma virion RNA to define Env-antibody coevolution leading to bNAb development, beginning with Env engagement of the naïve germline B cell followed by sequential rounds of affinity maturation resulting in neutralization breadth and potency. We estimate when each bNAb lineage precursor B cell was triggered, infer with high confidence the precise heavy and light chain sequences of these UCAs, and identify intermediate stage antibodies on the path toward neutralization breadth. Simultaneously, we identify and characterize Envs that were likely responsible for initial B cell priming along with Env escape variants that guided B cell lineage maturation to full breadth and potency. Finally, we augment these results with analysis of data from human cohort studies of HIV-1 infection to identify generalizable features, bottlenecks, and “rules” of V2 apex bNAb induction applicable to both humans and rhesus.

## RESULTS

### V2 apex is the most common bNAb target in SHIV-infected RMs

Most HIV-1 infections are established by a single transmitted/founder (T/F) virus (*29*). Because of the extraordinary genetic diversity of HIV-1, nearly every infected individual acquires a unique strain. This, along with the fact that most infections do not lead to bNAb development, has made identifying any particular Env as a promising vaccine candidate challenging. Nonetheless, Trkola and colleagues found neutralization breadth to be associated with certain donor-recipient transmission pairs in a cohort of epidemiologically linked HIV-1 infections, leading to the concept of “bNAb imprinting Envs” (*7*). In support of this concept, we found that certain HIV-1 Envs that had elicited V2 apex, V3-glycan, and CD4bs bNAbs in several human study participants induced the same antibody specificities in RMs when expressed as SHIVs, with the molecular patterns of Env-antibody coevolution in monkeys recapitulating those found in humans (*27, 30*). These findings led us to postulate that certain HIV-1 Envs may have a propensity for eliciting particular bNAb specificities, a hypothesis that we formally examine in the current study.

The SHIV model is uniquely poised to evaluate the concept of bNAb imprinting since it allows for the infection of multiple RMs with SHIVs expressing the same Env. In the present study, we infected 122 rhesus macaques with SHIVs bearing any one of 16 different primary HIV-1 Envs (*18, 19*) and followed them for up to six years (mean of 2 years, median of 1.7 years) for the development of neutralization breadth (**table S1**). The Envs selected for SHIV construction were chosen based on criteria that we hypothesized might increase the probability that they would elicit bNAbs, as previously described (*19*). These criteria included Envs that in humans induced bNAbs targeting V2 apex, V3 glycan, or CD4bs epitopes (*12, 27*); Envs that were previously shown to bind germline-reverted V2 apex bNAb precursors (*31–35*); Envs that corresponded to T/F sequences from acutely-infected human participants (*29*); and Envs that represented all major HIV-1 group M subtypes (*18, 19*). All 122 macaques became persistently infected with setpoint plasma virus loads of 10^2^-10^7^ vRNA/ml (geometric mean = 1.4 × 10^4^ vRNA/ml) (**table S1**). Seven SHIV-infected RMs experienced rapid disease progression and died or were euthanized within months of infection because of opportunistic infections, AIDS-related malignancies or generalized wasting (**table S1**). Each of these monkeys had extremely high setpoint plasma viral loads (∼10^7^ vRNA/ml) with little or no detectable anti-SHIV antibody responses as assessed by ELISA, western immunoblot and neutralization assays. This rapid progressor phenotype is similar to that reported for a subset of SIVmac-infected monkeys (*36, 37*). Rapid progressor animals were deemed unevaluable for bNAb induction but served as valuable controls for Env sequence evolution in the absence of adaptive immune selective pressure. Of the remaining RMs, 113/115 (98%) developed autologous, strain-specific NAbs with 50% plasma inhibitory dilution (ID_50_) titers against the infecting SHIV strain ranging from 1:24-1:120,000 (geometric mean = 1:750), generally within 3-6 months of infection (**table S1**).

Twenty-five of 115 (22%) monkeys developed neutralization breadth defined as ID_50_ titers of ≥1:80 against two or more members of an 18-strain tier 1B/2 heterologous virus panel (**Fig. 1A, table S1**). Epitope specificity was confirmed by differential neutralization against heterologous viruses containing site-directed mutations in canonical bNAb epitopes, by EMPEM analysis of heterologous virus-antibody binding, or by isolation of bNAb mAbs followed by cryoEM structural analysis as previously described (**fig. S1**) (*27, 28, 30*). We chose a low threshold for neutralization breadth combined with a stringent threshold for bNAb epitope confirmation to ensure sensitive detection of canonical bNAb lineages. We did this because a similarly wide range in neutralization breadth and potency would be expected in polyclonal responses to candidate vaccines where the summation of narrow but potent bNAb responses could contribute significantly to clinical protection (*38–41*). Based on these criteria, we identified 18 RMs with V2 apex bNAbs, four with V3 glycan high mannose patch bNAbs, two with silent face bNAbs, and one each with FP and CD4bs bNAbs (**Fig. 1A**). One RM developed two distinct bNAb lineages targeting both V2 apex and FP. If we increased the stringency of the definition of neutralization breadth to require ID_50_ titers of ≥1:80 against four or more members of the 18-strain virus test panel, then eight, three, one, one and one monkeys had confirmed V2, V3, SF, FP and CD4bs bNAbs, respectively, or 13/115 (11.3%) overall (**table S1**). This frequency of verified bNAbs in SHIV-infected RMs ranging from 11-22% is generally similar to that observed in people living with HIV (*4, 6*).

**Fig. 1.**
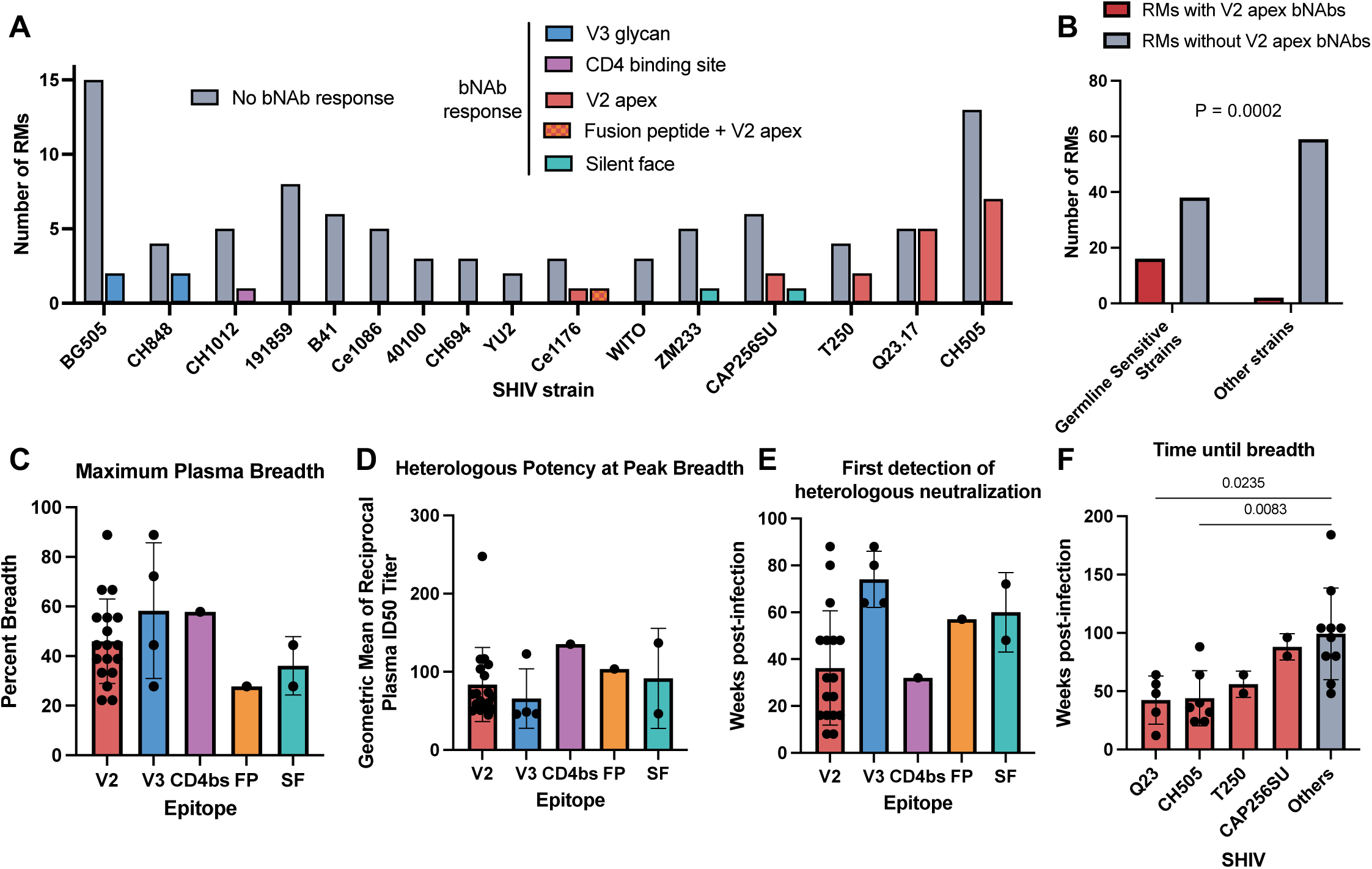
HIV-1 Envs with a predilection for V2 apex bNAb induction. (**A**) Numbers of RMs infected with SHIV strains expressing different primary HIV-1 Envs that elicited V2 apex, V3-glycan, fusion peptide, CD4-binding site, or silent face bNAbs. RMs with bNAbs were defined as RMs with ID_50_ titers of ≥1:80 against two or more members of an 18-strain tier 1B/2 heterologous virus panel. (**B**) Frequency of V2 apex bNAb elicitation by germline-sensitive SHIV strains expressing HIV-1 Envs that were previously shown to be sensitive to neutralization by V2 apex bNAb inferred germline (iGL) precursor antibodies or iGL-intermediate antibodies compared to all other SHIV strains (p = 0.0002; Fisher’s Exact test). (**C**) Maximum plasma neutralization breadth at ID_50_ titer ≥1:20 grouped by bNAb specificity. (**D**) Geometric means of heterologous plasma ID_50_ neutralization titers of RMs at the timepoint at which they showed maximum breadth. (**E**) Time (in weeks post-infection) until the first detection of heterologous plasma neutralization grouped by bNAb specificity. (**F**) Time (in weeks post-infection) until the acquisition of neutralization breadth grouped by infecting SHIV strain. Significance in panels **C**-**F** was determined by Kruskal–Wallis test followed by Dunn’s multiple comparisons test. Only significant differences (p < 0.05) are indicated. Error bars in **C**-**F** indicate one standard deviation from the mean.

### Identification of HIV-1 Envs with a predilection for V2 apex bNAb induction

V2 apex bNAbs were elicited by SHIVs Q23.17 (5 of 10 RMs), CH505 (7 of 20 RMs), CAP256SU (2 of 9 RMs), T250 (2 of 6 RMs) and Ce1176 (2 of 5 RMs) but by none of the other 11 SHIVs (**Fig. 1A**). This uneven distribution of bNAb elicitation indicated a significant association between particular Envs and V2 apex bNAb elicitation (Fisher’s Exact Test, p=0.011). Monte Carlo repeated random sampling indicated that the restriction of V2 apex bNAb elicitation to five SHIVs was highly significant (p < 0.0001, see Supplemental Text). Interestingly, the Envs that elicited V2 apex bNAbs — Q23.17, CH505, T250, CAP256SU — were significantly enriched among a group Envs previously shown to bind or be neutralized by germline-reverted human V2 apex bNAbs (16 of 54 versus 2 of 61; p = 0.0002, Fisher’s Exact Test; see **Fig. 1B**) (*31–35*).

Maximum neutralization breadth in the plasma was not significantly different among RMs with different bNAb specificities, with V2, V3, CD4bs, FP, and SF bNAbs achieving 50%, 61%, 63%, 32%, and 50% breadth, respectively, against the 18-virus panel (p > 0.05 by Kruskal–Wallis test followed by Dunn’s multiple comparisons test) (**Fig. 1C**). Similarly, the potency of heterologous neutralization was not different for V2, V3, CD4bs, FP and SF bNAbs with median geometric mean reciprocal plasma ID_50_ titers of 102, 92, 152, 139 and 77, respectively (p > 0.05 by Kruskal–Wallis test followed by Dunn’s multiple comparisons test) (**Fig. 1D**). There was also no overall difference in the time to the first detection of heterologous neutralization among the different bNAb specificities (p > 0.05 by Kruskal–Wallis test followed by Dunn’s multiple comparisons test) (**Fig. 1E**), although V2 apex bNAbs developed significantly faster in RMs infected by SHIVs Q23.17 and CH505 compared to all other monkeys (mean of 42 weeks and 44 weeks vs 99 weeks, p = 0.02 and p = 0.01 by Kruskal–Wallis test followed by Dunn’s multiple comparisons test) (**Fig. 1F**). Altogether, these findings indicated that certain Envs exhibited an enhanced predilection for inducing V2 apex bNAbs, with Envs Q23.17 and CH505 standing out for their ability to quickly and reproducibly elicit these antibodies.

### B cell lineage tracing identifies 12 new V2 apex bNAb UCAs with shared features

Essential features of mature human and rhesus V2 apex bNAbs include their long, negatively charged CDRH3s and unique CDRH3 topologies, which enable the antibodies to penetrate Env’s dense apical glycan canopy and engage positively charged C-strand residues (*27, 28, 31, 42–45*). Key questions in the field of germline-targeted immunogen design are how closely do V2 apex bNAb UCAs resemble their affinity-matured bNAb progeny, what distinguishes authentic bNAb UCAs from similarly appearing antibodies that lack the potential to affinity mature to acquire breadth and potency, and what design features must immunogens possess to selectively prime authentic bNAb germline precursors? For humans, only two authentic V2 apex bNAb UCAs have been inferred (*23, 46*), and for rhesus, none have been identified. Thus, to elucidate key features common to a much larger number of such precursors, we lineage-traced the maturation pathways of 12 rhesus V2 apex bNAbs. Previously, Roark and colleagues (*27, 28*) isolated mature bNAb mAbs from these monkeys and determined their immunogenetic and structural features (**Table S2)**. Here we used next generation sequencing (NGS) of IgG+ memory B cells sorted from multiple timepoints beginning soon after SHIV infection through the time of bNAb isolation to sequence immunoglobulin V(D)J gene mRNA and identify bNAb lineage members (**Fig. 2A**). Lineage members were identified based on similarity to mature bNAb heavy and light chain sequences (*28*), and their UCAs and lineage intermediates were determined by phylogenetic inference. This was done using a combination of SONAR (*47*) and IgDiscover (*48, 49*) analyses, which allowed for the assignment of V(D)J alleles not present in published rhesus immunoglobulin gene databases. UCA heavy and light chains were inferred independently and subsequently paired to generate a complete antibody sequence. Importantly, this B cell lineage tracing strategy enabled us to infer germline UCA sequences, including their non-templated residues, with high confidence (**fig. S2A**). Altogether, from ten RMs, we determined 5,461,808 unique IgG+ memory BCR sequences and 651,218 unique IgM+/IgD+ naïve BCR sequences and inferred 12 high-confidence bNAb UCAs (**Fig. 2A and fig S2A**). In addition, we reanalyzed previously reported longitudinal B cell NGS datasets for human participants PC64 (*46*) and CAP256 (*23*) to re-infer their UCAs. While our inference of the PCT64 UCA was identical to that previously published, our inference of the CAP256-VRC26 UCA differed from the previously published UCA by two amino acids in the CDRH3 D-gene templated region (**fig. S2B**).

**Fig. 2.**
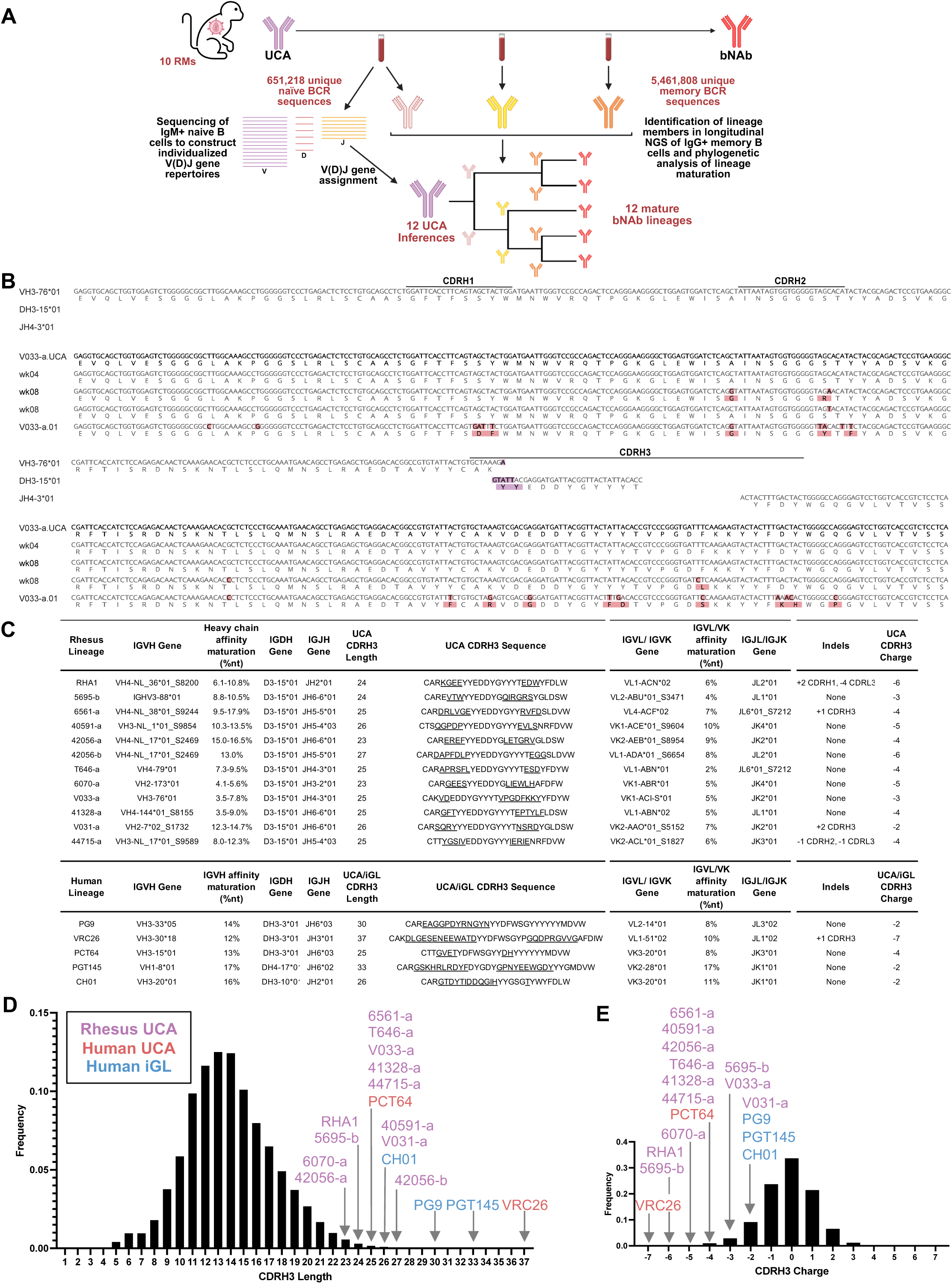
B lineage tracing identifies 12 rhesus V2 apex UCAs. (**A**) Overview of V2 apex bNAb lineage tracing. (**B**) Nucleotide and translation alignment of the V033-a.UCA heavy chain with assigned VDJ genes and B cell NGS-derived lineage members from weeks 4 and 8. Mismatches to the UCA are highlighted. (**C**) Table describing immunogenetic features of rhesus and human V2 apex bNAb UCAs and iGLs. Underlined residues in CDRH3 sequences indicate non-templated residues. Previous estimates of SHM for the rhesus bNAbs reported lower values based on V_H_ gene analysis only (*27, 28*); determinations here included mutations across the entire antibody heavy chain, including the CDRH3, which is under strong positive selection in V2 apex bNAbs. (**D**) Frequency histogram of naïve B cell receptor CDRH3 lengths in the rhesus repertoire. CDRH3 lengths of rhesus and human UCAs and iGLs are indicated. (**E**) Frequency histogram of naïve B cell receptor CDRH3 charge distribution in the rhesus repertoire. CDRH3 charges of rhesus and human UCAs and iGLs are indicated.

Because of frequent monthly sampling following SHIV infection, BCR sequences with very low levels of somatic hypermutation compared with the inferred bNAb UCA could be identified in every monkey (**fig. S3**). An example of the UCA inference for the V033-a lineage is shown in **Figure 2B**. The V033-a lineage was of particular interest because it gave rise to one of the broadest and most potent of all rhesus V2 apex bNAbs, and because it affinity-matured rapidly and with low somatic hypermutation (3.5-7.8% in the heavy chain) (**Figs. 2C** and **3A**) (*28*). In RM V033, a week 4 post-infection memory B cell NGS sequence was identified that exhibited 100% nucleotide sequence identity with the germline monkey-specific V, D, and J genes. Two NGS sequences from memory B cells 8 weeks post-infection bore two and three nucleotide changes, respectively, compared to the week 4 sequence. One of these mutations in the D-J non-templated region was nonsynonymous and corresponded to a positively selected codon where mutations continued to accumulate throughout bNAb maturation. This mutational pattern, together with 199 additional lineage sequences spanning weeks 4-20 (**fig. S4**), lent high confidence to the UCA heavy chain inference depicted in **Figure 2B**. As bNAb lineage members were first detectable at week 4 but were not detectable in B cells sampled prior to SHIV infection, the V033 bNAb lineage must have been primed soon after SHIV infection but before week 4. We could also identify heavy chain sequences from the 5695-b and T646-a lineages that exhibited 100% identity to germline genes in peripheral blood memory B cells from SHIV-CH505 infected RMs 5695 and T646 at weeks 12 and 16 post-infection, respectively (**figs. S2 and S3**). For each of the remaining nine V2 apex bNAb lineages, we identified BCR sequences that had very few heavy and light chain mutations compared with monkey-specific germline alleles (**fig. S3**). In each case, these sequences coalesced phylogenetically to unambiguous heavy and light chain sequences. Thus, for all 12 bNAb lineages we could infer high confidence UCAs.

**Fig. 3.**
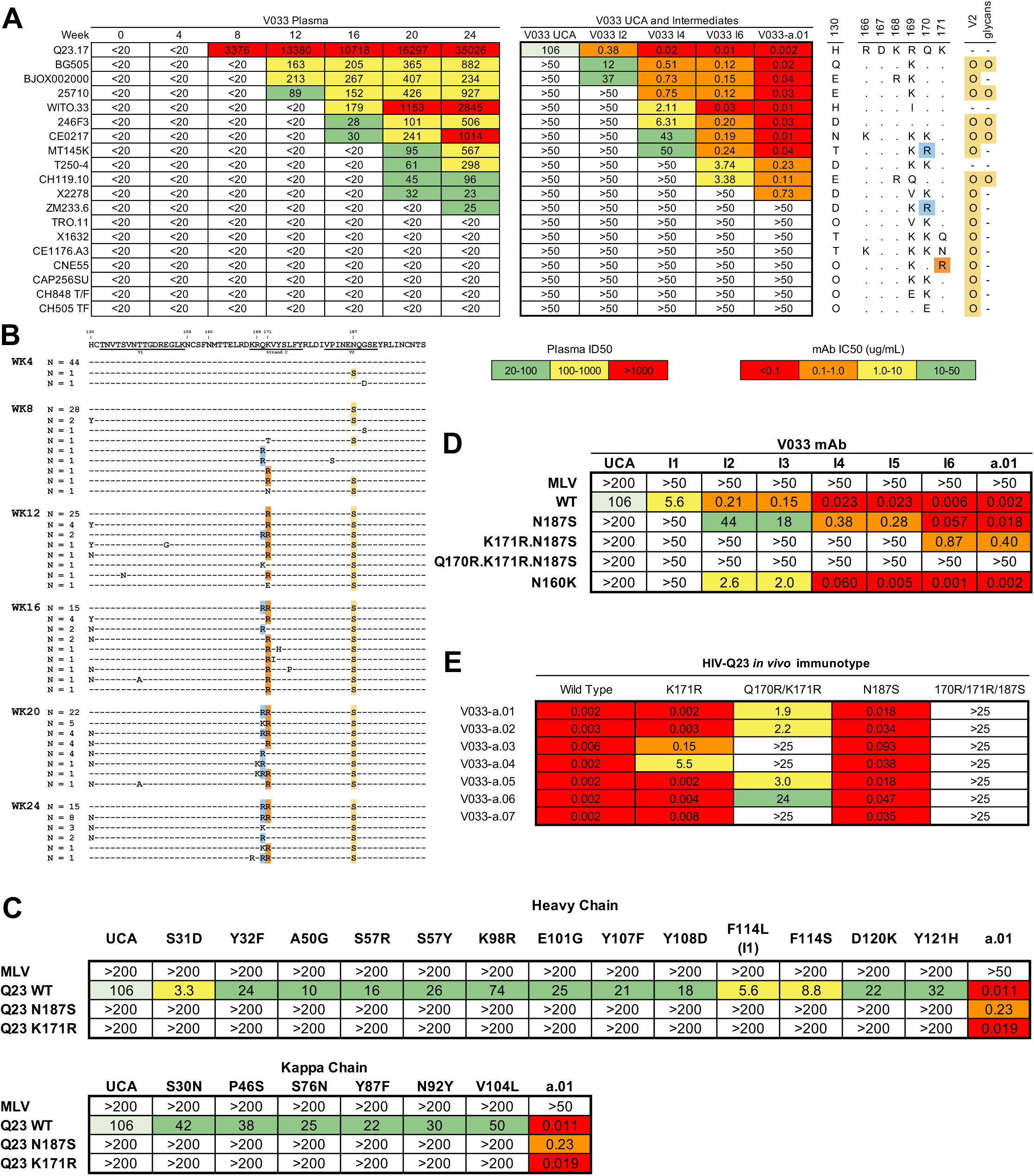
Env-antibody coevolution in RM V033. (**A**) (Left) Longitudinal autologous and heterologous reciprocal plasma neutralization ID_50_ values in SHIV-Q23.17-infected RM V033 adapted from (*28*). (Middle) Neutralization IC_50_ values of V033-a lineage intermediates and UCA against the same panel of heterologous viruses. Increasing intermediate numbers indicate more mature antibodies. (Right) Sequence features of viruses included in the neutralization panel. Residues 130, 166-171, and the number of glycans on the V2 hypervariable loops are shown. Dots indicate sequence identity, and N-linked glycan sequons are denoted by “O.” (**B**) Longitudinal Env V1V2 sequences from RM V033. Sequences are grouped by timepoint indicated on the left. Dashes indicate identity, and colored residues indicate escape mutations from the V033-a lineage. (**C**) Neutralization IC_50_ values of the V033-a.UCA and UCA site-directed mutants containing single mutations from the mature V033-a bNAbs against wildtype Q23.17 Env and its variants. (**D**) Neutralization IC_50_ values of V033-a lineage intermediates against Q23.17 Envs incorporating escape mutations from **B**. (**E**) Neutralization IC_50_ values of mature V033-a bNAb mAbs against Q23.17 Envs incorporating escape mutations from **B**.

Rhesus V2 apex bNAb UCA CDRH3s exclusively utilized the IGHD3-15*01 gene and ranged from 23-27 amino acids in length (**Fig 2C**). This CDRH3 length was slightly shorter than that of five human bNAbs ranging from 25-37, but these differences were not statistically significant (**Fig 2C**). Two of the 12 rhesus V2 apex bNAb lineages, 6561-a and V031-a, contained small CDRH3 insertions of one and two amino acids, respectively (**Fig. 2C**). Therefore, all rhesus lineages acquired their necessarily long CDRH3s during VDJ recombination rather than from insertions during affinity maturation. From these results and others (**Fig. 2C** and B.H.H. and R.A., unpublished), we inferred a minimum CDRH3 length for canonical V2 apex bNAbs to be 22-23 amino acids. This is consistent with findings by Kwong and colleagues who showed that the human mAb 2909, which targets the V2 apex C-strand but can only neutralize N160-glycan deficient viruses, is limited in its neutralization potential because its 21 residue CDRH3 is too short to penetrate the apical glycan shield of most heterologous Envs (*50*). Rhesus bNAb lineages exhibited relatively low levels of SHM, ranging from 3.5% (V033-a.07 and 41328-a.03) to 17.9% (6561-a.15), consistent with shorter maturation pathways compared to bNAbs targeting other epitopes (**Fig. 2C**).

To evaluate how the CDRH3s of rhesus V2 apex bNAbs compared with CDRH3 features in the naïve rhesus and human B cell repertoires, we analyzed ∼650,000 unique IgM+ naïve B cell receptor sequences from 10 RMs that made V2 apex bNAbs (**Fig. 2A**). Naïve B cells were analyzed to minimize bias from clonal expansions and somatic hypermutation following antigen exposure. Rhesus naïve B cell CDRH3s were normally distributed with a mean CDRH3 length of 13.6, a result that is slightly lower than the mean length of 14.8 for human CDRH3s (*51*) (**Fig. 2D**). In our analysis, rhesus B cells with CDRH3 lengths ≥ 23 comprised less than 1% of B cells, while the modal rhesus bNAb UCA CDRH3 length of 25 constituted only 0.16% of the naïve repertoire. Rhesus naïve B cell CDRH3 charge was similarly normally distributed around a net charge of 0 (**Fig. 2E**). Rhesus V2 apex bNAb UCA CDRH3s, however, all had negative charges ranging from −2 (V031-a, found in 9% of B cells) to −6 (RHA1 and 5695-b, found in 0.04% of B cells) (**Fig 2E**). Less than 1% of naïve rhesus B cell CDRH3s exhibited the modal net charge of - 4. Thus, the low frequency of naïve B cells with bNAb features contributes to the infrequency of V2 apex bNAb elicitation in both monkeys and humans. Together, the results indicate that V2 apex bNAb CDRH3s acquire their key features of length and charge during VDJ recombination rather than during affinity maturation, consistent with the interpretation that priming of B cells with these features is a rate-limiting step for V2 apex bNAb elicitation.

### Env-Antibody coevolution in RM V033 leading to neutralization breadth

We next sought to examine steps in bNAb development that followed UCA activation and ultimately led to breadth and potency. We did this first by investigating the V2 apex bNAb lineage V033-a, which exhibited one of the earliest, broadest, and most potent bNAb activities in the cohort. We then extended this analysis to all 12 bNAb lineages to look for generalizable features. RM V033 was infected with SHIV-Q23.17 and rapidly acquired neutralization breadth detectable in plasma as early as 12 weeks post-infection and approximately 10 weeks (range 8– 12 weeks) post-priming of the bNAb lineage UCA (**Fig. 3A**). Neutralization reached 68% breadth on our 18-virus panel with a geometric mean heterologous plasma ID_50_ titer of 1:250 by 24 weeks post-infection (**Fig. 3A**) (*28*). One of the bNAb mAbs isolated from the week 24 time point of V033 exhibited 37% neutralization breadth on an extended 208 virus panel with a geometric mean IC_50_ titer of 0.6 µg/ml (*28*). This rapid acquisition of breadth and potency within a timeframe desirable for a vaccine makes the developmental pathway of the V033-a bNAb lineage an instructive example. We used longitudinal B cell sequencing to infer the maturation pathway of this bNAb lineage, and SGS to characterize coevolving Env escape mutations that selected for affinity maturation and breadth development.

The V033-a.UCA neutralized the infecting SHIV-Q23.17 at an IC_50_ of 106 µg/ml (**Fig. 3A**). Sensitivity of a primary HIV-1 Env to UCA binding and neutralization is unusual, and this finding combined with the very early priming of the V033-a.UCA <4 weeks post-SHIV infection, suggested that the SHIV-Q23.17 Env and not an evolved derivative served as the priming “immunogen.” Analysis of plasma viral RNA sequences from week 4 showed the Env quasispecies to be extremely homogenous (99.85% identical, **Fig. 3B** and **fig. S5A**), supporting the conclusion that the WT Q23.17 Env primed the V033-a lineage. A phylogenetic tree of V033-a heavy chain sequences constructed using IgPhyML (*52, 53*) (**Fig. 4A**) revealed an early bifurcation in bNAb evolution followed by convergent evolution leading to as few as 12 amino acid substitutions in the heavy chain that were critical to neutralization breadth and potency (**Fig 4B**). Intermediates of the sublineage containing V033-a.01, designated V033-a.I1 – V033-a.I6, shared increasing numbers of these substitutions and demonstrated progressive increases in neutralization potency against the infecting SHIV-Q23.17 strain, beginning with an IC_50_ of 106 ug/ml for the UCA and ending at 0.002 ug/ml for the mature V033-a.01 bNAb (**Fig. 3A)**. This 50,000-fold increase in neutralization potency against SHIV-Q23.17 was accompanied by corresponding gains in binding affinity for the bNAb lineage Fabs to the Q23.17 Env trimer (8.47×10^−8^ M for V033-a UCA, 9.33×10^−9^ M for V033-a.I1, and <1×10^−12^ M for V033-a.01) (**fig. S6**). These increases in binding affinity and neutralization potency for the autologous Q23.17 Env were accompanied by corresponding increases in breadth and potency against heterologous viruses, recapitulating the pattern of breadth acquisition detected in the plasma (**Fig. 3A**). For example, just two heterologous viruses in the screening panel (BG505 and BJOX0020000) were neutralized by V033-a.I2 and these two viruses were also neutralized by plasma at week 12 post-infection (**Fig. 3A**). V033-a.I4 neutralized seven heterologous viruses in the panel, and six of these were neutralized by week 16 plasma. Ten viruses in the panel were neutralized by the mature bNAb V033-a.01, and all 10 were neutralized by week 24 plasma. Thus, the full developmental pathway of neutralization breadth in RM V033 plasma was recapitulated by the heavy chain sequences depicted in **Figure S6** and represented in V033-a.I1 – V033-a.I6 and the mature bNAb mAb V033-a.01. Neutralization analysis of a 119-virus panel (*28*) indicated a strong association between resistance to V033-a.01 and presence of the N130 glycan (p < 0.0001, Fisher’s exact test, **fig. S7**). Thus, N130 restriction limited the overall neutralization breadth of the V033-a.01 bNAb lineage.

**Fig. 4.**
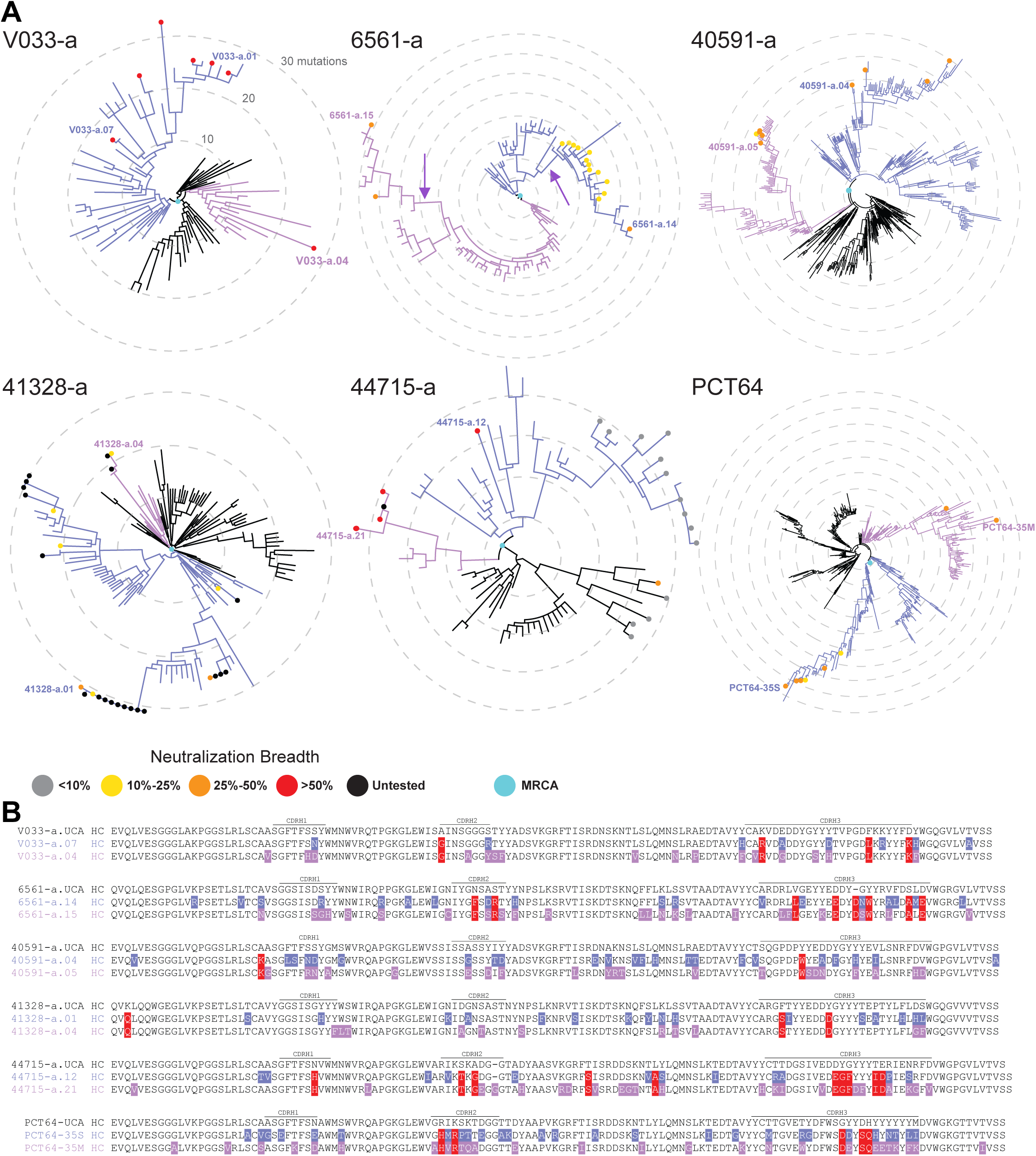
Independent mutational trajectories leading to V2 apex bNAb maturation. (**A**) Radial IgPhyML trees of V2 apex bNAb lineage heavy chains. All trees are rooted on the lineage’s inferred UCA. Grey, yellow, orange, red, and black dots indicate mature bNAb lineage members, colored by their neutralization breadth on our 18-virus panel as previously reported (*28*). Neutralization breadth values for the PCT64 lineage were reported by Landais et al. (*46*). Cyan dots indicate the most recent common ancestor (MRCA) of all broadly neutralizing members of the lineage. Different clades branching off from the MRCA and independently acquiring breadth are colored indigo and lilac. CDRH3 insertions in the 6561-a lineage are indicated by arrows. Scale bars are represented internally, with each concentric circle indicating 10 nucleotide mutations from the UCA. (**B**) Protein alignments of the lineages shown in **A**. UCA heavy chains are aligned with representative mature bNAb lineage members from each independent clade. Mismatches to the UCA are highlighted. Red highlights indicate mutations shared by the two clades.

To elucidate the earliest events in V033 Env-antibody coevolution following priming and how they impacted antibody affinity maturation, we created individual site-directed mutations in the V033 bNAb UCA heavy and light chains corresponding to positively selected mutations observed in memory B cell immunoglobulin sequences and in mature bNAb mAbs (**Fig. 3C and fig. S4**). Each of 19 mutations individually conferred to the bNAb UCA between 1.5- and 20-fold increases in neutralization potency against the autologous SHIV-Q23.17, with an average increase of 7-fold (**Fig. 3C**). The mutation conferring the largest increase in neutralization potency was an F100jL (Kabat numbering) mutation in the CDRH3 D-J non-templated region. This mutation was under strong positive selection as indicated by its high prevalence just 4 weeks after the expanding lineage first became detectable (**fig. S4)**. A structural explanation for this early, strongly selected mutation was provided by a cryoEM structure of the V033-a.I1 antibody in complex with a Q23.17 SOSIP trimer, where modeling showed a progressive accommodation of Env K168 by mutations at F100j from Phe to Leu to Ser (**fig. S8**). The fact that each of the 19 V033 bNAb UCA mutations appeared independently or in unique combinations and each exhibited enhanced neutralization potency against the Q23.17 Env suggests that there was minimal epistasis in the selection of these early mutations. This lack of epistasis and the absence of indels in the bNAb lineage contributed to the permissive maturation pathways that we observed leading rapidly to neutralization breadth. These findings, together with the stepwise >10,000-fold increase in neutralization potency and binding affinity of V033-a.01 bNAb lineage members for the infecting Q23.17 strain (**Fig. 3D**), predicted that the Q23.17 Env as a nonreplicating immunogen might prime and boost the initial steps in B cell affinity maturation of this lineage sufficiently to result in neutralization of homologous and heterologous viruses — a prediction borne out in a V033-a.UCA KI mouse model described in a companion manuscript (*54*).

To examine how SHIV-Q23.17 Env evolution selected for bNAb lineage affinity maturation in RM V033, we performed SGS of gp160 Env sequences spanning 24 weeks of infection (**Fig. 3B** and **fig. S5A**). V033 bNAbs developed maximally in plasma within the first 24 weeks, during which time strongly selected mutations in the replicating plasma virus quasispecies were limited to just a few positions in Env. Some of these mutations (R308H, K460E, D461G/N, N463H, V464E, T533A) corresponded to autologous, strain-specific NAb or cytotoxic T cell escape or fitness reversions (**fig. S5**, and see Supplementary Text) and likely did not contribute to bNAb development. Mutations in the V2 apex C-strand at residues 170 and 171 and in the V2 carboxy-terminus (V2′) at residue 187 (**Fig. 3B**) contributed directly to bNAb affinity maturation by providing a stepwise affinity gradient for evolving bNAb lineage B cells (**Fig. 3E**). While the V033-a.UCA was able to neutralize the wildtype Q23.17 Env, it was unable to neutralize any of the C-strand or V2′ escape variants. Intermediates I1-I6 developed increasing neutralization potency for N187S and N187S + K171R variants, as well as to the N160 glycan KO variant N160K (**Fig. 3D**). Once full plasma breadth developed by week 24 post-infection, the predominant Env variant in the virus quasispecies (Q170R+K171R+N187S) showed complete escape from neutralization by the coevolving bNAbs, presumably limiting their further affinity maturation (**Fig. 3**). These findings suggest a model wherein within-host Env escape mutations selected for more mature V033-a bNAb lineage members that can accommodate those residues in heterologous viruses, a capacity that conferred increasing neutralization breadth as the V033 bNAb lineage matured (see Supplemental Text). In sum, V033-a bNAb ontogeny followed a rapid and simple evolutionary pathway driven solely by the priming Q23.17 Env and three mutations in or near the V2 apex C-strand. Altogether, these findings support the conclusion that priming was the rate limiting step in V033-a bNAb elicitation.

### Common features of Env-antibody coevolution leading to V2 apex bNAbs

In RM V033, bNAb sublineages followed independent evolutionary trajectories leading to the acquisition of breadth. The isolated bNAb mAbs, despite their divergent pathways of development, acquired a limited set of 12-21 convergent amino acid substitutions that were positively selected by affinity for the evolving Env quasispecies (**fig. S4 and Fig. 4B**). To explore the generalizability of these findings to other rhesus V2 apex bNAbs, we constructed phylogenetic trees of eight of the remaining bNAb lineages. We hypothesized that population bottlenecks in phylogenetic trees might reflect potential barriers to affinity maturation and acquisition of neutralization breadth. Conversely, multiple independent maturation trajectories leading to bNAbs without such bottlenecks would suggest less restrictive pathways, similar to that in RM V033 (*55, 56*).

In most monkeys, we found multiple permissive mutational pathways for primed naïve B cells leading to neutralization breadth (**Fig. 4A**). In RMs V033, 6561, 40591, 41328, and 44715 as in the human subject PC64, we found mature bNAb mAbs distributed throughout the respective heavy chain sequence trees. In each case, the most recent common ancestor (MRCA) to the broadly neutralizing mAbs was located close to the UCA, indicating divergence of the lineages into two or more distinct sublineages shortly after priming with each leading successfully to neutralization breadth (**Fig. 4A**). Antibodies from the distinct clades acquired positionally convergent but often chemically distinct mutations that resulted in neutralization breadth (**Fig. 4B)**. This finding suggested multiple permissive diverging pathways to breadth, indicating that the majority of V2 apex bNAb UCAs were able to evolve diverse solutions to epitope recognition (**Fig. 4A)**.

Trees with late MRCAs were indicative of phylogenetic bottlenecks that immediately preceded the development of bNAbs (*21, 57*). These were observed for two lineages: V031-a and 42056-a (**fig. S9A**). The bottleneck in the V031-a lineage coincided with the acquisition of a two-amino acid CDRH3 insertion immediately preceding the acquisition of breadth in the plasma, suggesting that this insertion was a key event in the maturation of this lineage (**fig. S9A**). We deleted this insertion in V031-a bNAb mAbs and found that neutralization breadth was eliminated (**fig. S10**). A very similar result was reported for a CD4bs targeted bNAb lineage in another SHIV-infected monkey (*30*). The other V2 apex bNAb lineage that acquired a CDRH3 insertion, 6561-a, did so twice independently, suggesting strong positive selection (**Fig. 4A**). Members of the 42056-a lineage that exhibited heterologous Env binding and neutralization breadth also clustered tightly in a small section of the tree, and we found this sublineage to be distinguished by acquisition of a set of three disfavored mutations (**fig. S11**) (*58*). For the 5695-b and RHA1 lineages, an intermediate phenotype was observed, with divergence into multiple clades occurring closer to the base of the tree but not as early as for the lineages described above (**fig. S9B**). From this analysis we conclude that potential bottlenecks to bNAb development did occur in a subset of lineages but B cell evolution readily overcame these obstacles. Thus, for most V2 apex bNAb lineages, once a naïve germline B cell with potential for breadth was primed, evolutionary bottlenecks did not substantially hinder their development.

To gain insight into what Envs triggered V2 apex bNAb UCAs, we examined the reactivity of rhesus bNAb UCAs to autologous and heterologous Envs. As observed with other HIV-1 bNAb UCAs (*25*), the majority of rhesus UCAs failed to neutralize any T/F Env tested. The V033-a and V031-a lineage UCAs were exceptions, showing neutralization of the autologous Env Q23.17 at an IC_50_ of 106µg/ml and 183µg/ml respectively (**fig. S12A**). In these RMs, T/F V1V2 sequences comprised the vast majority of Env sequences detected in the plasma preceding the emergence of the bNAb lineages (**Fig. 3B** and **fig. S13**). In contrast, in most other animals, the emergence of bNAbs occurred at timepoints when Envs bearing T/F V1V2 sequences were undetectable in plasma (**fig. S13**). These findings suggest that in these RMs, the priming event for bNAb lineages necessarily involved exposure to evolved viral variants rather than the original T/F Env. Examples of this included N130 glycan deletion in the CH505 plasma virus quasispecies of RMs T279, T646, 5695, 5593, 6070, and T277 (**fig. S14**) and additional V1V2 changes in other monkeys (**fig. S13**), all occurring prior to V2 apex bNAb induction. Still other examples included the 6070-a and T646-a UCAs that were unable to neutralize any primary Env but were able to neutralize multiple N160-glycan knockout viruses (**fig. S12B**). Interestingly, N160-glycan deficient Envs were detectable in the plasma of 6070 and T646 monkeys prior to the triggering of the respective bNAb lineages (**figs. S15 and S16**). These findings lead to an emerging theme in this study and others (*21, 30, 46, 59–63*) that evolution of viral variants, including those with glycan deletions near canonical bNAb epitope supersites, is associated with bNAb lineage priming.

We next asked what Env mutations were responsible for selection of V2 apex bNAb lineage breadth following priming. Longitudinal SGS of the circulating plasma viral quasispecies in the 18 monkeys with V2 apex bNAbs demonstrated strong positive selection at residues 166-171 (**Fig. 5 and fig. S17**). Env residues 166-171, which comprise the N-terminus of the C-strand and two residues immediately preceding it (**Fig 5A**), constitute critical peptide contacts for all canonical V2 apex bNAbs thus far described (*27, 28, 31, 42–45, 64–66*). For brevity, we include residues 166 and 167 together with 168-171 when referring to the C-strand. In 16/18 RMs with V2 apex bNAbs, selection for complete (or near complete) escape occurred at the C-strand (**Fig. 5B and fig. S17**). In the other two monkeys, RMs 42056 and 6729, C-strand replacement reached 70-80%. These findings add substantially to data from two human trial participants with V2 apex bNAbs (PC64 and CAP256) (**fig. S18**) (*23, 46, 61*). Interestingly, distinct patterns of C-strand selection corresponded to each of the three canonical structural motifs found in rhesus and human bNAbs (*28*). This included axe-like bNAb lineages V033-a and 41328-a that selected for mutations primarily at residues 169-171; needle-like lineages 6070-a, T646-a, RHA1, 5695-b, 42056-a, 44715-a and PCT64 that selected for mutations primarily at residues 166-169; and bNAbs with combined modes of recognition — V031-a, 40591-a and CAP256-VRC26 — that showed selection throughout the 166-171 span (**fig. S19, table S2**). Thus, Env escape patterns, which are selected during bNAb lineage affinity maturation, reflected structural and chemical recognition patterns shared by rhesus and human V2 apex bNAb paratopes.

**Fig. 5.**
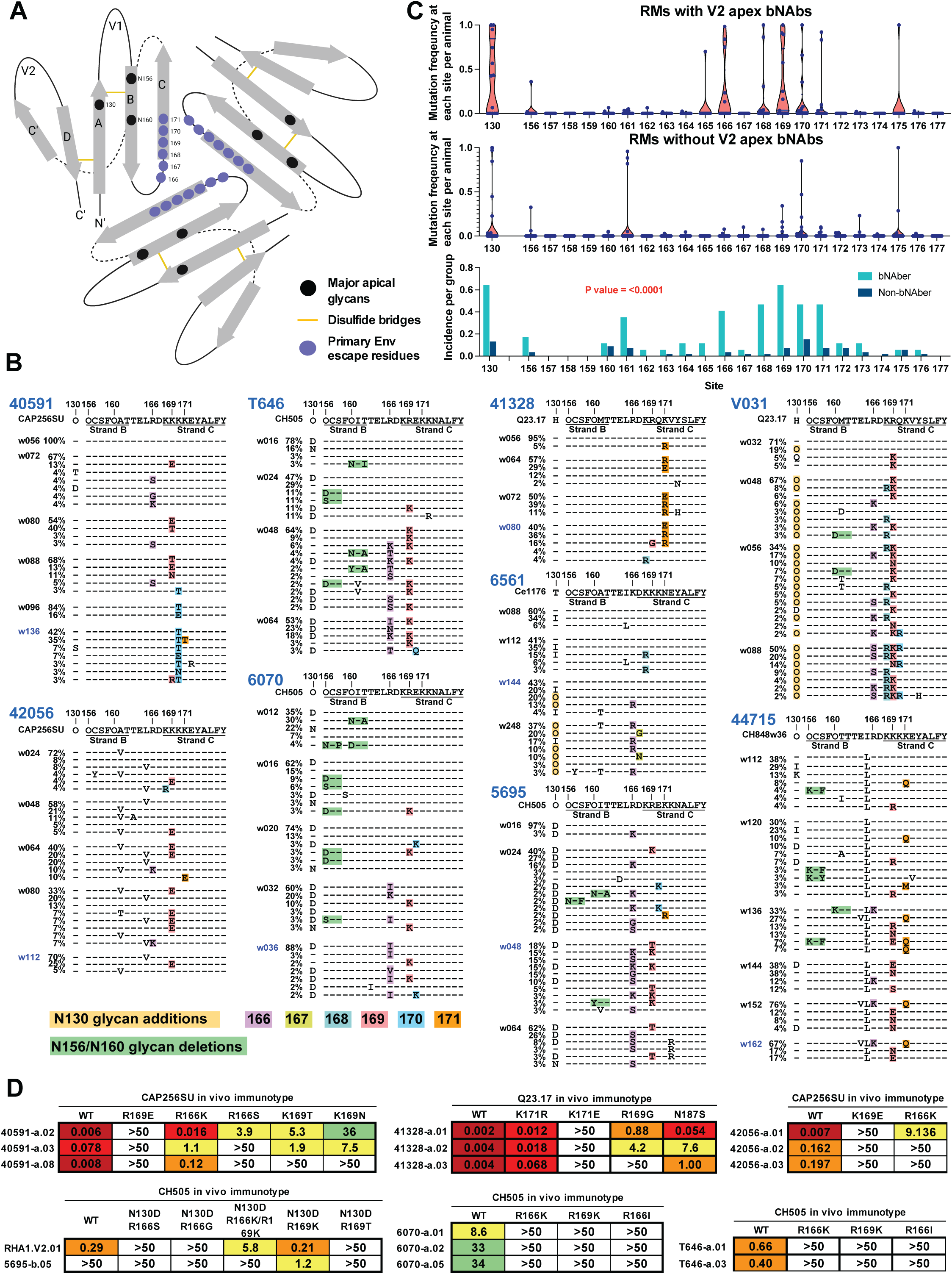
Limited convergent Env mutations guide V2 apex bNAb maturation. **(A)** Protein topology diagram of the Env V2 apex, highlighting positions of key escape residues and glycans. **(B)** Amino acid alignments of longitudinal Env sequences obtained by SGS of circulating plasma virion RNA in RMs with V2 apex bNAbs. Sequences show residues 130 and 156-177 (HXB2 numbering). Sequences are grouped by timepoint indicated on the left in weeks. Dashes indicate identity to the transmitted/founder sequence. Mutations at residues 166-171, mutations that delete glycans at N156 or N160, or mutations that introduce a glycan to residue 130 are highlighted. Glycan sequons are denoted by “O.” Highlighter plots for RMs 5695, 6070, 40591, and 42056 are adapted from Roark et al. (*27*). **(C)** Mutation frequencies of RMs with and without V2 apex bNAbs at residues 130 and 156-177 (top and middle panels) and comparison of mutation frequencies between the two groups (bottom panel; Wilcoxon matched-paired signed rank test, p < 0.0001) **(D)** Neutralization IC_50_ values of V2 apex bNAb lineage members against autologous Env mutants incorporating mutations identified in **B**.

To confirm phenotypically that the C-strand selection observed was responsible for virus escape from V2 apex bNAbs, we constructed site-directed mutants in homologous and heterologous Envs bearing these SGS-identified mutations and assessed their sensitivity to neutralization. All mutations tested resulted in diminished neutralization potency by homologous mature V2 apex bNAb lineage members (**Figs. 3C, S5B**) (*27*). These same mutations in heterologous viruses also reduced or abrogated neutralization by corresponding plasma samples (**fig. S1B**) (*27, 28*). In two RMs, escape also mapped to residues in the V2′ hypervariable loop. For example, in RM V033, partial escape from bNAb neutralization was conferred by mutations at either 170/171 or 187, while complete escape resulted from a combination of all three mutations (**Fig. 3C**). In RM 6561, in addition to C-strand selection (**Fig 5**), escape mapped to residue 190 (**fig. S20A**), where a K190E substitution disrupted a salt bridge between the bNAb CDRH3 and the Env V2′ loop (**fig. S20B**). These results thus indicate that few mutations in a six residue motif in or near the C-strand, and less commonly in the carboxyterminus of V2, are sufficient to allow for virus escape from autologous neutralization and select for bNAb lineage affinity maturation and acquisition of neutralization breadth and potency. An important implication of these findings for HIV vaccine development is that a similarly limited set of minimally divergent prime and boost immunogens may be sufficient to guide V2 apex bNAb maturation.

### C-strand selection is a sensitive and specific indicator of V2 apex bNAb elicitation

To further elucidate the underlying immunobiology of V2 apex bNAb elicitation, we explored the timing of first detection of bNAb lineage members by B cell NGS, first detection of neutralization breadth in plasma, and first detection of epitope-specific Env escape in the evolving plasma viral quasispecies. The question we sought to answer was this: In monkeys that fail to develop neutralization breadth, is this due to a failure to prime relevant naïve germline B cell precursors or a failure to affinity-mature such cells once primed? In other words, does SHIV infection frequently prime V2 apex bNAb B cell precursors that then go “off-track,” or instead, are precursors simply not primed? To address this question, we asked if C-strand selection might serve as a sensitive and specific indicator of C-strand targeted NAbs. Because plasma virus has a lifespan of about an hour, and the cells producing >99% of it, a lifespan of a day (*67–70*), the quasispecies composition of plasma virus serves as an extremely sensitive and dynamic indicator of antiviral selection pressures, including antiretroviral drugs, cytotoxic T cells, or NAbs (*71–76*). We showed previously that epitope-specific mutations in the evolving virus quasispecies can precede the detection of NAbs in the plasma (*30, 64, 72, 73*). Having established C-strand selection at residues 166-171 to be a signature of V2 apex bNAb activity (**Fig. 5**), we calculated Hamming distances of residues 166-171 compared with the infecting SHIV strain in sequential Env gp140 sequences in plasma samples from 101 RMs. In monkeys that developed bNAbs, we plotted longitudinal C-strand Hamming distances against the time of first detection of bNAbs in plasma by the TZM-bl neutralization assay and first detection of bNAb lineage members in circulating blood memory B cells by NGS (**Fig. 6A**). In all ten RMs and two human participants (CAP256 and PC64) with V2 apex bNAbs, a consistent temporal pattern was observed in which detection of bNAb lineage B cells by NGS occurred first, C-strand selection in plasma viral RNA was detected second, followed by detection of bNAb activity in plasma by the TZM assay third (**Fig. 6A**). In only one RM (RM6561) did bNAb activity precede C-strand selection, and in that monkey, we found the early neutralization breadth to be due to a coincident FP-targeted bNAb (**Fig. 1A**, Ce1176). Altogether, the findings confirmed that C-strand selection is an extremely sensitive indicator of V2 apex C-strand targeted bNAbs and underscored the consistency and rapidity with which C-strand selection occurs following V2 apex bNAb lineage priming.

**Fig. 6.**
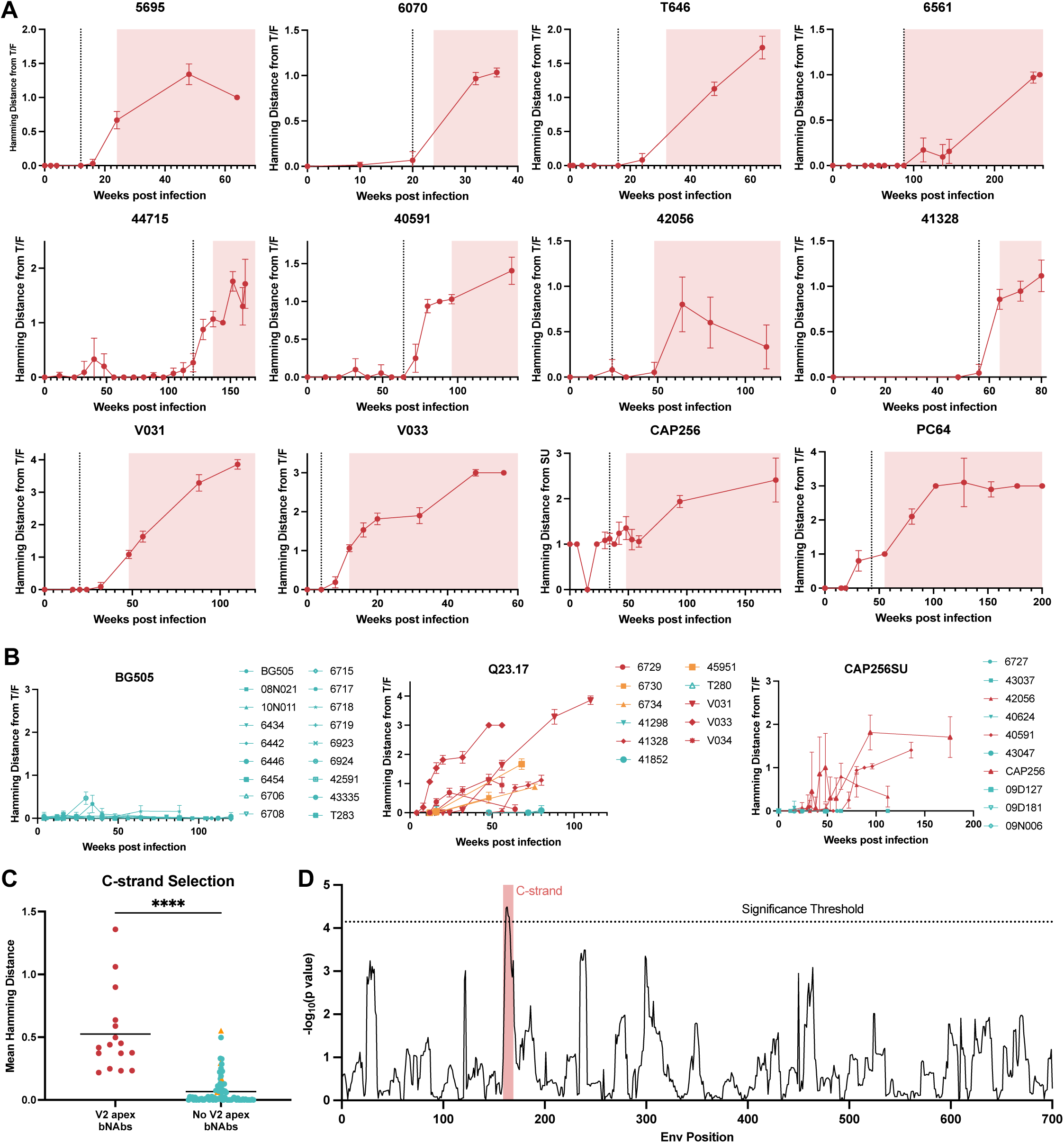
C-strand selection is detectable soon after a V2 apex lineage is triggered but rarely occurs in RMs without V2 apex bNAbs. (**A**) Mean longitudinal Hamming distances of Env residues 166-171 from the transmitted/founder sequence (red lines) increase rapidly in RMs with V2 apex bNAbs following V2 apex bNAb lineage priming as detected by NGS (vertical black dotted lines). Shaded regions indicate timepoints where heterologous tier-2 plasma neutralization was detected by TZM assay. Values for participant CAP256 indicate the distance from the superinfecting strain CAP256SU. (**B**) Mean longitudinal Hamming distances of Env residues 166-171 from the T/F sequence in 101 SHIV-infected rhesus macaques depicted here and in **fig. S21**. Data from human trial participants from which SHIV Envs were isolated are included where available. (**C**) Comparison of the mean C-strand Hamming distances between RMs with and without V2 apex bNAbs. Significance was determined by Welch’s t-test. (**D**) Test for significantly different mean longitudinal hamming distances for a 10 amino acid sliding window across the envelope gene. P values shown are from two-tailed Welch’s t-tests. The significance threshold was adjusted using a Bonferroni correction for multiple comparisons. For all panels, RMs with V2 apex bNAbs are colored red while those without are colored teal. RMs with C-strand targeted antibody responses that did not meet our criteria for breadth are colored orange. All error bars indicate 95% confidence intervals.

Heterologous plasma neutralization developed quickly (mean of 17 weeks) after bNAb lineage B cells were first detected by NGS (**Fig. 6A)**. This timing was similar to previous data for the PCT64 and VRC26 lineages that took 12 and 14 weeks, respectively, to acquire heterologous plasma neutralization (**Fig. 6A)** (*23, 46*). The maximum delay between a lineage being triggered and breadth becoming detectable in plasma was 28 weeks for the V031-a lineage. In this case, breadth appeared just four weeks after the acquisition of a two amino acid CDRH3 insertion. We could thus explain the long delay in breadth acquisition by the requirement for the insertion, which occurred late in lineage maturation (**figs. S9A and S10**). These results suggest that following the priming of V2 apex bNAb UCAs, antibody lineages rapidly acquire somatic hypermutation, which increases binding affinity to the priming virus Env. This exerts strong selection pressure on the autologous viral quasispecies that leads quickly to mutations in the C-strand, which in turn lead to further antibody affinity maturation and accommodation of these mutations resulting in neutralization breadth. The rapid acquisition of neutralization breadth following priming in all ten RMs again highlights the relative simplicity of V2 apex bNAb maturation pathways leading to breadth and potency.

Having established Env escape at residues 166-171 as a highly sensitive indicator of V2 apex bNAb development (sensitivity = 100%), we next sought to determine its specificity by asking what the frequency of C-strand mutations was in RMs that did not develop V2 apex-directed plasma breadth. We reasoned that if the bottleneck to V2 apex bNAb elicitation is at the priming stage, we would fail to see C-strand selection in the replicating virus quasispecies in monkeys without bNAbs. On the other hand, if priming of bNAb precursors occurred frequently but these cells went “off-track” prior to the development of neutralization breadth, we would expect to see evidence of C-strand selection from autologous NAb activity. From over 10,000 longitudinal Env gp140 sequences from 83 RMs lacking V2 apex bNAbs, we found few C-strand mutations.

SHIVs 191859, 40100, CH694, WITO, CH1012, and BG505, all of which failed to elicit V2 apex bNAbs, showed virtually no mutations at the C-strand (**Fig. 6B and fig. S21**). In monkeys infected by SHIVs that in some animals elicited V2 apex bNAbs — Q23.17, CAP256SU, T250, Ce1176, and CH505 — C-strand selection was mostly absent in the subset of RMs that did not develop V2 apex bNAbs (**Figs. 6B and fig. S21**), with five exceptions: RMs 45951 (SHIV-Q23.17), 6730 (SHIV-Q23.17), 6734 (SHIV-Q23.17), 6697 (SHIV-CH505), and 6701 (SHIV-CH505). These monkeys developed NAb responses with very limited breadth that mapped phenotypically to the C-strand (**table S1** and **fig. S22**). Other exceptions included macaques infected by SHIV-CH848 and SHIV-B41, which did not develop V2 apex targeted neutralization but nonetheless showed selection in the C-strand due to reversion to global consensus and/or gain of fitness (**figs. S23 and S24**). Overall, 23/23 RMs with phenotypically demonstrable C-strand targeted neutralizing antibody responses showed C-strand selection in the viral quasispecies as opposed to 8/72 RMs without such NAbs, indicating a sensitivity of (100%) and a specificity of (90%) (see Supplemental Text).

Finally, to determine if other mutations in Env, besides C-strand selection, might distinguish monkeys with V2 apex bNAbs from those without, we explored three different Env mutation metrics: (i) mutation frequency at sites spanning the non-hypervariable regions of V1V2 (**Fig. 5C**); (ii) a comparison of the mean Hamming distances for canonical bNAb epitope regions (C-strand residues 166-171; V3-glycan residues 324-344; fusion peptide residues 512-525; MPER residues 660-678) (**Fig. 6C and fig. S25**); and (iii) a sliding window analysis of time-adjusted Hamming distance area under the curve across the full-length Env (**Fig. 6D**). Each analysis showed a highly significant association between the presence of V2 apex bNAbs and detection of mutations in the C-strand but not other Env regions. These findings reinforce C-strand mutations in the evolving virus quasispecies as a sensitive and specific indicator of C-strand targeted NAb responses.

## DISCUSSION

The primary goal of this study was to determine why only a small subset of SHIV-infected RMs — and by extension, people living with HIV — develop V2 apex bNAbs and how this information can inform new vaccine design strategies. We established five lines of evidence indicating that the principal bottleneck to V2 apex bNAb elicitation lies at the B cell priming stage. First, we identified Envs that preferentially induced V2 apex bNAbs (**Fig. 1A)**. Most of these Envs had been reported previously to bind preferentially to germline-reverted V2 apex bNAbs (*31–35*), which suggests that the primary mechanism behind these Envs’ enhanced bNAb elicitation rates is their higher likelihood of engaging V2 apex bNAb UCAs. Second, we found that the frequency of V2 apex bNAb induction was disproportionately high in RMs and was exclusively linked to the expression of the rhesus D3-15*01 gene (**Figs. 1A, 2C**). In a separate study, we identified structural and biochemical explanations for this preferential D gene usage, including a germline-encoded EDDYG motif that binds positively charged C-strand residues (*28*). The combination of a favored D gene and HIV-1 Envs with a propensity for binding such bNAb UCAs, likely contributes to the high frequency of V2 apex bNAbs in RMs and again points to priming as the rate-limiting step. Third, inference of rhesus UCAs through B cell lineage tracing showed that, from the moment of VDJ recombination, they shared the necessary features of CDRH3 length, charge, and topology to become mature bNAbs. In fact, several of these UCAs were shown to bind Envs from the infecting virus strain (Q23.17 in RMs V033 and V031) or naturally occurring variants of such Envs (CH505 in RMs T646 and 6070). This suggests that priming of naïve B cells, and not complexities of affinity maturation, is the rate limiting step for bNAb elicitation. Fourth, once primed, V2 apex bNAb precursors evolved rapidly to acquire neutralization breadth and potency, following molecular pathways of affinity maturation guided by only few mutations in the Env C-strand. Affinity maturation generally did not require high levels of somatic hypermutation, indels, or show evidence of epistasis or evolutionary bottlenecks. The average time from UCA priming, as determined by NGS detection of bNAb lineage members in memory B cell mRNA, to acquisition of neutralization breadth in plasma was just 17 weeks. Fifth, C-strand selection in the evolving plasma virus quasispecies was found to be a sensitive and specific indicator of V2 apex C-strand targeted antibody responses. Our failure to find C-strand selection in most RMs that lacked V2 apex bNAbs suggested that such B cell precursors were either not primed, or if they were, then they failed to expand and affinity mature even to the point of producing autologous C-strand targeted NAbs. We conclude from these findings that, in the setting of SHIV/HIV infection, V2 apex bNAb elicitation is uncommon because B cell priming is inefficient.

Why then is V2 apex bNAb priming so inefficient during the course of most SHIV/HIV infections? Successful priming requires a sufficient number of suitable B cell precursors and Env ligands with adequate affinity to bind them. Necessary features of bNAb B cell precursors include, but are not limited to, adequate CDRH3 length (>22 residues), aromaticity, negative charge, and appropriate structural motifs (*28, 35, 64, 66, 77*). Based on such criteria, several studies have estimated the frequency of potential V2 apex bNAb precursors to be in the range of 1 in 10^5^–10^7^ naïve B cells (*64, 66, 77, 78*). But these features alone are unlikely to be the only requirements for B cell precursors to have the potential to affinity mature to neutralization breadth and potency. Thus, the identification of immunogenetic, structural and chemical features that distinguish actual V2 apex bNAb precursors from “bNAb-like” precursors will remain a high priority (*77*). Nonetheless, our findings of special “bNAb imprinting” Envs that exhibit a higher frequency of V2 apex bNAb precursor priming and bNAb elicitation (**Fig. 1**, Q23.17 and CH505), combined with promising results from recent immunization studies using germline-targeted, near-native Env trimers (see below), indicate that authentic V2 apex bNAb precursors circulate at appreciable frequencies. Altogether, the findings suggest a model wherein the critical bottleneck to V2 apex bNAb elicitation — priming — is not due to a paucity of suitable naïve germline precursors, but instead, to a lack of suitable priming Env “immunogens” that exhibit sufficient affinity to activate these precursors. When bNAbs develop later in infection, it may be due to evolution in the circulating virus quasispecies that enables enhanced precursor binding as a result of N130 or N160 glycan deletions or other changes in Env, as occurred in RMs 6070, T646 and other RMs (**figs. S14-S16**). In other words, our data suggest that stochastic processes associated with the transmission of particular Env variants, the evolution of these variants, and the randomness of VDJ recombination with N-nucleotide additions, combine to explain the infrequency of V2 apex bNAb elicitation in SHIV and HIV infection, a conclusion supported by other studies (*79*). A key implication of these findings for new HIV-1 vaccine designs is that if Env immunogens can be engineered to exhibit enhanced binding to multiple V2 apex bNAb UCAs, such precursors are likely present in sufficient numbers to generate a bNAb response in most rhesus and humans.

Efficient priming as a rate-limiting step to bNAb induction may not be limited to the V2 apex site. In **Fig. 1**, we show that CD4bs-targeted bNAbs developed in only one of 115 SHIV-infected RMs. Building on studies by Kwong, Wyatt, and Binley (*80–82*), we designed a novel SHIV.CH505 lacking glycans surrounding the CD4bs and found that it induced CD4bs bNAbs in 9 of 11 macaques within one year of infection, a highly significant enhancement (p<0.0001, Fisher’s Exact Test) (*30*). Similarly, in **Fig. 1** and **Table S1** we show that V3 glycan bNAbs developed in only four of 115 SHIV-infected RMs, and then only after more than a year of infection. By introducing mutations that shortened and reduced glycan occupancy in V1 adjacent to the canonical ^325^GDIR_328_ binding motif of the V3 glycan bNAb supersite in SHIV-BG505.5MUT, we enhanced the frequency of V3 glycan bNAb elicitation to 14 of 22 RMs after just four months of infection, again a highly significant increase (p<0.0001, Fisher’s Exact Test) (*83*). Similar results have been reported for fusion peptide targeted bNAbs (*73*). Altogether, of these findings point to priming as a rate-limiting step to the elicitation of many classes of bNAbs, and thus suggest new directions for immunogen design.

The finding that B cell priming represents a primary obstacle to V2 apex bNAb elicitation during infection has important implications for new HIV-1 vaccine design strategies. A key inference is that if the efficiency of V2 apex bNAb UCA priming can be substantially enhanced, it is likely to lead to more frequent and rapid bNAb elicitation. Recent findings reported in a companion paper (*54*) and elsewhere (*64, 78, 84*) show this to be the case. Ghosh and colleagues found that a stabilized version of the Q23.17 Env trimer (Q23.Apex.GT1) induced bNAbs in KI mice expressing the V033-a.I1 B cell receptor after a single prime and homologous boost (*54*). Andrabi and colleagues took this a step further with the design of Q23.Apex.GT2, which differs from GT1 by only two amino acids in gp120, and showed that this immunogen given as a homologous prime and boost to RMs could induce V2 apex bNAbs in germinal center B cells, circulating Bmem cells and in plasma (*78*). A key feature of both Q23.Apex.GT1 and .GT2 is that they are minimally modified from the wildtype Q23.17 Env trimer, which enables them to bind rhesus and human bNAb UCAs with enhanced affinity, and at the same time, allows affinity maturation to be driven from the outset by a native-like tier-2 HIV-1 Env trimer. This strategy is different from others where native-like Envs are used for late “polishing” steps (*77, 85*). Wyatt and Kulp also explored Q23.17-derived Envs as immunogens, and both groups observed V2 apex bNAb responses in KI Ig-rearranging mice and RMs (*64, 84*). Altogether, the findings suggest that if V2 apex bNAb UCA priming can be enhanced by germline-targeted, native-like Env trimer immunogens, bNAb development can readily follow with boosting strategies that are less complicated than previously envisioned.

With respect to new strategies for vaccine development, the current study provides a molecular guide and a toolkit of reagents and sequences for designing novel prime and boost immunogens. Previously, there were just two inferred human V2 apex bNAb UCAs and no rhesus UCAs (*23, 46*). The current study increases this number by six-fold by reporting 12 high confidence rhesus V2 apex bNAb UCAs (**Fig. 2B, C**). Already, these have proven to be extremely valuable as screening tools for new germline-targeted immunogens (*54, 78*). Similarly, the HIV-1 Envs identified in **Fig. 1A** as showing a propensity for eliciting V2 apex bNAbs have been incorporated into at least five new vaccine platforms all of which have been shown to induce V2 apex bNAbs in rhesus UCA KI mice, human immunoglobulin rearranging KI mice, or RMs (*54, 64, 78, 84*). Env sequences from Env-antibody coevolution analyses have also been used to develop boosting immunogens that have shown success in KI mouse models (*54*). The molecular pathways in Env-antibody coevolution reported here may thus provide enabling datasets to investigate shared patterns in V2 apex bNAb induction that can be translated into new vaccine designs.

Our study has certain limitations. First, it represents a prospective analysis of bNAb elicitation in 122 SHIV-infected monkeys with an average observational period of 2 years (range 0.6 to 6 years), as opposed to a retrospective characterization of bNAb development in highly selected individuals culled from much larger human cohorts where the duration of infection was generally much longer (*4, 6, 86–88*). In the latter case, investigators could choose just those individuals with the greatest neutralization breadth and potency for analysis, whereas in our studies, we included all SHIV-infected RMs from infection onward. One prediction, borne out by the results presented here, was that a prospective analysis of bNAb elicitation in rhesus was likely to identify animals with a far wider range of neutralization breadth and potency than might be considered desirable, yet such diverse outcomes are likely to be representative of the spectrum of antibody responses resulting from even the most promising vaccine candidates. Future vaccine trials will need to consider this possibility. A critical question that remains to be answered is whether the neutralization breadth observed in the 18 monkeys with V2 apex bNAbs represented in **Table S1**, which ranged from 11-94% against our 18-member virus panel, was limited by properties inherent to the particular bNAb UCAs that were primed and the particular escape pathways of the homologous viral quasispecies that selected for their affinity maturation. Or, is it possible that the neutralization breadth potential of these UCAs is much greater and could be realized by boosting with heterologous HIV-1 Envs. This is a question highly relevant to vaccine prime and boost designs and is one that can be addressed in the KI mouse model where individual V2 apex bNAb UCAs can be primed and boosted with homologous and heterologous Env immunogens (*54*).

A second limitation to the study relates to host immunogenetics. The rhesus D3-15*01 gene was expressed by all 12 rhesus V2 apex bNAbs (**Fig. 2C**). This gene is highly favorable for V2 apex bNAb priming and development as it encodes an EDDYG motif that provides negative charge, aromaticity, and potential for tyrosine sulfation, each of which can contribute to C-strand binding (*28*). There is no human homologue of the rhesus IGHD3-15*01 gene, although a similar EDDYG motif can be generated at appreciable frequencies by non-templated nucleotide addition during V-D-J rearrangement (*64, 77*). Moreover, an EDDYG motif can evolve in human D genes under positive selective pressure as exemplified by the PCT64 bNAb lineage (*46*). It remains to be seen whether the priming efficiency of human V2 apex bNAb UCAs can be enhanced as readily as for rhesus and whether affinity maturation of human V2 apex bNAb lineages can be as straightforward as for some rhesus V2 apex bNAbs. Answers may come soon from human clinical trials of promising new germline-targeted V2 apex immunogens that are planned for the near future by the HIV Vaccine Trials Network (protocol HVTN 322) and BRILLIANT Consortium (protocol B-003). In summary, the results presented in this paper indicate that the rhesus model is highly amenable for the iterative design, development and testing of immunogens targeting the V2 apex. If vaccination regimens can be developed in rhesus that consistently elicit V2 apex targeted bNAbs at titers that are clinically protective against heterologous tier 2 SHIV challenge, it would represent an important proof-of-concept advance in the pursuit of an effective HIV-1 vaccine for humans.

## MATERIALS AND METHODS

### Nonhuman primates

Indian rhesus macaques ages 3-12 years, evenly split male and female, were housed at Bioqual Inc. in accordance with the Association for Assessment and Accreditation of Laboratory Animal Care (AAALAC) guidelines. The study design and all experimental procedures were approved by the University of Pennsylvania and Bioqual Institutional Animal Care and Use Committees. Monkeys were naïve with respect to prior SHIV infections or inoculations. Eight monkeys were repurposed from prior immunization studies with HIV-1 gp120 monomeric protein or SOSIP Env trimer proteins. None of these animals had HIV-1 nAbs, bNAbs or bNAb lineage precursors detectable at the time of SHIV infection and study initiation. RMs were sedated for procedures including blood draws, anti-CD8 mAb infusions, and SHIV inoculations. A subset of animals received intravenous or subcutaneous infusions of 25 to 50 mg/kg of anti-CD8α mAb (MT807R1) or anti-CD8β mAb (CD8β255R1), administered at the time of SHIV inoculation or two or seven days prior. These anti-CD8 mAbs transiently depleted CD8 cells for a duration of approximately two months, resulting in elevated peak and set point viral loads compared with monkeys that did not receive anti-CD8 mAbs. Details concerning monkey demographics, study design, and clinical, immunological and virological outcomes are provided in **Table S1**.

### Processing and storage of specimens

RM blood samples were collected in sterile vacutainers containing the anticoagulant acid citrate dextrose solution A (ACD-A). A total of 40mL of ACD-A-anticoagulated blood was transferred into a sterile 50 mL conical tube and centrifuged at 1000×g for 10 minutes at 20°C. Plasma was carefully collected into a new 50 mL conical tube so as to not disturb the buffy coat and red blood cell pellet. To remove residual platelets and cells, the plasma was centrifuged again at 1500×g for 15 minutes at 20°C. The resulting plasma was aliquoted into 1mL cryovials and stored at –80°C. The remaining pellet was resuspended in an equal volume of Hanks’ Balanced Salt Solution (HBSS) without Ca² or Mg² and supplemented with 2mM EDTA. The mixture was divided into four 50 mL conical tubes, and additional HBSS-EDTA was added to bring each tube to a final volume of 30mL. Each cell suspension was then gently underlaid with 14mL of 96% Ficoll-Paque and centrifuged at 725×g for 15 minutes at 20°C with slow acceleration and deceleration to maintain the Ficoll-cell interface. Mononuclear cells at the interface were collected and suspended in HBSS-EDTA, followed by centrifugation at ∼500×g for 15 minutes at 20°C. The pellet was resuspended in HBSS containing Ca² and Mg² and supplemented with 1% fetal bovine serum (FBS). To remove additional platelet contamination, the suspension was centrifuged at 200×g for 15 minutes at 20°C, and the supernatant discarded. The resulting pellet was resuspended in 0.1-0.3mL of residual media and brought to 25mL with HBSS + 1% FBS. Cell counts and viability were determined using acridine orange and propidium iodide staining. Cells were then centrifuged at 300×g for 10 minutes at 20°C, resuspended at 5-10×10 cells/mL in CryoStor cell cryopreservation medium, and aliquoted into 1mL cryovials. Cryovials containing PBMCs were stored in Corning CoolCell LX freezing containers at –80°C overnight, followed by transfer to vapor phase liquid nitrogen for long-term storage.

### Plasma vRNA quantification

Plasma viral load measurements were performed by the NIH/NIAID-sponsored Nonhuman Primate Virology Core Laboratory at the Duke Human Vaccine Institute. This core facility is CLIA certified and operates highly standardized, quality-controlled Applied Biosystems Real-Time SIV and HIV vRNA PCR assays. QIAsymphony SP and QIAgility automated platforms (QIAGEN) were used for high throughput sample processing and PCR setup. Viral RNA was extracted and purified from plasma, annealed to a target specific primer and reverse transcribed into cDNA. The cDNA was treated with RNase and added to a custom real-time PCR master mix containing target specific primers and a fluorescently labeled hydrolysis probe. Thermal cycling was performed on a QuantStudio3 (ThermoFisher Scientific) real-time quantitative PCR (qPCR) instrument. Viral RNA cp/reaction was interpolated using quantification cycle data. Raw data were quality-controlled, positive and negative controls checked, and the mean viral RNA cp/ml calculated. The lower limit of quantification (LLOQ) of this assay ranged from 62 - 250 RNA copies/mL over the extended course of this study.

### Virus stock generation

SHIV and pseudovirus stocks were generated by transfection of HEK 293T/17 cells (ATCC, CRL-11268). 4-5 × 10^6^ cells were plated in 100mm dishes and grown in DMEM containing 10% FBS and 1% Penicillin/Streptomycin overnight at 37°C. SHIV or HIV-Env plus Env-minus proviral plasmid DNA was then transfected using FuGENE 6 transfection reagent (Promega, Cat # E2691) according to the manufacturer’s instructions. Transfected cells were then incubuated for 48h at 37°C. Supernatants were then centrifuged, aliquoted, and stored at −80°C. Site-directed mutants were generated using the Q5 Site-Directed Mutagenesis kit (New England BioLabs) according to the manufacturer’s instructions and produced as described above.

### Virus titration

Titration of 293T cell virus stocks were performed using TZM-bl cells. TZM-bl cells were grown in cell culture medium (DMEM supplemented with 10% FBS, 1% penicillin/streptomycin) and were plated in 96 well plates at a concentration of 1.5×10^4^ cells/well and incubated at 37°C overnight. Viruses were diluted five-fold in cell culture media supplemented with 40 ug/ml DEAE-dextran and incubated in quadruplicate with TZM-bl cells at 37° C for 48 hours. Infected cells were then fixed for 10 minutes at room temperature in PBS containing 0.8% glutaraldehyde and 2.2% formaldehyde, washed 3 times with PBS, and stained with PBS containing 4µM Magnesium Chloride, 4µM Potassium Ferricyanide, 4µM Potassium Ferrocyanide, and 400µg/ml X-Gal for 3 hours at 37°C. Stained cells were then washed 3 times with PBS, and imaged on a CTL analyzer (Immunospot). The number of spot counts per virus dilution were then averaged and divided by the volume of viral input to calculate the concentration of infectious viral units.

### Neutralization assays

Neutralizing antibody assays were performed using rhesus plasma and TZM-bl indicator cells, as previously described (*27*). Briefly, TZM-bl cells were grown in cell culture medium (DMEM supplemented with 10% FBS and 1% penicillin/streptomycin), were plated in 96 well plates at a concentration of 1.5×10^4^ cells/well and incubated at 37°C overnight. Plasma samples were heat inactivated at 56°C for 1 hour, clarified by brief (30 sec) microfugation, and serially diluted (5-fold) starting at a dilution of 1:20. Normal heat-inactivated human or rhesus plasma was used to adjust final plasma concentrations (test plus normal) in all wells to 5% in addition to 10% FBS. For monoclonal antibody neutralization assays, mAbs were serially diluted starting at a concentration of 50µg/ml for mature bNAbs and 200µg/ml for UCAs in cell culture media lacking normal rhesus or human serum. Viruses were diluted to achieve a multiplicity of infection (MOI) of 0.3 upon addition to TZM-bl cells. Viruses were incubated with serum or antibody dilutions at 37°C for one hour, after which the virus-serum or virus-antibody mixtures were plated onto adherent TZM-bl cells and incubated at 37°C for 48 hours. Care was taken to ensure that the total concentrations FBS and normal human or rhesus plasma were held constant across all wells. Following incubation, cells were lysed with 0.5% Triton-X 100 in PBS and luciferase activity levels measured using the Promega luciferase assay system (Promega Cat# E1501) on a BioTek Synergy Neo2 plate reader (Agilent Technologies). Neutralization IC_50_/ID_50_ values were calculated using Prism 10 software.

### Antibody production

To produce antibodies, paired heavy (VDJ) and light (VJ) chain variable gene sequences were commercially synthesized (GenScript) and cloned into antibody expression plasmids as previously described (*27*). Recombinant antibodies were produced by cotransfecting paired heavy and light chain expression plasmids into Expi293F cells using ExpiFectamine 293 transfection reagents (Gibco), purified from culture supernatants using the Protein A/Protein G GraviTrap kit, and buffer-exchanged into PBS as previously described (*27*).

### BLI affinity measurements

BLI was performed to determine the affinities of V033-a lineage members for Q23 trimers. Fabs (10 µg/ml) were immobilized on Protein A sensors (Sartorius) to a signal of 1.0 nm using an Octet Red96 instrument (ForteBio). The immobilized Fabs were then dipped in running buffer (PBS containing 0.1% bovine serum albumin and 0.02% Tween-20 at pH 7.4). Sensors were then dipped into wells containing Q23 trimers serially diluted in running buffer. Four two-fold dilutions were tested for each trimer, starting at a concentration of 500nM. Following a 120s association period, the tips were dipped into the running buffer and dissociation was measured for 240s.

### B cell next-generation sequencing

B cell NGS was performed as previously described (*27, 89*). Briefly, PBMCs were stained with LIVE/DEAD Aqua, CD3-PerCP-Cy55, CD4-BV785, CD8-BV711, CD14-PE-Cy7, CD20-BV605, IgD-FITC, IgG-AF680, and IgM-BV650. Memory B cells (CD20+, IgG+, IgD-, IgM-) were bulk sorted into RPMI with 10% FBS and 1% Pen-Strep using a BD FACSAria II. RNA was extracted using RNAzol RT according to the manufacturer’s guidelines (Molecular Research Center, Inc). cDNA was synthesized using a 5’ rapid amplification of cDNA ends (5’ RACE) approach, with SMARTer cDNA template switching and Superscript II RT. Following synthesis, cDNA was purified using AMPure XP beads (Beckman Coulter). Immunoglobulin transcripts were then selectively PCR amplified using IgG, IgK, or IgL constant region-specific primers and KAPA HiFI HotStart ReadyMix (Roche). Finally, an additional PCR step was used to append Illumina P5 and P7 sequencing adaptors to the libraries. Both heavy and lambda immunoglobulin libraries were sequenced on an Illumina Miseq sequencer with 2 × 300 bp runs using the MiSeq Reagent V3 kit (600-cycle).

### Antibody Lineage Tracing Workflow Details

To delineate the ontogeny of each antibody lineage, we analyzed antibody mRNA transcripts from an unbiased sampling of each macaque’s B cell receptor repertoire, both for naive antibodies (from IgM+/IgD+ B cells) and mature antibodies (from IgG+ B cells). We used a common amplification and sequencing approach and common early data processing steps, followed by more specific workflows for the two main aspects of the analysis corresponding to the naive and mature cells. Sequencing of the naive (IgM+/IgD+) B cells allowed us to indirectly infer the unrearranged, genomic antibody germline gene repertoire for each macaque, primarily using the software IgDiscover (*48, 49*) and MINING-D (*90*). We applied a separate workflow to antibody sequences from IgG+ B cells, using the program SONAR (*47*) to identify sequences belonging to our lineages of interest across longitudinal sampling and to construct phylogenies leading back to the inferred unmutated common ancestor (UCA) of each antibody lineage. The germline sequences derived from the naive repertoire dataset informed this analysis by supplying confident germline gene assignments for each lineage. In addition to the specific programs and techniques described in more detail below, we used custom code written in Python and Snakemake to coordinate the overall data processing and workflow organization. A final section provides software versions and URLs including for this custom code.

#### Sequencing and initial data processing for all samples

Library preparation and sequencing was as previously described (*28, 89*). In short, we used 3’ primers specific to the applicable rhesus macaque antibody constant regions and a universal 5’ RACE primer to amplify the beginning of the antibody transcript including the leader through the J segment to the beginning of the constant region. A second round of PCR incorporated Illumina sequencing adapters and custom sample barcoding. An “inline” barcode was placed at the start of the forward read (R1), including a varying-length randomized nucleotide prefix for higher nucleotide diversity during sequencing. A second barcode was placed in an index read (I1). To handle this custom barcoding strategy, we performed demultiplexing as part of the data processing after sequencing, using the igseq demux command in our Python program (described below), rather than with default Illumina software. The read layout was 309×8x309 NT for R1, I1, and R2, with the overlapping R1 and R2 sequences merged during processing following adapter trimming. The output from this step of our workflow was the trimmed and merged antibody reads with adapters and primers removed, spanning from the beginning of each antibody transcript to the end of the J segment.

#### Establishing individualized antibody germline repertoires

As previously described (*28*) we used the IgM+/IgD+ repertoire sequencing reads (demultiplexed, trimmed, and merged as described above) to develop individualized macaque-specific V and J segment germline references, as well as to check the presence of germline D gene sequences. We used IgDiscover to infer personalized germline sequence sets from a set of generic rhesus macaque starting databases. For the IGH locus, the starting database was KIMDB 1.1 (*91*), a recently published database of rhesus and cynomolgus macaque IGHV, IGHJ, and IGHD sequences. For IGK and IGL, we used an earlier published germline reference (*92*), as KIMDB does not currently include light chain sequences. In certain instances, we observed large numbers of spurious novel J sequences in IgDiscover’s output, particularly for the IGKJ2 family, but at times for heavy chain sequences as well. To exclude these from the final output we applied an additional filtering step for J sequences using IgDiscover’s “discoverjd” feature. As IgDiscover reports evidence of expression of known germline D sequences but does not infer new alleles, we also used the program MINING-D to infer possible D sequences and check for any signs of relevant novel D segments in our lineages of interest. We first annotated, filtered, and clustered the IgM+/IgD+ sequences using the steps described below for lineage tracing for IgG+ sequences, in order to get consensus CDRH3 sequences for each naive antibody repertoire, and gave these sequences as input to MINING-D. We used two p value thresholds (4.5×10^−36^, labeled “default” in the workflow, and 4.5×10^−16^, labeled “sensitive”) to produce two levels of sensitivity in the output for each macaque. We compared MINING-D’s output sequences with IgDiscover’s expression tables and with KIMDB’s rhesus macaque reference D sequences, as well as other known macaque antibody germline references. As MINING-D is intended as an initial check for possible novel alleles to be confirmed with genomic sequencing, we did not attempt to infer a complete set of germline D sequences per macaque with this approach; instead, we used it to investigate particular established germline D sequences such as those assigned for our lineages of interest. The output from this step was the inferred V and J allele sequences from IgDiscover and, for heavy chains, inferred D allele sequences from MINING-D with two tiers of sensitivity.

#### Lineage Tracing

In parallel to the steps described above for investigating per-macaque germline sequences, we used SONAR to analyze IgG reads from longitudinal samples and trace the development of our specific lineages of interest. SONAR’s operation is organized in three sequential stages: initial sequence clustering and annotation (module 1), lineage member identification (module 2), and phylogenetic analysis (module 3). The germline information inferred for each macaque was used directly in SONAR’s annotation steps and to inform the final germline assignments for each lineage to determine the most accurate unmutated common ancestor (UCA) inference.

#### Lineage Tracing: SONAR Module 1 (Annotation)

The IgG+ repertoire sequencing reads (demultiplexed, trimmed, and merged as described above) were supplied to SONAR’s module 1 scripts, which uses BLAST+ to assign germline gene segments using each macaque’s germline reference and then clustering similar reads (threshold of 99% for clustering and a minimum of 2 identical sequences per cluster). Only productive-looking antibody sequences (those with recognizable junction regions including codons for conserved amino acids, and without frame shifts or stop codons) were included for subsequent analysis. The output from this step was, for each sample, an AIRR-compatible (*93*) TSV table of annotations for clustered antibody sequences and associated FASTA files corresponding to a representative sequence from each cluster.

#### Lineage Tracing: SONAR Module 2 (Lineage Member Selection)

Following annotation and clustering, we used SONAR’s manually-guided identity/divergence feature for lineage member identification in each sample. This step calculates sequence identity to known antibodies for lineages of interest and divergence from assigned germline V sequences (as defined automatically module 1’s annotation and supplied in the AIRR file) for the productive-looking sequence clusters identified in module 1 (SONAR’s “goodVJ unique” sequence set). The identity/divergence file is then supplied to SONAR’s “island” selection script to identify groups (“islands”) of candidate lineage members on a specialized scatterplot of all sequences for each sample. These plots display divergence from germline V on the x axis and identity to one reference antibody on the y axis, with candidates selected by clicking points to define a boundary polygon on each plot. The final set of candidate lineage members is the combination of all selected repertoire sequences across the known representative lineage members. The list of sequence IDs for candidate lineage members is then supplied to SONAR’s getFasta utility script to produce a FASTA file of lineage members for each lineage for each sample.

Our default approach was to use all initial known lineage members as references for the island selection process, with extensive manual review of the candidates afterward to resolve any ambiguous cases (considering evidence such as shared mutations from germline or matching nontemplated nucleotides in the junction region). While relatively labor-intensive compared to automated alternatives, this approach allowed more flexibility in confidently identifying true lineage members than rule-based criteria (e.g., requiring matching germline V and J genes, junction lengths, and a fixed junction similarity threshold) since it can identify sequences with difficult-to-assign germline genes or with indels relative to known lineage members. While not built into our software workflow directly, candidate UCA sequences and new lineage member sequences identified in the repertoire NGS samples can also be supplied as references for additional rounds of identity calculations and island selection through manual addition to the relevant files. We found that the default behavior of SONAR’s island selection script, using kernel density estimation for a smoothed density visualization, could fail to display individual sequences in certain edge cases. We introduced an alternate plotting approach to instead display sequence counts per binned tile of identity and divergence intervals, which ensures all possible sequences are visible on the plot. This option is now available in SONAR’s script as “sonar get_island – plotmethod binned” and was used by default in our workflow. The output from this step was, for each sample and each lineage, a FASTA file with sequences of identified lineage members.

#### Lineage Tracing: SONAR Module 3 (Phylogeny Analysis)

To investigate the development of each antibody lineage over time, we used SONAR’s module 3 features for phylogeny analysis. SONAR includes a specific version of the Imcantation Framework’s IgPhyML (*52, 53*) and an associated script for inferring ancestor sequences for each tree node. SONAR’s procedure creates a tree using IgPhyML’s antibody-specific HLP19 codon substitution model (*52*) and a nucleotide sequence for each internal node in that tree, representing an inferred ancestor sequence for the clade beneath that node. The internal node closest to the tree root corresponds to a hypothetical UCA sequence as inferred by the software. The identified lineage member sequences from module 2 were gathered across time points for each subject and locus and labeled with a prefix for each time point to create a unified set of lineage member sequences discovered from the repertoire over time. For a first-pass phylogenetic analysis, we used SONAR’s default behavior, with a MUSCLE alignment of all lineage members including the repertoire-discovered sequences and the “native” mature isolated members, followed by IgPhyML with the tree rooted on our own assigned germline V sequence from that macaque’s germline reference. To refine the tree and inference, we then supplied IgPhyML with a manually adjusted codon alignment of all lineage member sequences and used a draft UCA sequence as the tree root rather than the germline V segment only. To avoid biasing the final tree and inferences, we replaced possible nontemplated nucleotides with N in this root sequence. The earliest inferred ancestor sequence from this final step was compared with all known germline information for the relevant macaque (as described above for IgDiscover and MINING-D) to produce a final inference for the UCA of the lineage. This included any necessary adjustments to revert suspected mutated positions in germline-templated areas to the germline state. For final visualization purposes, we used trees rooted on the full UCA sequence. Indels in relevant lineages also required extensive manual review, as insertions and deletions are not modeled by HLP19 or the ancestor sequence inference procedure. To account for this, we reviewed the observed sequences of lineage members in each tree, and attempted to infer the most plausible explanation using qualitative background knowledge on the frequency of indels in lineage maturation and possible specific events such as insertions through duplications of adjacent nucleotide motifs (*9, 30, 57*). The output from this step, representing the final lineage tracing results for one lineage and chain, was a FASTA file for the alignment used for each tree, the associated Newick-format tree file from IgPhyML, and a FASTA of the inferred sequences for each tree node. This also included our final UCA sequence for heavy and light chains for each lineage.

#### Software and Configuration

- Custom Python program for common tasks in our workflow such as demultiplexing and adapter trimming: https://github.com/shawhahnlab/igseq

◦ cutadapt (*94*): https://github.com/marcelm/cutadapt
◦ PEAR (*95*): https://cme.h-its.org/exelixis/web/software/pear/doc.html
- Snakemake-based (*96*) workflow files for ongoing lineage tracing analyses: https://github.com/shawhahnlab/igseqhelper
- Singularity (*97*) for running SONAR’s Docker container
- KIMDB 1.1 for IGH germline reference sequences: http://kimdb.gkhlab.se/datasets/ and igseq’s data/germ/rhesus/kimdb files
- Ramesh et al. 2017 IGK and IGL as supplied by SONAR’s included germDB directory and igseq’s data/germ/rhesus/sonarramesh files
- Dedicated code repository for the work presented here: https://github.com/ShawHahnLab/Habib_et_al_2025_igseq

The dedicated repository provides the full detail and commands needed for reproduction of the results reported here.

### Env single-genome sequencing

Using the single genome sequencing (SGS) method described previously (*18, 29*), we amplified SHIV 3′half genomes from the sequential plasma samples of 103 SHIV-infected animals. Briefly, viral RNA was extracted from plasma using QIAamp Viral RNA kit (Qiagen) and reverse transcribed using SuperScript III Reverse Transcriptase (Invitrogen). Viral cDNA was then endpoint diluted and amplified using nested PCR with primers and conditions as previously reported. Geneious Prime software was used for sequence analysis. Pixel plots of SGS data were generated using the LANL Pixel plot tools (https://www.hiv.lanl.gov/content/sequence/pixel/pixel.html).

### Env SGS Hamming distance analysis

SGS data was analyzed by translating each Env’s nucleotide sequence into amino acids and aligning it to the appropriate infecting Env for each individual. Regions of interest such as the C-strand (HXB2 numbering 166-171), V3-glycan (324–334), fusion peptide (512–525), and MPER (656–683) were extracted. The hamming distance between the infecting Env and each sequence was calculated and used to find the mean hamming distance within each timepoint. Mean hamming distance was then calculated by taking the mean of all timepoints within each individual. These values were then compared between individuals with and without V2-apex bNAbs using a two-tailed Welch’s t-test. Additionally, we repeated this process for all continuous 10 amino acid regions of Env (i.e. 1-10, 2-11, 3-12, etc…). Gaps in the alignment were treated as individual differences for the purpose of this analysis. Custom scripts used to perform these calculations can be found here: https://github.com/ShawHahnLab/Habib_et_al_2025_HammingDistance.git

### NS-EMPEM

For negative stain EM polyclonal epitope mapping, 1 mg of polyclonal serum Fab was mixed with 10-20 µg of Env SOSIP, incubated overnight at 4 °C and then fractionated by size exclusion chromotography over a Superose 6 Increase 10/300 column. Running buffer was PBS with 500 mM Na2SO4 added. Fractions corresponding to the presumptive Fab-Env complex were pooled and concentrated in a 0.5-ml, 100-kDa spin concentrator to ∼0.2 mg/ml, then applied to a glow-discharged carbon-coated EM grid for 10-12 second, then blotted, and stained with 2 g/dL uranyl formate for 1 min, blotted and air-dried. Grids were examined on a Philips EM420 electron microscope operating at 120 kV and nominal magnification of 49,000x, and ∼100-400 images were collected on a 76 Mpix CCD camera at 2.4 Å/pixel. Images were analyzed and 3D refinements obtained using standard protocols with Relion 3.0 (Zivanov et al. 2018. *eLife*. 7:e42166).

### Cryo-EM sample preparation and data collection

The structure of V033-a.I1 in complex with Q23.17 MD39 envelope trimer was determined using single-particle cryo-EM. Previous work revealed mature lineage member V033-a.01 to bind multiple Fabs per trimer (*28*). To determine if V033 lineage IgG could engage a single envelope trimer through both antigen-binding arms, V033-a.I1 F(ab)’2 was generated by pepsin digestion and complexed with Q23.17 MD39 at 1.5:1 F(ab)’2-to-trimer molar ratio for a final trimer concentration of 2.5 mg/mL. After incubation for 30 minutes at 4C, the detergent n-Dodecyl β-D-maltoside (DDM) was added to a final concentration of 0.005% (w/v). 3µL of sample was added to copper C-flat Holey carbon-coated grids (CF-1.2/1.3 300 mesh; EMS) that were glow discharged with a PELCO easiGlow. Grids were blotted for 3 seconds at room temperature with 100% humidity and vitrified by liquid ethane using a Vitrobot Mark IV. Single particle cryo-EM datasets were collected on a FEI Titan Krios 300 kV cryo-transmission electron microscope equipped with a Gatan K3 direct electron detector. Movies were collected in counting mode using Leginon (PMID: 15890530) with a total dose of 58 e-/Å2 fractionated over 50 raw frames, with defocus values set to cycle between −0.80 and −2.0 μm. Approximately 7,000 micrographs were collected in 20 hours.

### Cryo-EM data processing and model building

All processing was done in cryoSPARC v3.4 (*98*), including micrograph curation, motion correction, CTF estimation, non-templated blob particle picking, 2D classification, ab initio modeling, and iterative 3D refinements. All homogenous and non-uniform 3D refinements of were performed using C1 symmetry. Map resolution was estimated using the 0.143 Fourier shell correlation (FSC) cutoff criterion. Despite using F(ab)’2, the majority of particle complexes contained a single arm bound to trimer without any visible density for the remainder of the F(ab)’2 molecule; therefore, a final 3D reconstruction with a single Fab domain was used for atomic modeling building. The initial coordinates for the V033-a.I1 complex were obtained by docking the V033-a.01 Fab from PDB-9BNP and the Q23.17 MD39 trimer from PDB-9BNL into the present cryo-EM density using UCSF ChimeraX (*99*). The atomic model was solved by iterative manual rebuilding in Coot (*100*) and real-space refinement in Phenix (*101*). Overall structure quality was assessed using MolProbity (*102*) and EMRinger (*103*).

## List of Supplementary Materials

Fig. S1. Epitope specificity of heterologous neutralization in SHIV-infected RMs.

Fig. S2. Inference of V2 apex bNAb UCAs.

Fig. S3. Longitudinal somatic hypermutation of V2 apex bNAb lineages.

Fig. S4. Longitudinal V033-a lineage heavy chain sequences.

Fig. S5. Env-Antibody Coevolution in RM V033.

Fig. S6. Affinity changes during V033-a lineage maturation.

Fig. S7. Env sensitivity and resistance signatures to the V033-a bNAb lineage.

Fig. S8. Structural maturation of the V033-a lineage.

Fig. S9. Heavy chain phylogenetic trees of V2 apex bNAb lineages.

Fig. S10. A two-residue insertion in necessary for heterologous neutralization by the V031-a.01 mAb.

Fig S11. Mutational Bottleneck in the 42056-a lineage.

Fig. S12. Neutralization phenotype of inferred V2 apex bNAb UCAs.

Fig. S13. T/F V1V2 sequences are usually absent preceding bNAb initiation.

Fig. S14. N130 glycan deletion precedes V2 apex bNAb elicitation in SHIV-CH505-infected RMs.

Fig. S15. V1V2 sequence evolution in RM 6070.

Fig. S16. V1V2 sequence evolution in RM T646.

Fig. S17. Limited and convergent Env mutations guide rhesus V2 apex bNAb maturation.

Fig. S18. Limited and convergent Env mutations guide human V2 apex bNAb maturation.

Fig. S19. Env selection correlates with bNAb structural footprints.

Fig. S20. A K190E V2 hypervariable loop mutation mediates initial escape from the 6561-a bNAb lineage.

Fig. S21. Longitudinal C-strand Hamming distances in SHIV-infected RMs.

Fig. S22. Evidence of C-strand targeted NAbs in five RMs.

Fig. S23. C-strand selection in SHIV CH848-infected RMs usually consists of an E169K reversion to group M consensus.

Fig. S24. C-strand selection in SHIV B41-infected RMs.

Fig. S25. Selection at other bNAb epitopes is not significantly associated with the development of V2 apex bNAbs.

Fig. S26. Model of V2 apex bNAb priming

Fig. S27. An early fitness reversion in SHIV-Q23.17 infected RMs.

Fig. S28. Transient C-strand selection in SHIV-Ce1086 infected RMs reverts after replacement of the N160 glycan.

Fig. S29. Cryo-EM details of V033-a.I1 in complex with Q23.17 MD39 envelope trimer.

Table. S1. Summary of the 122 SHIV-infected RMs analyzed in this study.

Table. S2. Summary of previously described rhesus V2 apex bNAbs.

Table. S3. Cryo-EM data collection, processing, and refinement validation statistics

## Acknowledgements

We thank Dr. James Theiler for statistical advice. We thank the staff at Bioqual for exceptional care of nonhuman primates and collection of clinical specimens. We thank the University of Pennsylvania flow cytometry core for technical assistance with B cell sorting.

## Author Contributions

RH, RSR, HL, MPH, KJS, SJP, FBR, AS, CLM, JR, NC, JL, WD, MSC, and CZ contributed to rhesus plasma and monoclonal antibody neutralization assays. RSR, RJE, LS, PDK contributed to structural analyses. JL, BL, RRC, KA contributed to binding assays and affinity measurements. RH, AJC, EVI, KW (Wiehe) contributed to phylogenetic analyses. RH, HL, SW, JWC, KA, YL contributed to Env single genome sequencing. RH, RSR, MPH, LM, ANS, HL, JWC, FBR, WBW, KW (Wiehe), contributed to B cell sorting and next-generation sequencing. YP, WL contributed to antibody production and synthesis. RH, AJC, MPH, KW (Wagh), CJA, ZS, BTK, contributed to statistical analyses. KJS, SJP, FBR, AS, CLM, JR, NC, JL, WD, MSC, CZ ML contributed to rhesus macaque sample collection and processing. KJB, WBW, KW (Wiehe), KOS, RJE, DWC, ML, FDB, DB, RA, DWK, BFH, BTK, LS, PDK, BHH, and GMS supervised the work. KJB, WBW, RA, BFH, LS, PDK, BHH, and GMS contributed to funding acquisition. RH and GMS reviewed and analyzed all data and wrote the initial drafts of the manuscript. All authors contributed to manuscript review and editing.

## Data and materials availability

All SHIV sequences, plasmid DNA clones of SHIVs, and infectious stocks of SHIVs have been deposited in Genbank, the NIH AIDS Research and Reference Reagent Program, and ATCC/American Tissue Culture Collection, as described (*19*). V2 apex bNAb UCA sequences are deposited at Genbank under accession numbers PV467379-PV467402. The B cell next-generation sequencing data generated in this study are available under the NCBI Bioproject PRJNA1121265. The atomic model of the V033-a.I1/Q23.MD39 complex generated in this study is available at the Protein Data Bank (PDB, https://www.rcsb.org) under the PDB accession code 9OMG. The corresponding cryo-EM reconstruction is available at the Electron Microscopy Data Bank (EMDB, https://www.ebi.ac.uk/emdb/) under the EMDB access code 70613. Longitudinal Env SGS gp140 sequences are deposited at Genbank under accession numbers XXXX-XXXX.

## References and Notes

1. E. S. Gray, M. C. Madiga, T. Hermanus, P. L. Moore, C. K. Wibmer, N. L. Tumba, L. Werner, K. Mlisana, S. Sibeko, C. Williamson, S. S. A. Karim, L. Morris, The CAPRISA002 Study Team, The neutralization breadth of HIV-1 develops incrementally over four years and is associated with CD4+ T cell decline and high viral load during acute infection. Journal of Virology 85, 4828–4840 (2011).

2. N. A. Doria-Rose, R. M. Klein, M. G. Daniels, S. O’Dell, M. Nason, A. Lapedes, T. Bhattacharya, S. A. Migueles, R. T. Wyatt, B. T. Korber, J. R. Mascola, M. Connors, Breadth of human immunodeficiency virus-specific neutralizing activity in sera: clustering analysis and association with clinical variables. Journal of Virology 84, 1631–1636 (2010).

3. I. Mikell, D. N. Sather, S. A. Kalams, M. Altfeld, G. Alter, L. Stamatatos, Characteristics of the earliest cross-neutralizing antibody response to HIV-1. PLoS Pathogens 7, e1001251 (2011).

4. E. Landais, X. Huang, C. Havenar-Daughton, B. Murrell, M. A. Price, L. Wickramasinghe, A. Ramos, C. B. Bian, M. Simek, S. Allen, E. Karita, W. Kilembe, S. Lakhi, M. Inambao, A. Kamali, E. J. Sanders, O. Anzala, V. Edward, L.-G. Bekker, J. Tang, J. Gilmour, S. L. Kosakovsky-Pond, P. Phung, T. Wrin, S. Crotty, A. Godzik, P. Poignard, Broadly neutralizing antibody responses in a large longitudinal sub-Saharan HIV primary infection cohort. PLOS Pathogens 12, e1005369 (2016).

5. G. D. Tomaras, J. M. Binley, E. S. Gray, E. T. Crooks, K. Osawa, P. L. Moore, N. Tumba, T. Tong, X. Shen, N. L. Yates, J. Decker, C. K. Wibmer, F. Gao, S. M. Alam, P. Easterbrook, S. A. Karim, G. Kamanga, J. A. Crump, M. Cohen, G. M. Shaw, J. R. Mascola, B. F. Haynes, D. C. Montefiori, L. Morris, Polyclonal B cell responses to conserved neutralization epitopes in a subset of HIV-1-infected individuals. Journal of Virology 85, 11502–11519 (2011).

6. P. Rusert, R. D. Kouyos, C. Kadelka, H. Ebner, M. Schanz, M. Huber, D. L. Braun, N. Hozé, A. Scherrer, C. Magnus, J. Weber, T. Uhr, V. Cippa, C. W. Thorball, H. Kuster, M. Cavassini, E. Bernasconi, M. Hoffmann, A. Calmy, M. Battegay, A. Rauch, S. Yerly, V. Aubert, T. Klimkait, J. Böni, J. Fellay, R. R. Regoes, H. F. Günthard, A. Trkola, P. Rusert, R. D. Kouyos, C. Kadelka, H. Ebner, M. Schanz, M. Huber, D. L. Braun, N. Hozé, A. Scherrer, C. Magnus, J. Weber, T. Uhr, V. Cippa, C. W. Thorball, H. Kuster, M. Cavassini, E. Bernasconi, M. Hoffmann, A. Calmy, M. Battegay, A. Rauch, S. Yerly, V. Aubert, T. Klimkait, J. Böni, J. Fellay, R. R. Regoes, H. F. Günthard, A. Trkola, Determinants of HIV-1 broadly neutralizing antibody induction. Nature Medicine 22, 1260–1267 (2016).

7. R. D. Kouyos, P. Rusert, C. Kadelka, M. Huber, A. Marzel, H. Ebner, M. Schanz, T. Liechti, N. Friedrich, D. L. Braun, A. U. Scherrer, J. Weber, T. Uhr, N. S. Baumann, C. Leemann, H. Kuster, J.-P. Chave, M. Cavassini, E. Bernasconi, M. Hoffmann, A. Calmy, M. Battegay, A. Rauch, S. Yerly, V. Aubert, T. Klimkait, J. Böni, K. J. Metzner, H. F. Günthard, A. Trkola, R. D. Kouyos, P. Rusert, C. Kadelka, M. Huber, A. Marzel, H. Ebner, M. Schanz, T. Liechti, N. Friedrich, D. L. Braun, A. U. Scherrer, J. Weber, T. Uhr, N. S. Baumann, C. Leemann, H. Kuster, J.-P. Chave, M. Cavassini, E. Bernasconi, M. Hoffmann, A. Calmy, M. Battegay, A. Rauch, S. Yerly, V. Aubert, T. Klimkait, J. Böni, K. J. Metzner, H. F. Günthard, A. Trkola, Tracing HIV-1 strains that imprint broadly neutralizing antibody responses. Nature 561, 406–410 (2018).

8. P. L. Moore, C. Williamson, L. Morris, Virological features associated with the development of broadly neutralizing antibodies to HIV-1. Trends in Microbiology 23, 204–211 (2015).

9. C. Joyce, S. Murrell, B. Murrell, O. Omorodion, L. S. Ver, N. Carrico, R. Bastidas, R. Nedellec, M. Bick, J. Woehl, F. Zhao, A. Burns, S. Barman, M. Appel, A. Ramos, L. Wickramasinghe, K. Eren, T. Vollbrecht, D. M. Smith, S. L. K. Pond, R. McBride, C. Worth, F. Batista, D. Sok, The IAVI Protocol C Investigators & The IAVI African HIV Research Network, P. Poignard, B. Briney, I. A. Wilson, E. Landais, D. R. Burton, Antigen pressure from two founder viruses induces multiple insertions at a single antibody position to generate broadly neutralizing HIV antibodies. PLOS Pathogens 19, e1011416 (2023).

10. V. Cortez, K. Odem-Davis, R. S. McClelland, W. Jaoko, J. Overbaugh, HIV-1 superinfection in women broadens and strengthens the neutralizing antibody response. PLOS Pathogens 8, e1002611 (2012).

11. D. N. Sather, J. Armann, L. K. Ching, A. Mavrantoni, G. Sellhorn, Z. Caldwell, X. Yu, B. Wood, S. Self, S. Kalams, L. Stamatatos, Factors associated with the development of cross-reactive neutralizing antibodies during human immunodeficiency virus type 1 infection. Journal of Virology 83, 757–769 (2009).

12. K. Wagh, E. F. Kreider, Y. Li, H. J. Barbian, G. H. Learn, E. Giorgi, P. T. Hraber, T. G. Decker, A. G. Smith, M. V. Gondim, L. Gillis, J. Wandzilak, G.-Y. Chuang, R. Rawi, F. Cai, P. Pellegrino, I. Williams, J. Overbaugh, F. Gao, P. D. Kwong, B. F. Haynes, G. M. Shaw, P. Borrow, M. S. Seaman, B. H. Hahn, B. Korber, Completeness of HIV-1 Envelope glycan shield at transmission determines neutralization breadth. Cell Reports 25, 893–908.e897 (2018).

13. B. F. Haynes, K. Wiehe, P. Borrow, K. O. Saunders, B. Korber, K. Wagh, A. J. McMichael, G. Kelsoe, B. H. Hahn, F. Alt, G. M. Shaw, B. F. Haynes, K. Wiehe, P. Borrow, K. O. Saunders, B. Korber, K. Wagh, A. J. McMichael, G. Kelsoe, B. H. Hahn, F. Alt, G. M. Shaw, Strategies for HIV-1 vaccines that induce broadly neutralizing antibodies. Nature Reviews Immunology 23, 142–158 (2022).

14. D. Sok, D. R. Burton, Recent progress in broadly neutralizing antibodies to HIV. Nature Immunology 19, 1179–1188 (2018).

15. P. D. Kwong, J. R. Mascola, HIV-1 vaccines based on antibody identification, B cell ontogeny, and epitope structure. Immunity 48, 855–871 (2018).

16. G. Y. Chuang, J. Zhou, P. Acharya, R. Rawi, C. H. Shen, Z. Sheng, B. Zhang, T. Zhou, R. T. Bailer, V. P. Dandey, N. A. Doria-Rose, M. K. Louder, K. McKee, J. R. Mascola, L. Shapiro, P. D. Kwong, Structural survey of broadly neutralizing antibodies targeting the HIV-1 Env trimer delineates epitope categories and characteristics of recognition. Structure 27, 196–206.e196 (2019).

17. L. M. Walker, M. D. Simek, F. Priddy, J. S. Gach, D. Wagner, M. B. Zwick, S. K. Phogat, P. Poignard, D. R. Burton, A limited number of antibody specificities mediate broad and potent serum neutralization in selected HIV-1 infected individuals. PLoS Pathogens 6, e1001028 (2010).

18. H. Li, S. Wang, R. Kong, W. Ding, F. H. Lee, Z. Parker, E. Kim, G. H. Learn, P. Hahn, B. Policicchio, E. Brocca-Cofano, C. Deleage, X. Hao, G. Y. Chuang, J. Gorman, M. Gardner, M. G. Lewis, T. Hatziioannou, S. Santra, C. Apetrei, I. Pandrea, S. M. Alam, H. X. Liao, X. Shen, G. D. Tomaras, M. Farzan, E. Chertova, B. F. Keele, J. D. Estes, J. D. Lifson, R. W. Doms, D. C. Montefiori, B. F. Haynes, J. G. Sodroski, P. D. Kwong, B. H. Hahn, G. M. Shaw, Envelope residue 375 substitutions in simian-human immunodeficiency viruses enhance CD4 binding and replication in rhesus macaques. Proceedings of the National Academy of Sciences 113, E3413–E3422 (2016).

19. H. Li, S. Wang, F.-H. Lee, R. S. Roark, A. I. Murphy, J. Smith, C. Zhao, J. Rando, N. Chohan, Y. Ding, E. Kim, E. Lindemuth, K. J. Bar, I. Pandrea, C. Apetrei, B. F. Keele, J. D. Lifson, M. G. Lewis, T. N. Denny, B. F. Haynes, B. H. Hahn, G. M. Shaw, New SHIVs and improved design strategy for modeling HIV-1 transmission, immunopathogenesis, prevention and cure. Journal of Virology 95, e00071–00021 (2021).

20. B. F. Haynes, G. Kelsoe, S. C. Harrison, T. B. Kepler, B-cell–lineage immunogen design in vaccine development with HIV-1 as a case study. Nature biotechnology 30, 423–433 (2012).

21. M. Bonsignori, E. F. Kreider, D. Fera, R. R. Meyerhoff, T. Bradley, K. Wiehe, S. M. Alam, B. Aussedat, W. E. Walkowicz, K. K. Hwang, K. O. Saunders, R. Zhang, M. A. Gladden, A. Monroe, A. Kumar, S. M. Xia, M. Cooper, M. K. Louder, K. McKee, R. T. Bailer, B. W. Pier, C. A. Jette, G. Kelsoe, W. B. Williams, L. Morris, J. Kappes, K. Wagh, G. Kamanga, M. S. Cohen, P. T. Hraber, D. C. Montefiori, A. Trama, H. X. Liao, T. B. Kepler, M. A. Moody, F. Gao, S. J. Danishefsky, J. R. Mascola, G. M. Shaw, B. H. Hahn, S. C. Harrison, B. T. Korber, B. F. Haynes, Staged induction of HIV-1 glycan-dependent broadly neutralizing antibodies. Science Translational Medicine 9, eaai7514 (2017).

22. M. Bonsignori, T. Zhou, Z. Sheng, L. Chen, F. Gao, M. G. Joyce, G. Ozorowski, G.-Y. Chuang, Chaim A. Schramm, K. Wiehe, S. M. Alam, T. Bradley, Morgan A. Gladden, K.-K. Hwang, S. Iyengar, A. Kumar, X. Lu, K. Luo, Michael C. Mangiapani, Robert J. Parks, H. Song, P. Acharya, Robert T. Bailer, A. Cao, A. Druz, Ivelin S. Georgiev, Young D. Kwon, Mark K. Louder, B. Zhang, A. Zheng, Brenna J. Hill, R. Kong, C. Soto, James C. Mullikin, Daniel C. Douek, David C. Montefiori, Michael A. Moody, George M. Shaw, Beatrice H. Hahn, G. Kelsoe, Peter T. Hraber, Bette T. Korber, Scott D. Boyd, Andrew Z. Fire, Thomas B. Kepler, L. Shapiro, Andrew B. Ward, John R. Mascola, H.-X. Liao, Peter D. Kwong, Barton F. Haynes, Maturation pathway from germline to broad HIV-1 neutralizer of a CD4-mimic antibody. Cell 165, 449–463 (2016).

23. N. A. Doria-Rose, C. A. Schramm, J. Gorman, P. L. Moore, J. N. Bhiman, B. J. DeKosky, M. J. Ernandes, I. S. Georgiev, H. J. Kim, M. Pancera, R. P. Staupe, H. R. Altae-Tran, R. T. Bailer, E. T. Crooks, A. Cupo, A. Druz, N. J. Garrett, K. H. Hoi, R. Kong, M. K. Louder, N. S. Longo, K. McKee, M. Nonyane, S. O’Dell, R. S. Roark, R. S. Rudicell, S. D. Schmidt, D. J. Sheward, C. Soto, C. K. Wibmer, Y. Yang, Z. Zhang, J. C. Mullikin, J. M. Binley, R. W. Sanders, I. A. Wilson, J. P. Moore, A. B. Ward, G. Georgiou, C. Williamson, S. S. A. Karim, L. Morris, P. D. Kwong, L. Shapiro, J. R. Mascola, Developmental pathway for potent V1V2-directed HIV-neutralizing antibodies. Nature 508, 55–62 (2014).

24. H. X. Liao, R. Lynch, T. Zhou, F. Gao, S. Munir Alam, S. D. Boyd, A. Z. Fire, K. M. Roskin, C. A. Schramm, Z. Zhang, J. Zhu, L. Shapiro, J. C. Mullikin, S. Gnanakaran, P. Hraber, K. Wiehe, G. Kelsoe, G. Yang, S. M. Xia, D. C. Montefiori, R. Parks, K. E. Lloyd, R. M. Scearce, K. A. Soderberg, M. Cohen, G. Kamanga, M. K. Louder, L. M. Tran, Y. Chen, F. Cai, S. Chen, S. Moquin, X. Du, M. Gordon Joyce, S. Srivatsan, B. Zhang, A. Zheng, G. M. Shaw, B. H. Hahn, T. B. Kepler, B. T. M. Korber, P. D. Kwong, J. R. Mascola, B. F. Haynes, J. Becker, B. Benjamin, R. Blakesley, G. Bouffard, S. Brooks, H. Coleman, M. Dekhtyar, M. Gregory, X. Guan, J. Gupta, J. Han, A. Hargrove, S. L. Ho, T. Johnson, R. Legaspi, S. Lovett, Q. Maduro, C. Masiello, B. Maskeri, J. McDowell, C. Montemayor, J. Mulliki, M. Park, N. Riebow, K. Schandler, B. Schmidt, C. Sison, M. Stantripop, J. Thomas, P. Thomas, M. Vemulapalli, A. Young, Co-evolution of a broadly neutralizing HIV-1 antibody and founder virus. Nature 496, 469–476 (2013).

25. N. A. Doria-Rose, E. Landais, Coevolution of HIV-1 and broadly neutralizing antibodies. Current Opinion in HIV and AIDS 14, 286–293 (2019).

26. F. Gao, M. Bonsignori, H.-X. Liao, A. Kumar, S.-M. Xia, X. Lu, F. Cai, K.-K. Hwang, H. Song, T. Zhou, Rebecca M. Lynch, S. M. Alam, M. A. Moody, G. Ferrari, M. Berrong, G. Kelsoe, George M. Shaw, Beatrice H. Hahn, David C. Montefiori, G. Kamanga, Myron S. Cohen, P. Hraber, Peter D. Kwong, Bette T. Korber, John R. Mascola, Thomas B. Kepler, Barton F. Haynes, Cooperation of B cell lineages in induction of HIV-1-broadly neutralizing antibodies. Cell 158, 481–491 (2014).

27. R. S. Roark, H. Li, W. B. Williams, H. Chug, R. D. Mason, J. Gorman, S. Wang, F. H. Lee, J. Rando, M. Bonsignori, K. K. Hwang, K. O. Saunders, K. Wiehe, M. A. Moody, P. T. Hraber, K. Wagh, E. E. Giorgi, R. M. Russell, F. Bibollet-Ruche, W. Liu, J. Connel, A. G. Smith, J. DeVoto, A. I. Murphy, J. Smith, W. Ding, C. Zhao, N. Chohan, M. Okumura, C. Rosario, Y. Ding, E. Lindemuth, A. M. Bauer, K. J. Bar, D. Ambrozak, C. W. Chao, G. Y. Chuang, H. Geng, B. C. Lin, M. K. Louder, R. Nguyen, B. Zhang, M. G. Lewis, D. D. Raymond, N. A. Doria-Rose, C. A. Schramm, D. C. Douek, M. Roederer, T. B. Kepler, G. Kelsoe, J. R. Mascola, P. D. Kwong, B. T. Korber, S. C. Harrison, B. F. Haynes, B. H. Hahn, G. M. Shaw, Recapitulation of HIV-1 Env-antibody coevolution in macaques leading to neutralization breadth. Science 371, eabd2638 (2021).

28. R. S. Roark, R. Habib, J. Gorman, H. Li, A. J. Connell, M. Bonsignori, Y. Guo, M. P. Hogarty, A. S. Olia, K. Sowers, B. Zhang, F. Bibollet-Ruche, S. Callaghan, J. W. Carey, G. Cerutti, D. R. Harris, W. He, E. Lewis, T. Liu, R. D. Mason, Y. Park, J. M. Rando, A. Singh, J. Wolff, Q. P. Lei, M. K. Louder, N. A. Doria-Rose, R. Andrabi, K. O. Saunders, M. S. Seaman, B. F. Haynes, D. W. Kulp, J. R. Mascola, M. Roederer, Z. Sheng, B. H. Hahn, G. M. Shaw, P. D. Kwong, L. Shapiro, Structural and genetic basis of HIV-1 envelope V2 apex recognition by rhesus broadly neutralizing antibodies. The Journal of Experimental Medicine 222, e20250638 (2025).

29. B. F. Keele, E. E. Giorgi, J. F. Salazar-Gonzalez, J. M. Decker, K. T. Pham, M. G. Salazar, C. Sun, T. Grayson, S. Wang, H. Li, X. Wei, C. Jiang, J. L. Kirchherr, F. Gao, J. A. Anderson, L. H. Ping, R. Swanstrom, G. D. Tomaras, W. A. Blattner, P. A. Goepfert, J. M. Kilby, M. S. Saag, E. L. Delwart, M. P. Busch, M. S. Cohen, D. C. Montefiori, B. F. Haynes, B. Gaschen, G. S. Athreya, H. Y. Lee, N. Wood, C. Seoighe, A. S. Perelson, T. Bhattacharya, B. T. Korber, B. H. Hahn, G. M. Shaw, Identification and characterization of transmitted and early founder virus envelopes in primary HIV-1 infection. Proceedings of the National Academy of Sciences 105, 7552–7557 (2008).

30. D. J. Morris, J. Gorman, T. Zhou, J. Lora, A. J. Connell, H. Li, W. Liu, R. S. Roark, M. S. Campion, J. W. Carey, R. Habib, Y. Li, C. L. Martella, Y. Park, A. Singh, K. J. Sowers, I.-T. Teng, S. Wang, N. Chohan, W. Ding, C. Lauer, E. Lewis, R. D. Mason, J. M. Rando, L. Peyton, C. A. Schramm, K. Wagh, B. Korber, M. S. Seaman, D. C. Douek, B. F. Haynes, D. W. Kulp, M. Roederer, B. H. Hahn, P. D. Kwong, G. M. Shaw, Transient glycan-shield reduction induces CD4-binding site broadly neutralizing antibodies in SHIV-infected macaques. Cell Reports 44, 115848 (2024).

31. J. Gorman, C. Soto, M. M. Yang, T. M. Davenport, M. Guttman, R. T. Bailer, M. Chambers, G. Y. Chuang, B. J. Dekosky, N. A. Doria-Rose, A. Druz, M. J. Ernandes, I. S. Georgiev, M. C. Jarosinski, M. G. Joyce, T. M. Lemmin, S. Leung, M. K. Louder, J. R. McDaniel, S. Narpala, M. Pancera, J. Stuckey, X. Wu, Y. Yang, B. Zhang, T. Zhou, J. C. Mullikin, U. Baxa, G. Georgiou, A. B. McDermott, M. Bonsignori, B. F. Haynes, P. L. Moore, L. Morris, K. K. Lee, L. Shapiro, J. R. Mascola, P. D. Kwong, Structures of HIV-1 Env V1V2 with broadly neutralizing antibodies reveal commonalities that enable vaccine design. Nature Structural and Molecular Biology 23, 81–90 (2016).

32. J. E. Voss, R. Andrabi, L. E. McCoy, N. de Val, R. P. Fuller, T. Messmer, C. Y. Su, D. Sok, S. N. Khan, F. Garces, L. K. Pritchard, R. T. Wyatt, A. B. Ward, M. Crispin, I. A. Wilson, D. R. Burton, Elicitation of neutralizing antibodies targeting the V2 apex of the HIV Envelope trimer in a wild-type animal model. Cell Reports 21, 222–235 (2017).

33. M. Bonsignori, K. K. Hwang, X. Chen, C. Y. Tsao, L. Morris, E. Gray, D. J. Marshall, J. A. Crump, S. H. Kapiga, N. E. Sam, F. Sinangil, M. Pancera, Y. Yongping, B. Zhang, J. Zhu, P. D. Kwong, S. O’Dell, J. R. Mascola, L. Wu, G. J. Nabel, S. Phogat, M. S. Seaman, J. F. Whitesides, M. A. Moody, G. Kelsoe, X. Yang, J. Sodroski, G. M. Shaw, D. C. Montefiori, T. B. Kepler, G. D. Tomaras, S. M. Alam, H. X. Liao, B. F. Haynes, Analysis of a clonal lineage of HIV-1 envelope V2/V3 conformational epitope-specific broadly neutralizing antibodies and their inferred unmutated common ancestors. Journal of Virology 85, 9998–10009 (2011).

34. K. O. Saunders, L. K. Verkoczy, C. Jiang, J. Zhang, R. Parks, H. Chen, M. Housman, H. Bouton-Verville, X. Shen, A. M. Trama, R. Scearce, L. Sutherland, S. Santra, A. Newman, A. Eaton, K. Xu, I. S. Georgiev, M. G. Joyce, G. D. Tomaras, M. Bonsignori, S. G. Reed, A. Salazar, J. R. Mascola, M. A. Moody, D. W. Cain, M. Centlivre, S. Zurawski, G. Zurawski, H. P. Erickson, P. D. Kwong, S. M. Alam, Y. Levy, D. C. Montefiori, B. F. Haynes, Vaccine induction of heterologous tier 2 HIV-1 neutralizing antibodies in animal models. Cell Reports 21, 3681–3690 (2017).

35. R. Andrabi, J. E. Voss, C. H. Liang, B. Briney, L. E. McCoy, C. Y. Wu, C. H. Wong, P. Poignard, D. R. Burton, Identification of common features in prototype broadly neutralizing antibodies to HIV Envelope V2 apex to facilitate vaccine design. Immunity 43, 959–973 (2015).

36. C. R. Brown, M. Czapiga, J. Kabat, Q. Dang, I. Ourmanov, Y. Nishimura, M. A. Martin, V. M. Hirsch, Unique pathology in simian immunodeficiency virus-infected rapid progressor macaques is consistent with a pathogenesis distinct from that of classical AIDS. Journal of Virology 81, 5594–5606 (2007).

37. V. M. Hirsch, S. Santra, S. Goldstein, R. Plishka, A. Buckler-White, A. Seth, I. Ourmanov, C. R. Brown, R. Engle, D. Montefiori, J. Glowczwskie, K. Kunstman, S. Wolinsky, N. L. Letvin, Immune failure in the absence of profound CD4+ T-lymphocyte depletion in simian immunodeficiency virus-infected rapid progressor macaques. Journal of Virology 78, 275–284 (2004).

38. M. G. Pauthner, J. P. Nkolola, C. Havenar-Daughton, B. Murrell, S. M. Reiss, R. Bastidas, J. Prévost, R. Nedellec, B. v. Bredow, P. Abbink, C. A. Cottrell, D. W. Kulp, T. Tokatlian, B. Nogal, M. Bianchi, H. Li, J. H. Lee, S. T. Butera, D. T. Evans, L. Hangartner, A. Finzi, I. A. Wilson, R. T. Wyatt, D. J. Irvine, W. R. Schief, A. B. Ward, R. W. Sanders, S. Crotty, G. M. Shaw, D. H. Barouch, D. R. Burton, Vaccine-induced protection from homologous tier 2 SHIV challenge in nonhuman primates depends on serum-neutralizing antibody titers. Immunity 50, 241–252.e246 (2019).

39. A. Pegu, B. Borate, Y. Huang, M. G. Pauthner, A. J. Hessell, B. Julg, N. A. Doria-Rose, S. D. Schmidt, L. N. Carpp, M. D. Cully, X. Chen, G. M. Shaw, D. H. Barouch, N. L. Haigwood, L. Corey, D. R. Burton, M. Roederer, P. B. Gilbert, J. R. Mascola, Y. Huang, A meta-analysis of passive immunization studies shows an association of serum neutralizing antibody titer with protection against SHIV challenge. Cell Host & Microbe 26, 336–346.e333 (2019).

40. B. K. Felber, Z. Lu, X. Hu, A. Valentin, M. Rosati, C. A. Remmel, J. A. Weiner, M. C. Carpenter, K. Faircloth, S. Stanfield-Oakley, W. B. Williams, X. Shen, G. D. Tomaras, C. C. LaBranche, D. Montefiori, H. V. Trinh, M. Rao, M. S. Alam, N. A. Vandergrift, K. O. Saunders, Y. Wang, W. Rountree, J. Das, G. Alter, S. G. Reed, P. P. Aye, F. Schiro, B. Pahar, J. P. Dufour, R. S. Veazey, P. A. Marx, D. J. Venzon, G. M. Shaw, G. Ferrari, M. E. Ackerman, B. F. Haynes, G. N. Pavlakis, Co-immunization of DNA and protein in the same anatomical sites induces superior protective immune responses against SHIV challenge. Cell Reports 31, 107624 (2020).

41. K. O. Saunders, L. Wang, M. G. Joyce, Z.-Y. Yang, A. B. Balazs, C. Cheng, S.-Y. Ko, W.-P. Kong, R. S. Rudicell, I. S. Georgiev, L. Duan, K. E. Foulds, M. Donaldson, L. Xu, S. D. Schmidt, J.-P. Todd, D. Baltimore, M. Roederer, A. T. Haase, P. D. Kwong, S. S. Rao, J. R. Mascola, G. J. Nabel, Broadly neutralizing Human Immunodeficiency Virus Type 1 antibody gene transfer protects nonhuman primates from mucosal Simian-Human Immunodeficiency Virus infection. Journal of Virology 89, 8334–8345 (2015).

42. J.-P. Julien, J. H. Lee, A. Cupo, C. D. Murin, R. Derking, S. Hoffenberg, M. J. Caulfield, C. R. King, A. J. Marozsan, P. J. Klasse, R. W. Sanders, J. P. Moore, I. A. Wilson, A. B. Ward, Asymmetric recognition of the HIV-1 trimer by broadly neutralizing antibody PG9. Proceedings of the National Academy of Sciences 110, 4351–4356 (2013).

43. J. S. McLellan, M. Pancera, C. Carrico, J. Gorman, J. P. Julien, R. Khayat, R. Louder, R. Pejchal, M. Sastry, K. Dai, S. O’Dell, N. Patel, S. Shahzad-Ul-Hussan, Y. Yang, B. Zhang, T. Zhou, J. Zhu, J. C. Boyington, G. Y. Chuang, D. Diwanji, I. Georgiev, Y. Do Kwon, D. Lee, M. K. Louder, S. Moquin, S. D. Schmidt, Z. Y. Yang, M. Bonsignori, J. A. Crump, S. H. Kapiga, N. E. Sam, B. F. Haynes, D. R. Burton, W. C. Koff, L. M. Walker, S. Phogat, R. Wyatt, J. Orwenyo, L. X. Wang, J. Arthos, C. A. Bewley, J. R. Mascola, G. J. Nabel, W. R. Schief, A. B. Ward, I. A. Wilson, P. D. Kwong, Structure of HIV-1 gp120 V1/V2 domain with broadly neutralizing antibody PG9. Nature 480, 336–343 (2011).

44. R. Pejchal, L. M. Walker, R. L. Stanfield, S. K. Phogat, W. C. Koff, P. Poignard, D. R. Burton, I. A. Wilson, R. Pejchal, L. M. Walker, R. L. Stanfield, S. K. Phogat, W. C. Koff, P. Poignard, D. R. Burton, I. A. Wilson, Structure and function of broadly reactive antibody PG16 reveal an H3 subdomain that mediates potent neutralization of HIV-1. Proceedings of the National Academy of Sciences 107, (2010).

45. J. Gorman, G.-Y. Chuang, Y.-T. Lai, C.-H. Shen, J. C. Boyington, A. Druz, H. Geng, M. K. Louder, K. McKee, R. Rawi, R. Verardi, Y. Yang, B. Zhang, N. A. Doria-Rose, B. Lin, P. L. Moore, L. Morris, L. Shapiro, J. R. Mascola, P. D. Kwong, Structure of super-potent antibody CAP256-VRC26.25 in complex with HIV-1 Envelope reveals a combined mode of trimer-apex recognition. Cell Reports 31, 107488 (2020).

46. E. Landais, B. Murrell, B. Briney, S. Murrell, K. Rantalainen, Z. T. Berndsen, A. Ramos, L. Wickramasinghe, M. L. Smith, K. Eren, N. de Val, M. Wu, A. Cappelletti, J. Umotoy, Y. Lie, T. Wrin, P. Algate, P. Y. Chan-Hui, E. Karita, A. B. Ward, I. A. Wilson, D. R. Burton, D. Smith, S. L. K. Pond, P. Poignard, HIV Envelope glycoform heterogeneity and localized diversity govern the initiation and maturation of a V2 apex broadly neutralizing antibody lineage. Immunity 47, 990–1003.e1009 (2017).

47. C. A. Schramm, Z. Sheng, Z. Zhang, J. R. Mascola, P. D. Kwong, L. Shapiro, SONAR: A high-throughput pipeline for inferring antibody ontogenies from longitudinal sequencing of B cell transcripts. Frontiers in Immunology 7, 372 (2016).

48. M. M. Corcoran, G. E. Phad, N. V. Bernat, C. Stahl-Hennig, N. Sumida, M. A. A. Persson, M. Martin, G. B. K. Hedestam, Production of individualized V gene databases reveals high levels of immunoglobulin genetic diversity. Nature Communications 7, 1–14 (2016).

49. N. Vázquez Bernat, M. Corcoran, U. Hardt, M. Kaduk, G. E. Phad, M. Martin, G. B. Karlsson Hedestam, High-quality library preparation for NGS-based immunoglobulin germline gene inference and repertoire expression analysis. Frontiers in Immunology 10, 660 (2019).

50. M. Lee, A. Changela, J. Gorman, R. Rawi, T. Bylund, C. W. Chao, B. C. Lin, M. K. Louder, A. S. Olia, B. Zhang, N. A. Doria-Rose, S. Zolla-Pazner, L. Shapiro, G.-Y. Chuang, P. D. Kwong, M. Lee, A. Changela, J. Gorman, R. Rawi, T. Bylund, C. W. Chao, B. C. Lin, M. K. Louder, A. S. Olia, B. Zhang, N. A. Doria-Rose, S. Zolla-Pazner, L. Shapiro, G.-Y. Chuang, P. D. Kwong, Extended antibody-framework-to-antigen distance observed exclusively with broad HIV-1-neutralizing antibodies recognizing glycan-dense surfaces. Nature Communications 12, 6470 (2021).

51. B. Briney, A. Inderbitzin, C. Joyce, D. R. Burton, B. Briney, A. Inderbitzin, C. Joyce, D. R. Burton, Commonality despite exceptional diversity in the baseline human antibody repertoire. Nature 566, 393–397 (2019).

52. K. B. Hoehn, J. A. V. Heiden, J. Q. Zhou, G. Lunter, O. G. Pybus, S. H. Kleinstein, K. B. Hoehn, J. A. Vander Heiden, J. Q. Zhou, G. Lunter, O. G. Pybus, S. H. Kleinstein, Repertoire-wide phylogenetic models of B cell molecular evolution reveal evolutionary signatures of aging and vaccination. Proceedings of the National Academy of Sciences 116, 22664–22672 (2019).

53. K. B. Hoehn, G. Lunter, O. G. Pybus, A phylogenetic codon substitution model for antibody lineages. Genetics 206, 417–427 (2017).

54. A. R. Ghosh, R. Habib, N. Mishra, S. Callaghan, R. S. Roark, K. J. Sowers, U. Nair, G. Dale, L. Maiorino, Y. Park, D. W. Kulp, B. H. Hahn, L. Shapiro, P. D. Kwong, D. R. Burton, D. J. Irvine, R. Andrabi, G. M. Shaw, F. D. Batista., Acquisition of neutralization breadth following a single priming immunization in an HIV-1 V2 apex bnAb precursor mouse model. Science Immunology, (2025).

55. B. T. Grenfell, O. G. Pybus, J. R. Gog, J. L. N. Wood, J. M. Daly, J. A. Mumford, E. C. Holmes, Unifying the epidemiological and evolutionary dynamics of pathogens. Science 303, 327–332 (2004).

56. D. T. MacLeod, N. M. Choi, B. Briney, F. Garces, L. S. Ver, E. Landais, B. Murrell, T. Wrin, W. Kilembe, C. H. Liang, A. Ramos, C. B. Bian, L. Wickramasinghe, L. Kong, K. Eren, C. Y. Wu, C. H. Wong, S. L. Kosakovsky Pond, I. A. Wilson, D. R. Burton, P. Poignard, The IAVI Protocol C. Investigators, Early antibody lineage diversification and independent limb maturation lead to broad HIV-1 neutralization targeting the Env high-mannose patch. Immunity 44, 1215–1226 (2016).

57. T. B. Kepler, H.-X. Liao, M. S. Alam, R. Bhaskarabhatla, R. Zhang, C. Yandava, S. Stewart, K. Anasti, G. Kelsoe, R. Parks, K. E. Lloyd, C. Stolarchuk, J. Pritchett, E. Solomon, E. Friberg, L. Morris, S. S. A. Karim, M. S. Cohen, E. Walter, M. A. Moody, B. F. Haynes, Immunoglobulin gene insertions and deletions in the affinity maturation of HIV-1 broadly reactive neutralizing antibodies. Cell Host & Microbe 16, 304–313 (2014).

58. K. Wiehe, T. Bradley, R. R. Meyerhoff, C. Hart, W. B. Williams, D. Easterhoff, W. J. Faison, T. B. Kepler, K. O. Saunders, S. M. Alam, M. Bonsignori, B. F. Haynes, Functional relevance of improbable antibody mutations for HIV broadly neutralizing antibody development. Cell Host & Microbe 23, 759–765.e756 (2018).

59. J. van Schooten, E. Farokhi, A. Schorcht, T. L. G. M. van den Kerkhof, H. Gao, P. v. d. Woude, J. A. Burger, T. G. R. Meesters, T. Bijl, R. Ghalaiyini, H. L. Turner, J. Dorning, B. D. C. van Schaik, A. H. C. van Kampen, C. C. Labranche, R. L. Stanfield, D. Sok, D. C. Montefiori, D. R. Burton, M. S. Seaman, G. Ozorowski, I. A. Wilson, R. W. Sanders, A. B. Ward, M. J. van Gils, Identification of IOMA-class neutralizing antibodies targeting the CD4-binding site on the HIV-1 envelope glycoprotein. Nature Communications 13, 4515 (2022).

60. R. M. Lynch, P. Wong, L. Tran, S. O’Dell, M. C. Nason, Y. Li, X. Wu, J. R. Mascola, HIV-1 fitness cost associated with escape from the VRC01 class of CD4 binding site neutralizing antibodies. Journal of Virology 89, 4201–4213 (2015).

61. J. N. Bhiman, C. Anthony, N. A. Doria-Rose, O. Karimanzira, C. A. Schramm, T. Khoza, D. Kitchin, G. Botha, J. Gorman, N. J. Garrett, S. S. A. Karim, L. Shapiro, C. Williamson, P. D. Kwong, J. R. Mascola, L. Morris, P. L. Moore, Viral variants that initiate and drive maturation of V1V2-directed HIV-1 broadly neutralizing antibodies. Nature Medicine 21, 1332–1336 (2015).

62. C. K. Wibmer, J. N. Bhiman, E. S. Gray, N. Tumba, S. S. Abdool Karim, C. Williamson, L. Morris, P. L. Moore, Viral escape from HIV-1 neutralizing antibodies drives increased plasma neutralization breadth through sequential recognition of multiple epitopes and immunotypes. PLoS Pathogens 9, e1003738 (2013).

63. F. A. Schleich, S. Bale, J. Guenaga, G. Ozorowski, M. Adori, X. Lin, X. Castro Dopico, R. Wilson, M. Chernyshev, A. T. Cotgreave, M. Mandolesi, J. Cluff, E. D. Doyle, L. M. Sewall, W. H. Lee, S. Zhang, S. O’Dell, B. S. Healy, D. Lim, V. R. Lewis, E. Ben-Akiva, D. J. Irvine, N. A. Doria-Rose, M. Corcoran, D. Carnathan, G. Silvestri, I. A. Wilson, A. B. Ward, G. B. Karlsson Hedestam, R. T. Wyatt, Vaccination of nonhuman primates elicits a broadly neutralizing antibody lineage targeting a quaternary epitope on the HIV-1 Env trimer. Immunity, 10.1016/j.immuni.2025.1004.1010 (2025).

64. R. Habib, S. O. Solieva, Z. J. Lin, S. Ghosh, K. Bayruns, M. Singh, C. J. Agostino, N. J. Tursi, K. J. Sowers, J. Huang, R. S. Roark, M. Purwar, Y. Park, K. Ayyanathan, H. Li, J. W. Carey, A. Kim, J. Park, M. E. McCanna, A. N. Skelly, N. Chokkalingam, S. Kriete, N. Shupin, A. Huynh, S. Walker, N. Laenger, J. Du, J. Cui, B. H. Hahn, A. Patel, A. Escolano, P. D. Kwong, L. Shapiro, G. R. Bowman, G. M. Shaw, D. B. Weiner, J. Pallesen, D. W. Kulp, Deep mining of the human antibody Rrepertoire identifies frequent and immunogenetically diverse CDRH3 topologies targetable by vaccination. Nature Communications, in review, (bioRxiv doi: 10.1101/2024.1110.1104.616739) (2024).

65. J. H. Lee, R. Andrabi, C. Y. Su, A. Yasmeen, J. P. Julien, L. Kong, N. C. Wu, R. McBride, D. Sok, M. Pauthner, C. A. Cottrell, T. Nieusma, C. Blattner, J. C. Paulson, P. J. Klasse, I. A. Wilson, D. R. Burton, A. B. Ward, A broadly neutralizing antibody targets the dynamic HIV Envelope trimer apex via a long, rigidified, and anionic β-hairpin structure. Immunity 46, 690–702 (2017).

66. J. R. Willis, Z. T. Berndsen, K. M. Ma, J. M. Steichen, T. Schiffner, E. Landais, A. Liguori, O. Kalyuzhniy, J. D. Allen, S. Baboo, O. Omorodion, J. K. Diedrich, X. Hu, E. Georgeson, N. Phelps, S. Eskandarzadeh, B. Groschel, M. Kubitz, Y. Adachi, T.-M. Mullin, N. B. Alavi, S. Falcone, S. Himansu, A. Carfi, I. A. Wilson, J. R. Yates, J. C. Paulson, M. Crispin, A. B. Ward, W. R. Schief, Human immunoglobulin repertoire analysis guides design of vaccine priming immunogens targeting HIV V2-apex broadly neutralizing antibody precursors. Immunity 55, 2149–2167.e2149 (2022).

67. X. Wei, S. K. Ghosh, M. E. Taylor, V. A. Johnson, E. A. Emini, P. Deutsch, J. D. Lifson, S. Bonhoeffer, M. A. Nowak, B. H. Hahn, M. S. Saag, G. M. Shaw, X. Wei, S. K. Ghosh, M. E. Taylor, V. A. Johnson, E. A. Emini, P. Deutsch, J. D. Lifson, S. Bonhoeffer, M. A. Nowak, B. H. Hahn, M. S. Saag, G. M. Shaw, Viral dynamics in human immunodeficiency virus type 1 infection. Nature 373, 117–122 (1995).

68. D. D. Ho, A. U. Neumann, A. S. Perelson, W. Chen, J. M. Leonard, M. Markowitz, D. D. Ho, A. U. Neumann, A. S. Perelson, W. Chen, J. M. Leonard, M. Markowitz, Rapid turnover of plasma virions and CD4 lymphocytes in HIV-1 infection. Nature 373, 123–126 (1995).

69. M. Markowitz, M. Louie, A. Hurley, E. Sun, M. Di Mascio, A. S. Perelson, D. D. Ho, A novel antiviral intervention results in more accurate assessment of human immunodeficiency virus type 1 replication dynamics and T-cell decay in vivo. Journal of Virology 77, 5037–5038 (2003).

70. V. Simon, D. D. Ho, HIV-1 dynamics in vivo: implications for therapy. Nature Reviews Microbiology 1, 181–190 (2003).

71. X. Wei, J. M. Decker, S. Wang, H. Hui, J. C. Kappes, X. Wu, J. F. Salazar-Gonzalez, M. G. Salazar, J. M. Kilby, M. S. Saag, N. L. Komarova, M. A. Nowak, B. H. Hahn, P. D. Kwong, G. M. Shaw, Antibody neutralization and escape by HIV-1. Nature 422, 307–312 (2003).

72. K. J. Bar, C.-y. Tsao, S. S. Iyer, J. M. Decker, Y. Yang, M. Bonsignori, X. Chen, K.-K. Hwang, D. C. Montefiori, H.-X. Liao, P. Hraber, W. Fischer, H. Li, S. Wang, S. Sterrett, B. F. Keele, V. V. Ganusov, A. S. Perelson, B. T. Korber, I. Georgiev, J. S. McLellan, J. W. Pavlicek, F. Gao, B. F. Haynes, B. H. Hahn, P. D. Kwong, G. M. Shaw, Early low-titer neutralizing antibodies impede HIV-1 replication and select for virus escape. PLoS Pathogens 8, e1002721 (2012).

73. H. Wang, C. Cheng, J. L. D. Santo, C.-H. Shen, T. Bylund, A. R. Henry, C. A. Howe, J. Hwang, N. C. Morano, D. J. Morris, S. Pletnev, R. S. Roark, T. Zhou, B. T. Hansen, F. H. Hoyt, T. S. Johnston, S. Wang, B. Zhang, D. R. Ambrozak, J. E. Becker, M. F. Bender, A. Changela, R. Chaudhary, M. Corcoran, A. R. Corrigan, K. E. Foulds, Y. Guo, M. Lee, Y. Li, B. C. Lin, T. Liu, M. K. Louder, M. Mandolesi, R. D. Mason, K. McKee, V. Nair, S. O’Dell, A. S. Olia, L. Ou, A. Pegu, N. Raju, R. Rawi, J. Roberts-Torres, E. K. Sarfo, M. Sastry, A. J. Schaub, S. D. Schmidt, C. A. Schramm, C. L. Schwartz, S. C. Smith, T. Stephens, J. Stuckey, I.-T. Teng, J.-P. Todd, Y. Tsybovsky, D. J. V. Wazer, S. Wang, N. A. Doria-Rose, E. R. Fischer, I. S. Georgiev, G. B. K. Hedestam, Z. Sheng, R. A. Woodward, D. C. Douek, R. A. Koup, T. C. Pierson, L. Shapiro, G. M. Shaw, J. R. Mascola, P. D. Kwong, Potent and broad HIV-1 neutralization in fusion peptide-primed SHIV-infected macaques. Cell 187, 7214–7231.e7223 (2024).

74. N. Goonetilleke, M. K. Liu, J. F. Salazar-Gonzalez, G. Ferrari, E. Giorgi, V. V. Ganusov, B. F. Keele, G. H. Learn, E. L. Turnbull, M. G. Salazar, K. J. Weinhold, S. Moore, CHAVI Clinical Core B, N. Letvin, B. F. Haynes, M. S. Cohen, P. Hraber, T. Bhattacharya, P. Borrow, A. S. Perelson, B. H. Hahn, G. M. Shaw, B. T. Korber, A. J. McMichael, The first T cell response to transmitted/founder virus contributes to the control of acute viremia in HIV-1 infection. The Journal of Experimental Medicine 206, 1253–1272 (2009).

75. J. F. Salazar-Gonzalez, M. G. Salazar, B. F. Keele, G. H. Learn, E. E. Giorgi, H. Li, J. M. Decker, S. Wang, J. Baalwa, M. H. Kraus, N. F. Parrish, K. S. Shaw, M. B. Guffey, K. J. Bar, K. L. Davis, C. Ochsenbauer-Jambor, J. C. Kappes, M. S. Saag, M. S. Cohen, J. Mulenga, C. A. Derdeyn, S. Allen, E. Hunter, M. Markowitz, P. Hraber, A. S. Perelson, T. Bhattacharya, B. F. Haynes, B. T. Korber, B. H. Hahn, G. M. Shaw, Genetic identity, biological phenotype, and evolutionary pathways of transmitted/founder viruses in acute and early HIV-1 infection. The Journal of Experimental Medicine 206, 1273–1289 (2009).

76. H. Song, J. W. Pavlicek, F. Cai, T. Bhattacharya, H. Li, S. S. Iyer, K. J. Bar, J. M. Decker, N. Goonetilleke, M. K. Liu, A. Berg, B. Hora, M. S. Drinker, J. Eudailey, J. Pickeral, M. A. Moody, G. Ferrari, A. McMichael, A. S. Perelson, G. M. Shaw, B. H. Hahn, B. F. Haynes, F. Gao, Impact of immune escape mutations on HIV-1 fitness in the context of the cognate transmitted/founder genome. Retrovirology 9, 89 (2012).

77. K. M. Ma, H. J. Sutton, P. P. Pratap, J. M. Steichen, D. Carnathan, J. Quinn, O. Kalyuzhniy, A. Liguori, S. Agrawal, S. Baboo, P. Madden, C. A. Cottrell, J. R. Willis, J. H. Lee, E. Landais, X. Hu, P. Ramezani-Rad, G. Ozorowski, V. R. Lewis, J. K. Diedrich, X. Zhou, T. K. Altheide, N. Phelps, E. Georgeson, N. B. Alavi, D. Lu, S. Eskandarzadeh, M. Kubitz, Y. Adachi, T. M. Mullen, M. Silva, M. B. Melo, S. Himansu, D. J. Irvine, D. R. Burton, J. R. Yates, 3rd, J. C. Paulson, D. Sok, I. A. Wilson, G. Silvestri, A. B. Ward, S. Crotty, W. R. Schief, HIV broadly neutralizing antibody precursors to the Apex epitope induced in nonhuman primates. Sci Immunol 10, eadt6660 (2025).

78. N. Mishra, B. Liang, R. S. Roark, A. Ghosh, S. Callaghan, W. H. Lee, X. Li, A. L. Vo, G. Avillion, R. R. Chowdhury, R. Habib, F. Bibollet-Ruche, G. Giese, P. Oberoi, K. Amereh, A. Somanathan, Y. Zhou, Y. Zhang, M. Kassab, L. Tijo, S. Andrabi, R. A. Reyes, J. D. Allen, N. E. James, K. N. R. Jr, L. van der Maas, E. Ben-Akiva, K. Kacmarek-Michaels, S. Plante, C. L. Martella, A. N. Skelly, A. Singh, J. Hurtado, K. Dueker, T. Capozzola, R. Nedellec, G. Ozorowski, M. M. Lewis, S. Falcone, A. Carfi, S. Himansu, L. Shapiro, M. Crispin, B. H. Hahn, B. Briney, D. J. Irvine, D. R. Burton, A. B. Ward, F. D. Batista, P. D. Kwong, G. M. Shaw, R. Andrabi, Germline-targeting HIV immunogen induces cross-neutralizing antibodies in outbred macaques. Immunity (in press), (2025).

79. , (!!! INVALID CITATION !!! (21, 30, 46, 60-62, 79)).

80. V. Dubrovskaya, J. Guenaga, N. d. Val, R. Wilson, Y. Feng, A. Movsesyan, G. B. K. Hedestam, A. B. Ward, R. T. Wyatt, Targeted N-glycan deletion at the receptor-binding site retains HIV Env NFL trimer integrity and accelerates the elicited antibody response. PLoS Pathogens 13, e1006614 (2017).

81. T. Zhou, N. A. Doria-Rose, C. Cheng, G. B. E. Stewart-Jones, G.-Y. Chuang, M. Chambers, A. Druz, H. Geng, K. McKee, Y. D. Kwon, S. O’Dell, M. Sastry, S. D. Schmidt, K. Xu, L. Chen, R. E. Chen, M. K. Louder, M. Pancera, T. G. Wanninger, B. Zhang, P. D. Kwong, Quantification of the impact of the HIV-1-glycan shield on antibody elicitation. Cell Reports 19, 719–732 (2017).

82. E. T. Crooks, K. Osawa, T. Tong, S. L. Grimley, Y. D. Dai, R. G. Whalen, D. W. Kulp, S. Menis, W. R. Schief, J. M. Binley, Effects of partially dismantling the CD4 binding site glycan fence of HIV-1 Envelope glycoprotein trimers on neutralizing antibody induction. Virology 505, (2017).

83. A. N. Skelly, H. B. Gristick, H. Li, E. Gavor, A. J. Connell, E. F. Kreider, M. P. Hogarty, L. Marchitto, M. Newby, J. D. Allen, W. Liu, A. P. W. Jr, M. S. Campion, K. Winters, C. G. Gordon, R. A. Osbaldeston, M. J. Akeley, K. Ayyanathan, Y. Li, A. Singh, K. Cruickshank, Y. Park, C. Zhao, X. Li, E. Van Itallie, J. W. Carey, A. Albertus, A. T. DeLaitsch, J. R. Keeffe, R. Habib, D. J. Morris, F. Bibollet-Ruche, N. Koranda, S. J. Plante, C. Martella, J. Lora, E. J. D. Wang, M. G. Lituchy, M. G. Lewis, K. J. Wiehe, K. Wagh, B. Korber, B. F. Haynes, M. A. Martin, M. C. Nussenzweig, M. S. Seaman, D. J. Irvine, R. Andrabi, M. Crispin, D. Weissman, P. J. Bjorkman, B. H. Hahn, G. M. Shaw, Rapid induction of HIV-1 V3 glycan broadly-neutralizing antibodies by a novel two-step priming mechanism guides immunogen design. bioRxiv, (2025).

84. J. Guenaga, M. Adori, S. Bale, S. Phulera, I. Zygouras, S. Agrawal, F.-A. Schleich, M. Ota, R. Wilson, J. Cluff, T. Dzvelaia, M. Mandolesi, L. Verkoczy, D. J. Irvine, M. Corcoran, I. A. Wilson, D. Carnathan, G. Silvestri, A. B. Ward, G. Ozorowski, G. B. K. Hedestam, R. T. Wyatt, HIV Env trimers elicit NHP apex cross-neutralizing antibodies mimicking human bNAbs. bioRxiv, (2025).

85. J. M. Steichen, I. Phung, E. Salcedo, G. Ozorowski, J. R. Willis, S. Baboo, A. Liguori, C. A. Cottrell, J. L. Torres, P. J. Madden, K. M. Ma, H. J. Sutton, J. H. Lee, O. Kalyuzhniy, J. D. Allen, O. L. Rodriguez, Y. Adachi, T.-M. Mullen, E. Georgeson, M. Kubitz, A. Burns, S. Barman, R. Mopuri, A. Metz, T. K. Altheide, J. K. Diedrich, S. Saha, K. Shields, S. E. Schultze, M. L. Smith, T. Schiffner, D. R. Burton, C. T. Watson, S. E. Bosinger, M. Crispin, J. R. YatesIII, J. C. Paulson, A. B. Ward, D. Sok, S. Crotty, W. R. Schief, Vaccine priming of rare HIV broadly neutralizing antibody precursors in nonhuman primates. Science 384, (2024).

86. L. M. Walker, S. K. Phogat, P.-Y. Chan-Hui, D. Wagner, P. Phung, J. L. Goss, T. Wrin, M. D. Simek, S. Fling, J. L. Mitcham, J. K. Lehrman, F. H. Priddy, O. A. Olsen, S. M. Frey, P. W. Hammond, Protocol G Principal Investigators, S. Kaminsky, T. Zamb, M. Moyle, W. C. Koff, P. Poignard, D. R. Burton, Broad and potent neutralizing antibodies from an African donor reveal a new HIV-1 vaccine target. Science 326, 285–289 (2009).

87. M. D. Simek, W. Rida, F. H. Priddy, P. Pung, E. Carrow, D. S. Laufer, J. K. Lehrman, M. Boaz, T. Tarragona-Fiol, G. Miiro, J. Birungi, A. Pozniak, D. A. McPhee, O. Manigart, E. Karita, A. Inwoley, W. Jaoko, J. DeHovitz, L.-G. Bekker, P. Pitisuttithum, R. Paris, L. M. Walker, P. Poignard, T. Wrin, P. E. Fast, D. R. Burton, W. C. Koff, Human immunodeficiency virus type 1 elite neutralizers: individuals with broad and potent neutralizing activity identified by using a high-throughput neutralization assay together with an analytical selection algorithm. Journal of Virology 83, 7337–7348 (2009).

88. M. A. Moody, I. Pedroza-Pacheco, N. A. Vandergrift, C. Chui, K. E. Lloyd, R. Parks, K. A. Soderberg, A. T. Ogbe, M. S. Cohen, H.-X. Liao, F. Gao, A. J. McMichael, D. C. Montefiori, L. Verkoczy, G. Kelsoe, J. Huang, P. R. Shea, M. Connors, P. Borrow, B. F. Haynes, Immune perturbations in HIV-1–infected individuals who make broadly neutralizing antibodies. Science Immunology 1, aag0851 (2016).

89. S. J. Krebs, Y. D. Kwon, C. A. Schramm, W. H. Law, G. Donofrio, K. H. Zhou, S. Gift, V. Dussupt, I. S. Georgiev, S. Schätzle, J. R. McDaniel, Y. T. Lai, M. Sastry, B. Zhang, M. C. Jarosinski, A. Ransier, A. L. Chenine, M. Asokan, R. T. Bailer, M. Bose, A. Cagigi, E. M. Cale, G. Y. Chuang, S. Darko, J. I. Driscoll, A. Druz, J. Gorman, F. Laboune, M. K. Louder, K. McKee, L. Mendez, M. A. Moody, A. M. O’Sullivan, C. Owen, D. Peng, R. Rawi, E. Sanders-Buell, C. H. Shen, A. R. Shiakolas, T. Stephens, Y. Tsybovsky, C. Tucker, R. Verardi, K. Wang, J. Zhou, T. Zhou, G. Georgiou, S. M. Alam, B. F. Haynes, M. Rolland, G. R. Matyas, V. R. Polonis, A. B. McDermott, D. C. Douek, L. Shapiro, S. Tovanabutra, N. L. Michael, J. R. Mascola, M. L. Robb, P. D. Kwong, N. A. Doria-Rose, Longitudinal analysis reveals early development of three MPER-directed neutralizing antibody lineages from an HIV-1-infected individual. Immunity 50, 677–691.e613 (2019).

90. V. Bhardwaj, M. Franceschetti, R. Rao, P. A. Pevzner, Y. Safonova, Automated analysis of immunosequencing datasets reveals novel immunoglobulin D genes across diverse species. PLOS Computational Biology 16, e1007837 (2020).

91. N. Vázquez Bernat, M. Corcoran, I. Nowak, M. Kaduk, X. Castro Dopico, S. Narang, P. Maisonasse, N. Dereuddre-Bosquet, B. Murrell, G. B. Karlsson Hedestam, Rhesus and cynomolgus macaque immunoglobulin heavy-chain genotyping yields comprehensive databases of germline VDJ alleles. Immunity 54, 355–366.e354 (2021).

92. A. Ramesh, S. Darko, A. Hua, G. Overman, A. Ransier, J. R. Francica, A. Trama, G. D. Tomaras, B. F. Haynes, D. C. Douek, T. B. Kepler, Structure and diversity of the rhesus macaque immunoglobulin loci through multiple de novo genome assemblies. Frontiers in Immunology 8, 1407 (2017).

93. F. Breden, E. T. Luning Prak, B. Peters, F. Rubelt, C. A. Schramm, C. E. Busse, J. A. Vander Heiden, S. Christley, S. A. C. Bukhari, A. Thorogood, F. A. Matsen IV, Y. Wine, U. Laserson, D. Klatzmann, D. C. Douek, M.-P. Lefranc, A. M. Collins, T. Bubela, S. H. Kleinstein, C. T. Watson, L. G. Cowell, J. K. Scott, T. B. Kepler, Reproducibility and reuse of adaptive immune receptor repertoire data. Frontiers in Immunology 8, 1418 (2017).

94. M. Martin, Cutadapt removes adapter sequences from high-throughput sequencing reads. EMBnet Journal 17, 10–12 (2011).

95. J. Zhang, K. Kobert, T. Flouri, A. Stamatakis, PEAR: a fast and accurate Illumina Paired-End reAd mergeR. Bioinformatics 30, 614–620 (2014).

96. F. Mölder, K. P. Jablonski, B. Letcher, M. B. Hall, C. H. Tomkins-Tinch, V. Sochat, J. Forster, S. Lee, S. O. Twardziok, A. Kanitz, A. Wilm, M. Holtgrewe, S. Rahmann, S. Nahnsen, J. Köster, F. Mölder, K. P. Jablonski, B. Letcher, M. B. Hall, C. H. Tomkins-Tinch, V. Sochat, J. Forster, S. Lee, S. O. Twardziok, A. Kanitz, A. Wilm, M. Holtgrewe, S. Rahmann, S. Nahnsen, J. Köster, Sustainable data analysis with Snakemake. F1000Research 10, 33 (2021).

97. D. Trudgian, G. M. Kurtzer, cclerget, M. Bauer, I. Kaneshiro, D. Godlove, Vanessasaurus, Y. Cote, C. E. A. Gutierrez, G. Vallee, DrDaveD, J. Cook, A. Hughes, J. Stover, B. P. Bockelman, M. Magallon, J. Chappell, M. Frisch, D. Tamino, C. Madison, S. Yakovtseva, A. Duffy, S. Ghosh, VP, T. Huynh, M. Gray, Y. Halchenko, F. Abecassis, sylabs/singularity: SingularityCE 4.3.1. Zenodo, (2015).

98. A. Punjani, J. L. Rubinstein, D. J. Fleet, M. A. Brubaker, A. Punjani, J. L. Rubinstein, D. J. Fleet, M. A. Brubaker, cryoSPARC: algorithms for rapid unsupervised cryo-EM structure determination. Nature Methods 14, 290–296 (2017).

99. E. F. Pettersen, T. D. Goddard, C. C. Huang, E. C. Meng, G. S. Couch, T. I. Croll, J. H. Morris, T. E. Ferrin, UCSF ChimeraX: Structure visualization for researchers, educators, and developers. Protein Science 30, 70–82 (2021).

100. P. Emsley, K. Cowtan, Coot: model-building tools for molecular graphics. Acta Crystallographica Section D 60, 2126–2132 (2004).

101. P. D. Adams, K. Gopal, R. W. Grosse-Kunstleve, L.-W. Hung, T. R. Ioerger, A. J. McCoy, N. W. Moriarty, R. K. Pai, R. J. Read, T. D. Romo, J. C. Sacchettini, N. K. Sauter, L. C. Storoni, T. C. Terwilliger, Recent developments in the PHENIX software for automated crystallographic structure determination. Journal of Synchrotron Radiation 11, 53–55 (2004).

102. I. W. Davis, L. W. Murray, J. S. Richardson, D. C. Richardson, MolProbity: structure validation and all-atom contact analysis for nucleic acids and their complexes. Nucleic Acids Research 35, W375–W383 (2004).

103. B. A. Barad, N. Echols, R. Y.-R. Wang, Y. Cheng, F. DiMaio, P. D. Adams, J. S. Fraser, B. A. Barad, N. Echols, R. Y.-R. Wang, Y. Cheng, F. DiMaio, P. D. Adams, J. S. Fraser, EMRinger: side chain–directed model and map validation for 3D cryo-electron microscopy. Nature Methods 12, 943–946 (2015).

104. C. A. Bricault, K. Yusim, M. S. Seaman, H. Yoon, J. Theiler, E. E. Giorgi, K. Wagh, M. Theiler, P. Hraber, J. P. Macke, E. F. Kreider, G. H. Learn, B. H. Hahn, J. F. Scheid, J. M. Kovacs, J. L. Shields, C. L. Lavine, F. Ghantous, M. Rist, M. G. Bayne, G. H. Neubauer, K. McMahan, H. Peng, C. Chéneau, J. J. Jones, J. Zeng, C. Ochsenbauer, J. P. Nkolola, K. E. Stephenson, B. Chen, S. Gnanakaran, M. Bonsignori, L. D. Williams, B. F. Haynes, N. Doria-Rose, J. R. Mascola, D. C. Montefiori, D. H. Barouch, B. Korber, HIV-1 neutralizing antibody signatures and application to epitope-targeted vaccine design. Cell Host & Microbe 25, 59–72.e58 (2019).

105. A. Changela, X. Wu, Y. Yang, B. Zhang, J. Zhu, G. A. Nardone, S. O’Dell, M. Pancera, M. K. Gorny, S. Phogat, J. E. Robinson, L. Stamatatos, S. Zolla-Pazner, J. R. Mascola, P. D. Kwong, Crystal structure of human antibody 2909 reveals conserved features of quaternary structure-specific antibodies that potently neutralize HIV-1. Journal of Virology 85, 2524–2535. (2010).

