## Supplemental Figures and Text for "Env-antibody coevolution identifies B cell priming as the principal bottleneck to HIV-1 V2 apex broadly neutralizing antibody development"

**Supplemental Text**

***Monte Carlo random sampling analysis of bNAb elicitation rates***

For each trial in the Monte-Carlo experiments, the 18 positives for V2 apex and 4 positives for V3-glycan were randomly distributed among the 115 monkeys, with each RM keeping its assigned SHIV. SHIVs were sorted according to the fraction of bNAb positive monkeys associated with that SHIV. The top *n* SHIVs were ranked by the fraction positive, and then a statistic computed for the sum of positives among the top *n* SHIVs. Based on 100,000 trials, the below estimated p-values for different values of n were found for the V2 data:

| n | p |
|---|---|
| **1** | 0.03297 |
| **2** | 0.08272 |
| **3** | 0.00073 |
| **4** | 0.00074 |
| **5** | 0.00006 |

Applied to the V3-glycan data:

| n | p |
|---|---|
| **1** | 0.18157 |
| **2** | 0.04934 |

| | |
|---|---|
| **3** | 0.41074 |
| **4** | 1.0 |
| **5** | 1.0 |

These results indicate the restriction of both V2 apex and V3-glycan bNAbers to five and two SHIVs respectively to be significant.

***Mapping of early plasma responses in RM V033***

Figure **S6** shows sequential Env SGS sequences from RM V033 spanning weeks 0-56 post-infection. At week 8 post-infection, SGS revealed striking positive selection in Env at four sites: an N187S substitution that introduced a glycan to the V2 hypervariable loop at residue 185, an R308H mutation on the V3 loop, several mutations clustered around K460 (K460E, D461G/N, N463H, V464E) at the V5 loop, and a T533A mutation. These mutations were followed at week 12 by additional mutations at Q170R and K171R on the V2 apex C-strand (**Fig. 3B, S6**). To understand which of these mutations constituted escape from NAb lineages, we constructed a series of mutants that incorporated these mutations individually, as well as week 8 and 12 consensus viruses bearing 3 and 5 mutations respectively. All apical mutations (those at residues 170, 171, and 187) as well as R308H conferred decreases in plasma neutralization potency, indicating these to be escape mutations from NAb responses (**fig. S6**). On the other hand, the K460E mutation resulted in no change in neutralization suggesting that this escape arose to escape cytotoxic T cell responses (**fig. S6**). The T533A substitution was not necessary for abrogation of neutralization by week 8 or 12 plasmas, suggesting that it did not arise to escape Nab responses.

Furthermore, the same mutation was identified in all SHIV-Q23.17 infected RMs at the earliest timepoint sequenced (**fig. S27**). A533 is a highly conserved residue, present in >99% of the 10,000+ global HIV-1 Envs in the LANL database (https://www.hiv.lanl.gov). This led us to determine that this mutation was a fitness reversion, rather than an immune escape variant. Overall, these results indicate that the initial immune response in RM V033 consisted of NAbs targeting two epitopes, the V2 apex and linear V3 loop, as well as V5 loop directed cytotoxic T cell responses.

***Temporal dynamics of Env escape and breadth acquisition in RM V033***

To understand how Env-bNAb coevolution led to breadth development in RM V033, we examined the relationship between Env escape mutations and heterologous virus neutralization. We found that the temporal emergence of within-host V2 apex escape mutations could explain the dynamics of serum breadth acquisition in this macaque. While the Q23.17 T/F Env lacked glycans in the V2 hypervariable loop, the Env quasispecies evolved to acquire a potential N-linked glycan site at 185 through an N187S substitution first detectable at week 4 (**Fig. 3B**). This mutation dominated the quasispecies at week 8 (92%, 34 out of 37) and reached fixation from week 12 onwards. While week 12 serum could neutralize 3 heterologous viruses with one or two glycans in hypervariable V2 loop, week 4 and 8 sera could not, thus indicating that N187S arose to dominance just prior to detection of serum neutralization of heterologous Envs with V2 loop glycans (**Fig. 3A**). A similar trend was also seen for Q170R that dominates the quasispecies beginning week 16 (61%, 17 of 28), preceding the neutralization of R170 containing heterologous viruses MT145K (by week 20 serum, 4 weeks later) and ZM233.6 (by week 24 serum, 8 weeks later). These results suggest that serum neutralization of heterologous viruses carrying certain V2 apex amino acids could only be detected after those Env features were selected for in the within-host Env quasispecies. N187S provides neutralization escape from the first bNAb intermediate I1 but is sensitive to I2, indicating this mutation was responsible for selecting for maturation from I1 to I2 (**Fig. 3D**). Similarly, Q170R likely was involved in maturation from I6 to the fully mature V033-a.01. These findings suggest a scenario in which within-host Env escape mutations from previous bNAb intermediates select for more matured bNAb lineage members that can neutralize those escape mutations in heterologous viruses, and it is this capacity that confers increasing breadth acquisition as the V033 bNAb lineage matures. Similar patterns were also detected for the other SHIV Q23.17 infected V2 bNAber V031, in which the mutations H130N (which introduces a glycan at N130) and R169K are first detected at week 32 in longitudinal Envs 16 weeks prior to the first detection of serum neutralization of heterologous Envs with either of these amino acids, observed beginning week 48 (**Fig. 5B, Table S1**).

***Sensitivity and specificity analysis of C-strand selection in SHIV-infected RMs***

Out of the 115 RMs in this study, 18 were found to have V2 apex bNAbs (**Fig. 1A**), and five were found to have C-strand targeted NAbs responses with limited heterologous neutralization that did not meet our criteria for breadth (**fig. S22**). Nineteen out of 23 RMs with C-strand targeted NAbs showed complete replacement of the WT C-strand, while 23/23 showed at least 50% replacement (**Fig. 5B and fig. S17**). These results established 50% C-strand replacement to have a sensitivity of 100% as an indicator of the presence of a C-strand targeted NAb. To probe the specificity of C-strand selection, we looked for the frequency of C-strand selection among RMs that did not make C-strand targeted NAbs. No RMs infected with SHIVs CH694, 40100, WITO, 191859, CH1012, or BG505 achieved 50% replacement of the C-strand (**Fig. 6B and fig. S21**). Two out of three

RMs infected with SHIV-ZM233 showed no C-strand selection (**fig. S21)**. Amongst SHIVs that reproducibly elicited V2 apex bNAbs — Q23.17, CAP256SU, T250, Ce1176, and CH505 — C-strand selection was absent in the majority of RMs that did not develop C-strand-targeted responses, indicating this selection was specifically the result of antibody escape (**Fig. 6B and fig. S21**). Only one RM, 6927 (SHIV-T250), showed over 50% C-strand replacement that was not associated with a C-strand targeted NAb response.

SHIVs B41 and CH848, and Ce1086 represented outliers. Six out of six SHIV-B41 infected RMs showed rapid C-strand selection as early as week 12 post-infection (**fig. S21 and S24**), although this selection only reached 50% replacement in three out of six RMs. Three out of six RMs infected with SHIV-CH848 acquired an E169K reversion to group M consensus also detectable in the initial trial participant CH848 (**Fig. S23**). However, this mutation is expected to sensitize the V2 apex to neutralization by V2 apex bNAbs (*104*), suggesting that it was acquired to increase viral replicative fitness rather than as escape from an antibody lineage. C-strand selection in SHIV Ce1086-infected RMs was transient, reverted to the T/F sequence following acquisition of a K160N mutation introducing a glycan at residue 160, and only one RM (T683) reached 50% C-strand replacement (**fig. S28**). Genetic linkage between acquisition of the N160 glycan and C-strand selection suggest that C-strand escape in those RMs was the result of a 2909-like N160 glycan hole-specific antibody response (*50, 105*). Despite the frequency of selection, none of the SHIV-B41, Ce1086, or CH848 infected RMs showed any evidence of V2 apex-directed breadth.

Overall, 17/23 RMs with C-strand targeted responses showed complete T/F loss at the C-strand, while 23/23 RMs had at least 50% replacement of the WT C-strand. Analysis of RMs without V2 apex-directed NAbs yielded a final number of 3/78 RMs showing complete replacement of the WT C-strand, and 8/78 showing at least 50% T/F loss. These results identify 50% C-strand replacement as a highly significant, sensitive, and specific indicator of C-strand targeted responses ($p < 0.0001$, Fisher's Exact Test; Sensitivity = 1.000; Specificity = 0.897).

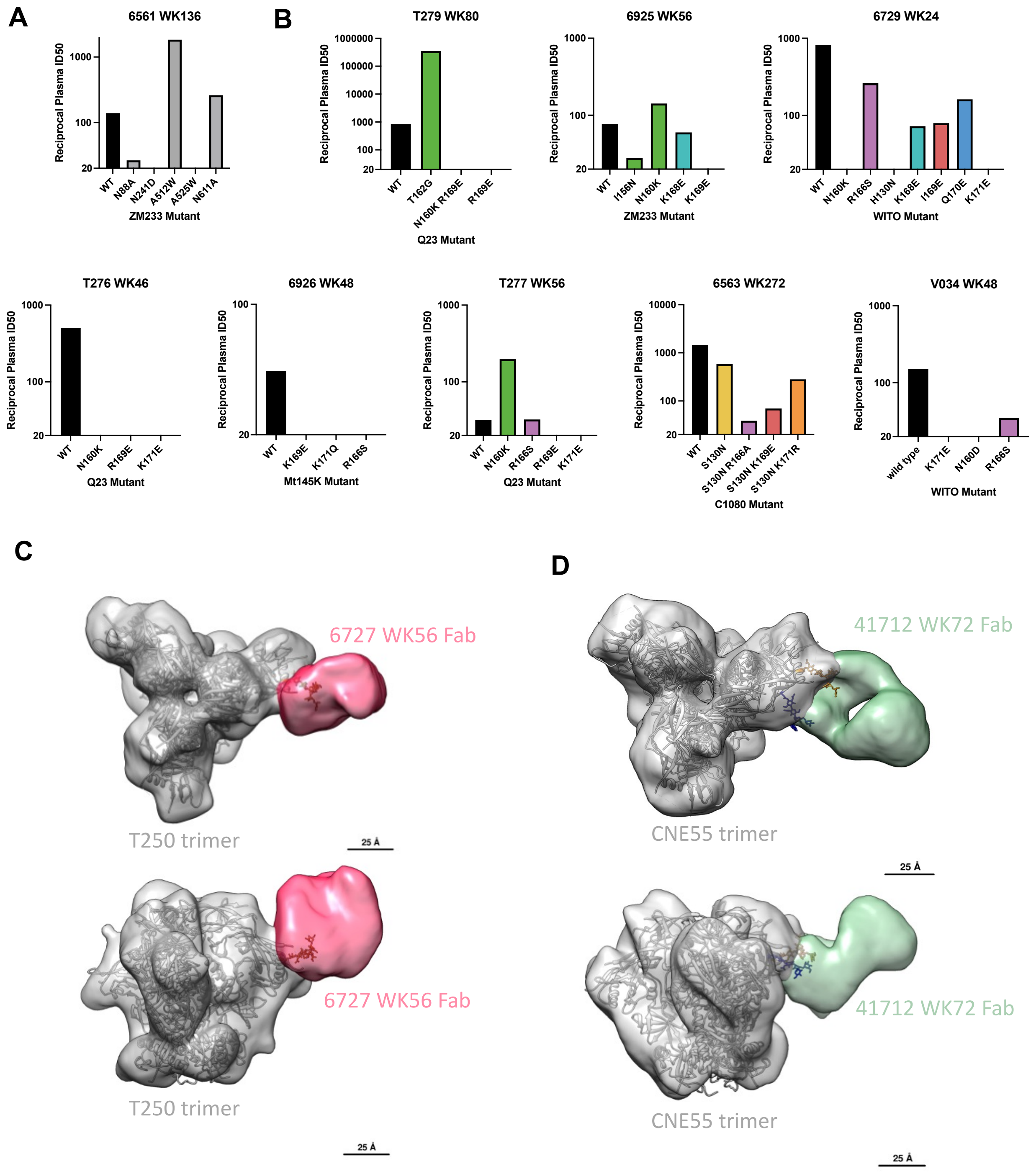


**Fig. S1. Epitope specificity of heterologous neutralization in SHIV-infected RMs.** (**A**) Epitope mapping against heterologous wild type and FP-epitope mutant viruses by plasma from RM6561. (**B**) Epitope mapping against heterologous wild type and V2 apex-epitope mutant viruses by plasma from eight RMs. (**C-D**) Negative-stain electron microscopy polyclonal epitope mapping (NS-EMPEM) of polyclonal Fabs from RMs with silent face bNAbs 6727 (**C**) and 41712 (**D**) against soluble heterologous Env trimers.

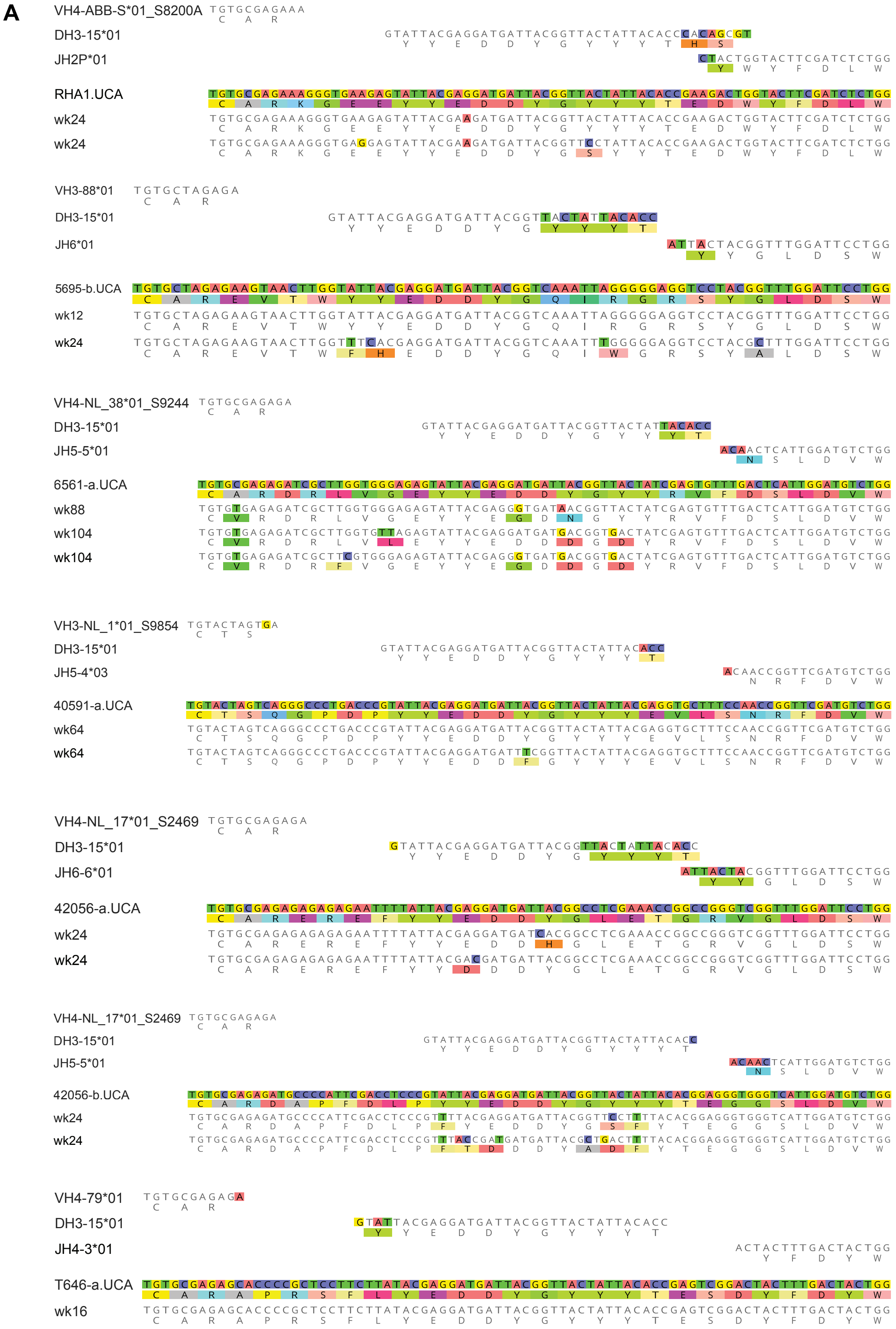
A
VH4-ABB-S*01_S8200A
DH3-15*01
JH2P*01
RHA1.UCA
wk24
wk24
VH3-88*01
DH3-15*01
JH6*01
5695-b.UCA
wk12
wk24
VH4-NL_38*01_S9244
DH3-15*01
JH5-5*01
6561-a.UCA
wk88
wk104
wk104
VH3-NL_1*01_S9854
DH3-15*01
JH5-4*03
40591-a.UCA
wk64
wk64
VH4-NL_17*01_S2469
DH3-15*01
JH6-6*01
42056-a.UCA
wk24
wk24
VH4-NL_17*01_S2469
DH3-15*01
JH5-5*01
42056-b.UCA
wk24
wk24
VH4-79*01
DH3-15*01
JH4-3*01
T646-a.UCA
wk16

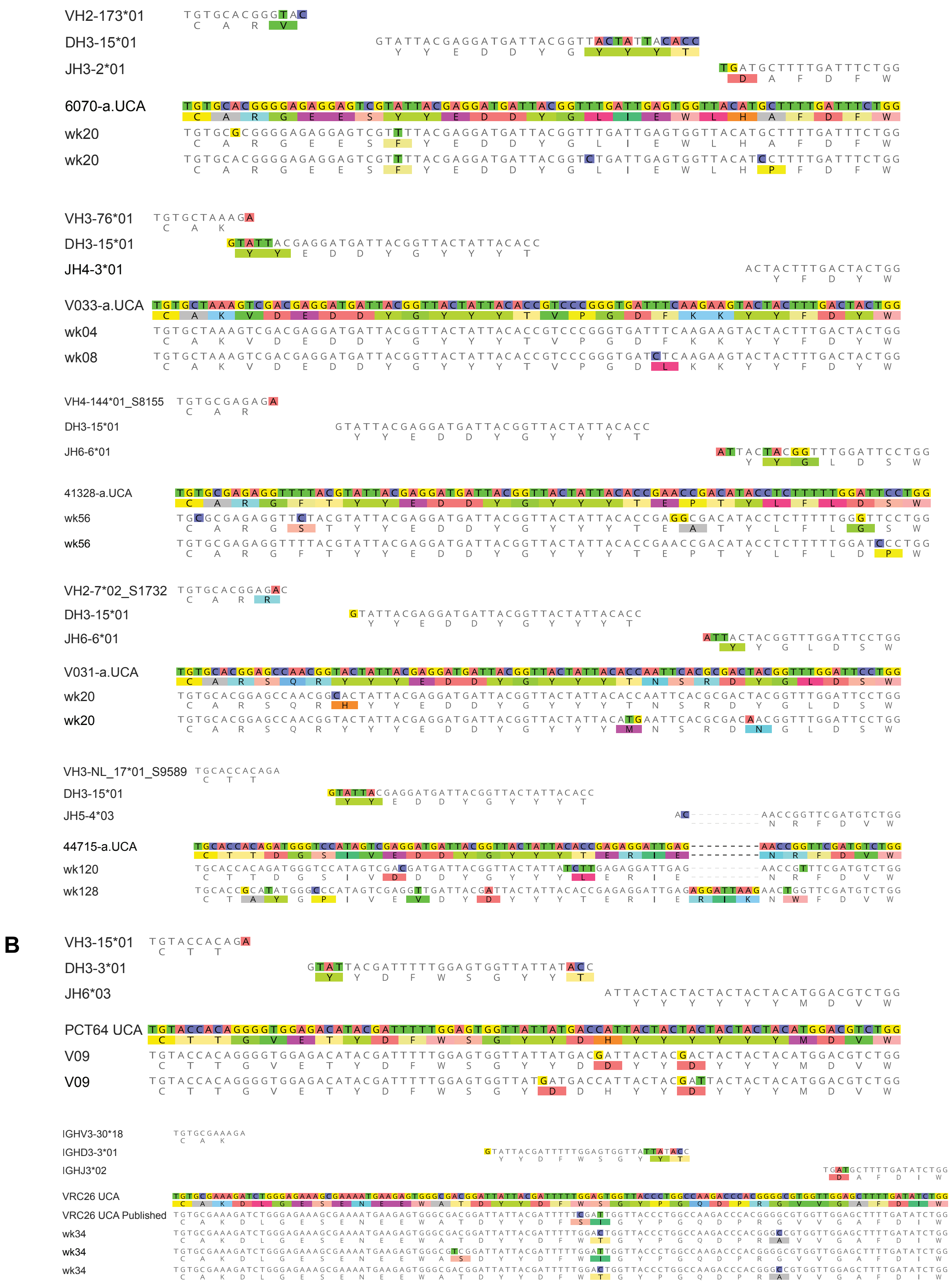
VH2-173*01
DH3-15*01
JH3-2*01
6070-a.UCA
wk20
wk20
VH3-76*01
DH3-15*01
JH4-3*01
V033-a.UCA
wk04
wk08
VH4-144*01_S8155
DH3-15*01
JH6-6*01
41328-a.UCA
wk56
wk56
VH2-7*02_S1732
DH3-15*01
JH6-6*01
V031-a.UCA
wk20
wk20
VH3-NL_17*01_S9589
DH3-15*01
JH5-4*03
44715-a.UCA
wk120
wk128
B
VH3-15*01
DH3-3*01
JH6*03
PCT64 UCA
V09
V09
IGHV3-30*18
IGHD3-3*01
IGHJ3*02
VRC26 UCA
VRC26 UCA Published
wk34
wk34
wk34

**Fig. S2. Inference of V2 apex bNAb UCAs.** Nucleotide and translation alignments of (**A**) newly inferred rhesus and (**B**) re-inferred human V2 apex bNAb UCAs with assigned VDJ genes and early B cell NGS-derived lineage members. Mismatches to the UCA are highlighted. Data in panel B are re-analyzed from previous studies (*23, 46*). Both previously published and revised inferences of the VRC26 UCA are shown.

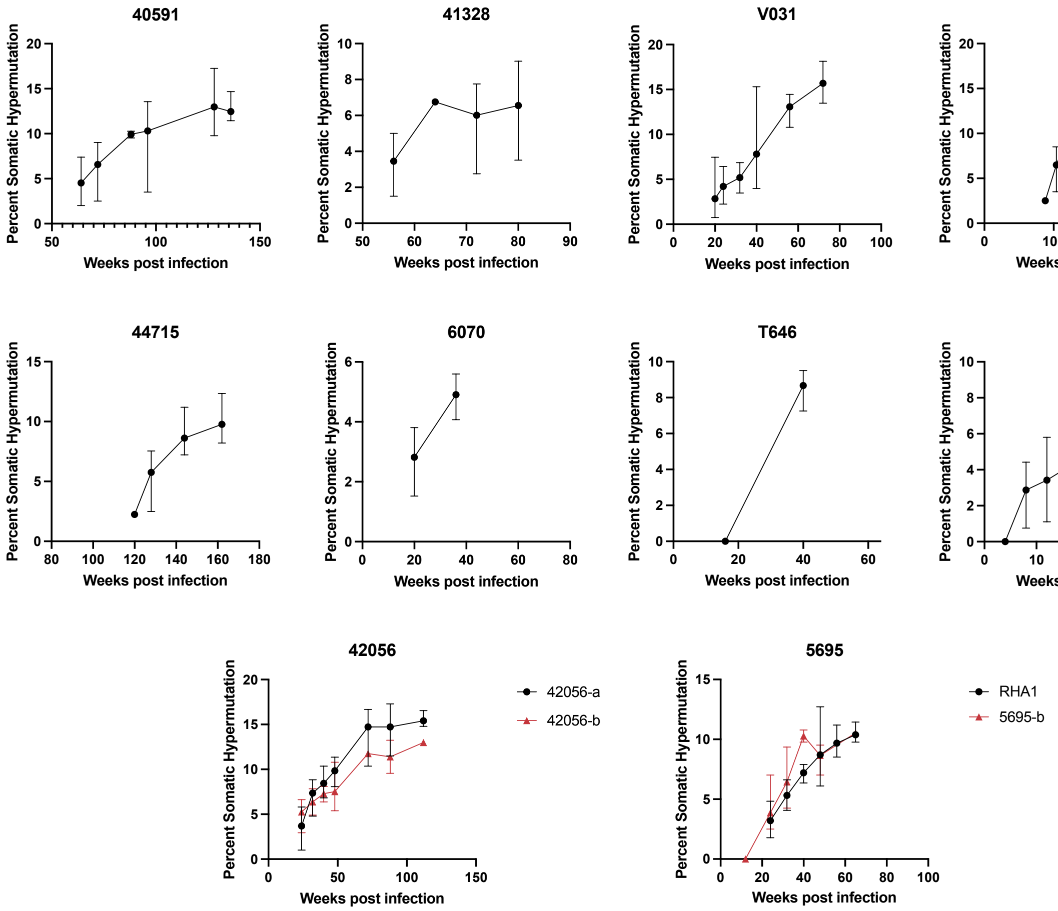


**Fig. S3. Longitudinal somatic hypermutation of V2 apex bNAb lineages.** Percent somatic hypermutation of rhesus V2 apex bNAb lineages members at timepoints where lineages were detectable by B cell NGS**.** Error bars represent the range of values.

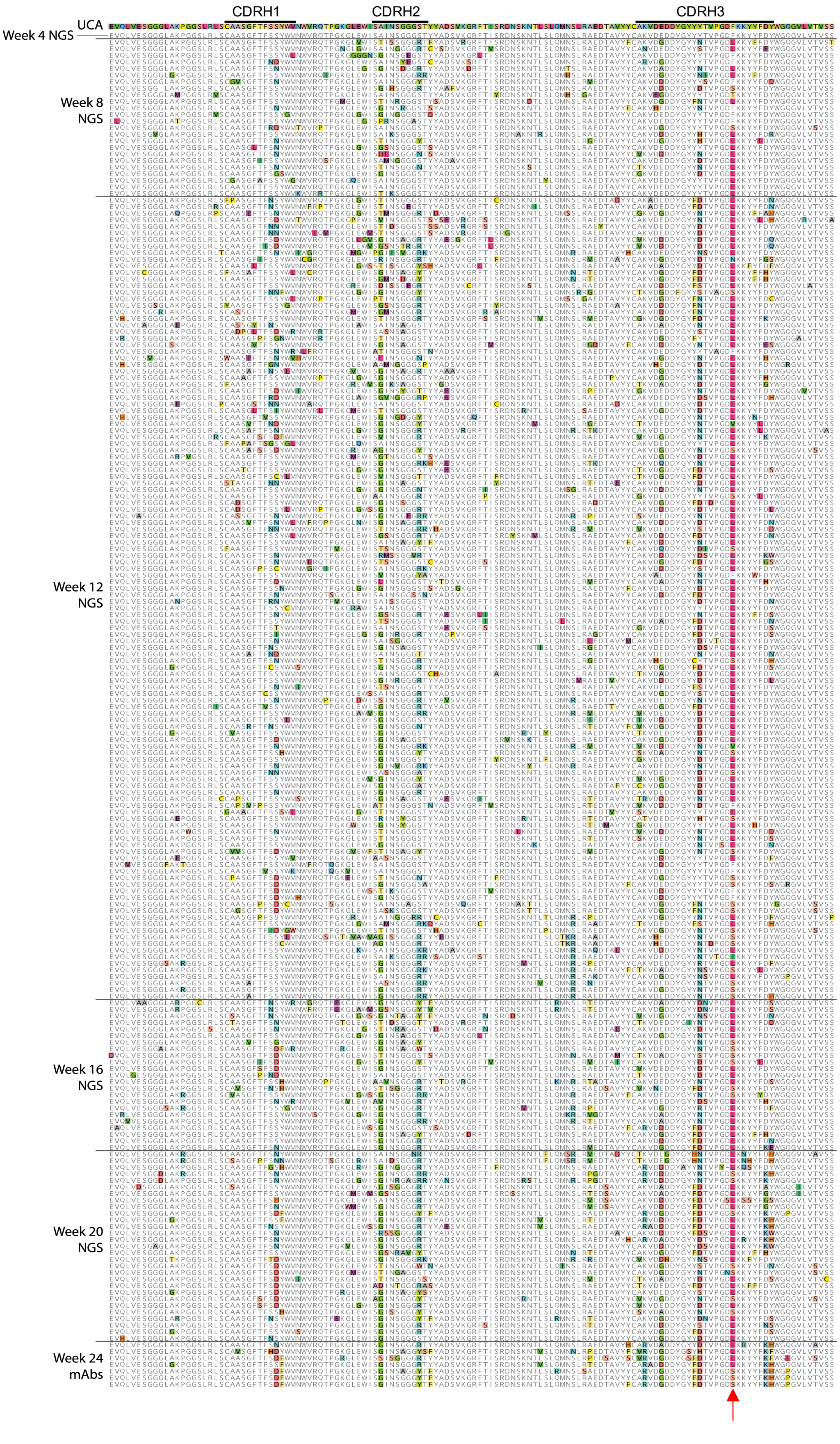
CDRH1
CDRH2
CDRH3
UCA
Week 4 NGS
Week 8 NGS
Week 12 NGS
Week 16 NGS
Week 20 NGS
Week 24 mAbs

**Fig. S4. Longitudinal V033-a lineage heavy chain sequences.** Sequence alignment of B cell NGS-derived V033-a lineage members from weeks 4, 8, and 12 post-infection, and mature bNAb sequences isolated at week 24 post-infection. Mismatches to the UCA are highlighted. Residue 100j is indicated by a red arrow.

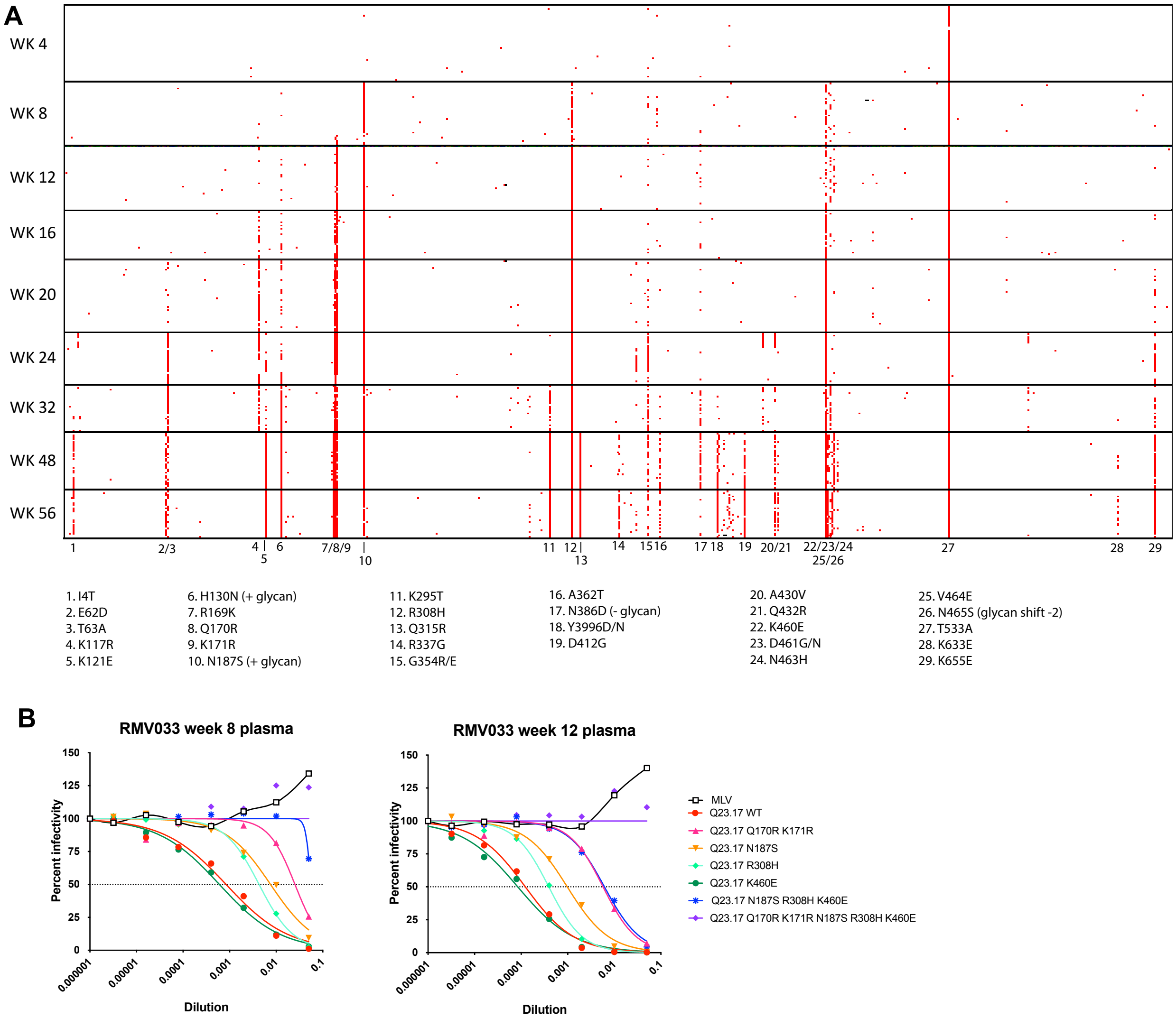


**Fig. S5. Env-Antibody Coevolution in RM V033.** (**A**) A pixel plot of longitudinal Env sequences from RM V033. Mismatches to the T/F sequence are colored red, and major mutations are indicated below. (**B**) Neutralization of wild type and mutant Q23.17 pseudoviruses by plasma from RM V033 at weeks 8 and 12 post-infection. Q23.17 pseudoviruses contained mutations identified from week 8 and week 12 SGS sequences.

A

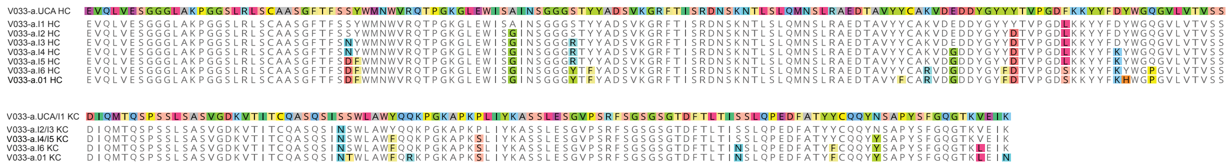


B

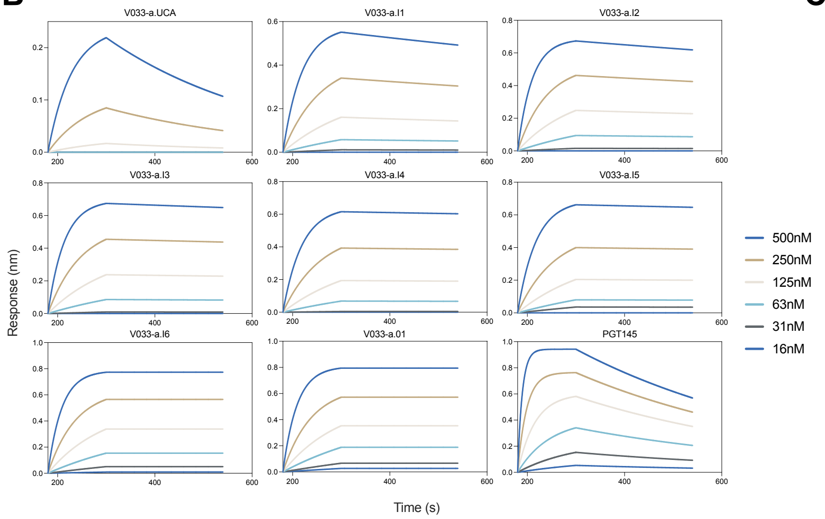


C

| | KD (M) |
|---|---|
| V033-a.UCA | 8.472E-08 |
| V033-a.I1 | 9.326E-09 |
| V033-a.I2 | 5.259E-09 |
| V033-a.I3 | 2.710E-09 |
| V033-a.I4 | 1.662E-09 |
| V033-a.I5 | 1.872E-09 |
| V033-a.I6 | <1.0E-12 |
| V033-a.01 | <1.0E-12 |
| PGT145 | 1.012E-08 |

**Fig. S6. Affinity changes during V033-a lineage maturation.** (**A**) Sequences of V033-a lineage heavy (top) and light (bottom) chain intermediates. (**B**) Curves over time showing Biolayer Interferometry (BLI) binding kinetics of V033-a lineage Fabs for Q23 SCT27, a stabilized WT Q23 trimer. (**C**) Inferred $K_D$ values of V033-a lineage Fabs for Q23 SCT27.

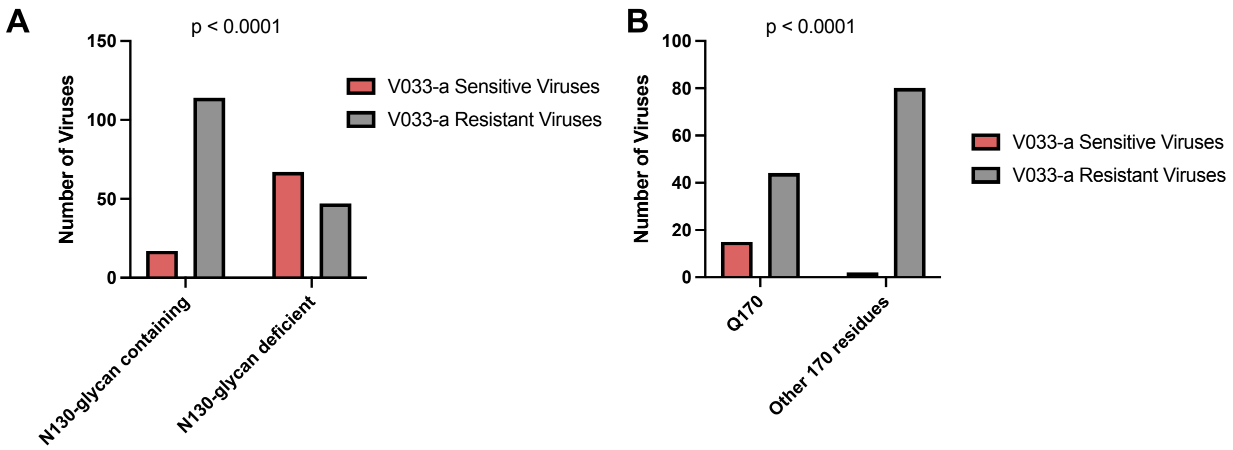


**Fig. S7. Env sensitivity and resistance signatures to the V033-a bNAb lineage.** (**A**) The number of viruses sensitive and resistant to neutralization by the V033-a.01 bNAb that contain or lack a glycan at N130 ($p < 0.0001$; Fischer's Exact test). (**B**) The number of N130-glycan containing viruses sensitive and resistant to neutralization by the V033-a lineage grouped by the presence or absence of Q170 ($p < 0.0001$; Fischer's Exact test). These numbers were calculated using data reported in Roark et al. (*28*).

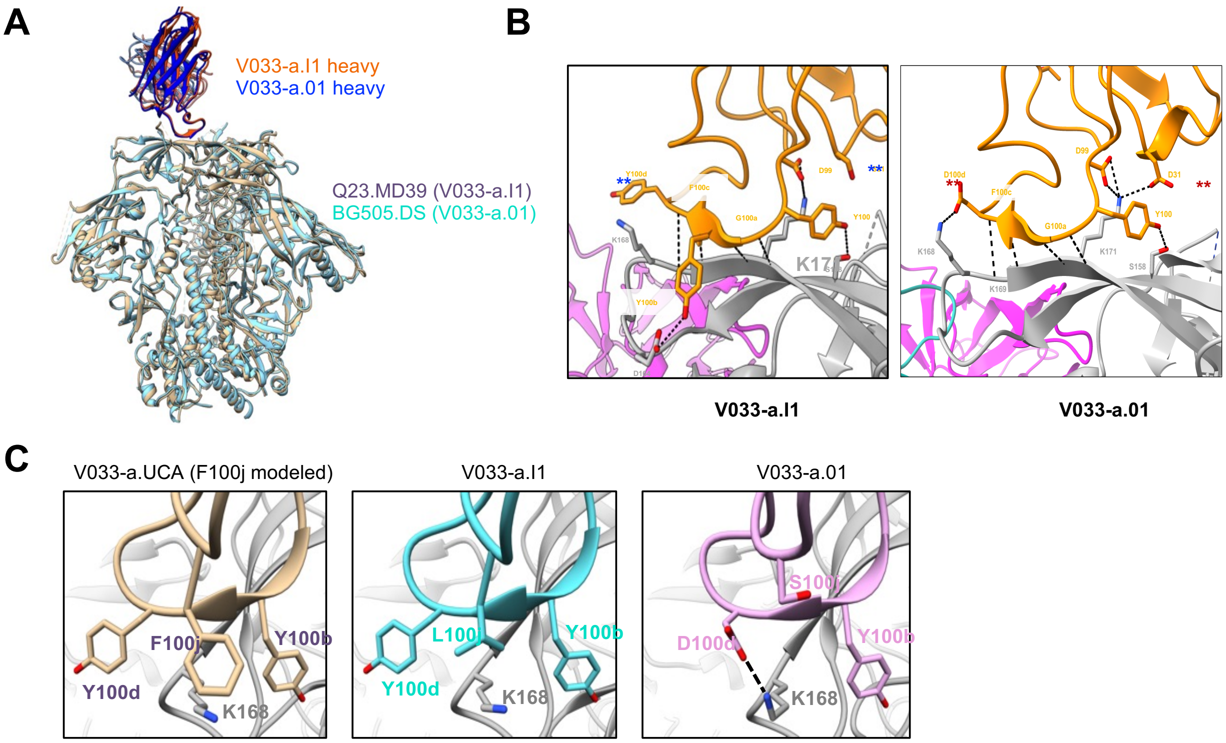


**Fig. S8. Structural maturation of the V033-a lineage (A)** Cryo-EM structure of the V033-a.I1 antibody in complex with a stabilized Q23.MD39 trimer aligned to the structure of the mature V033-a.01 bNAb (PDB: 9BNP). **(B)** Maturation of CDRH3 contacts with the V2 apex C-strand during V033-a lineage affinity maturation. **(C)** Maturation of CDRH3 interactions with Env K168 during V033-a lineage affinity maturation.

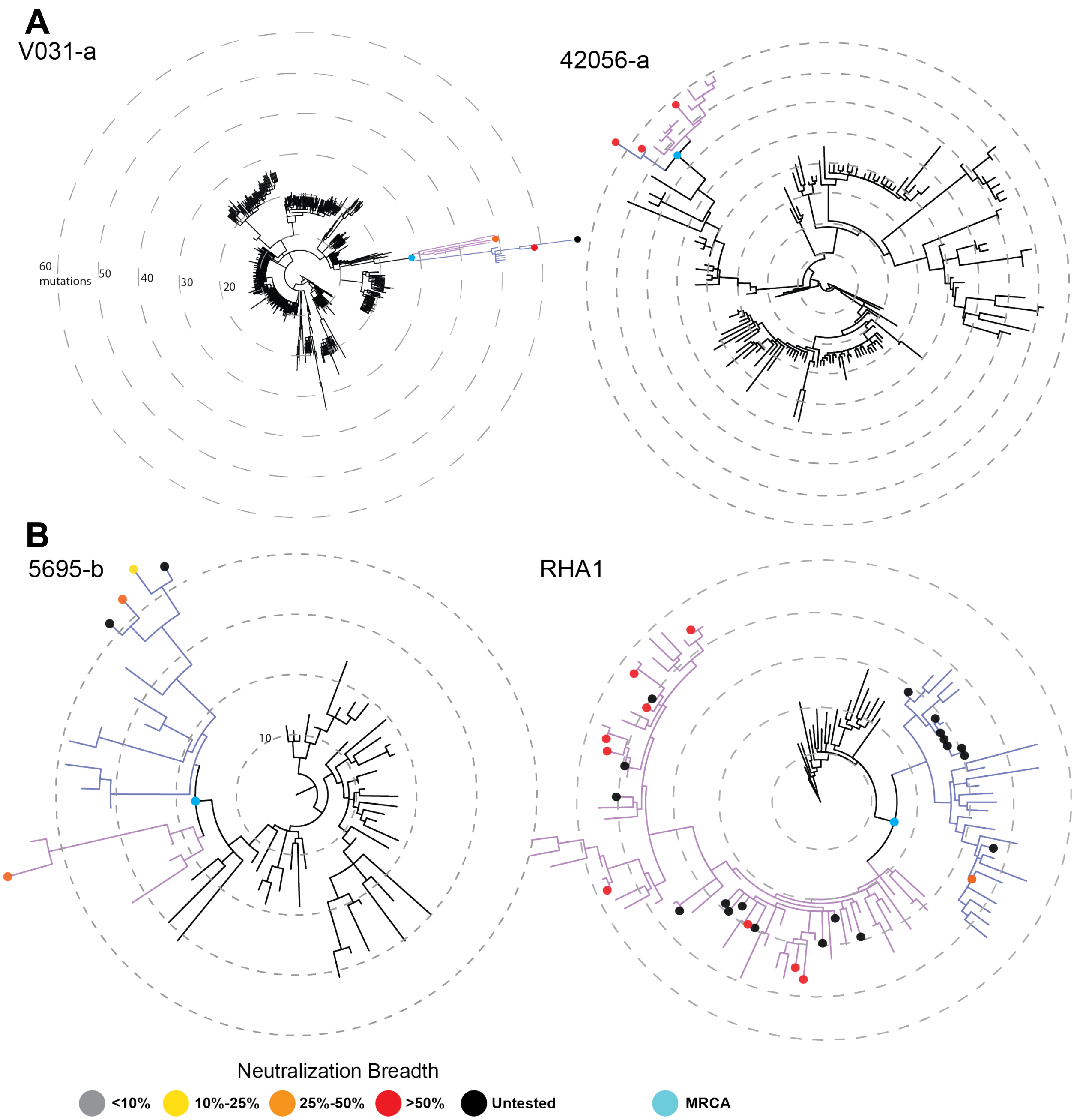


**Fig. S9. Heavy chain phylogenetic trees of V2 apex bNAb lineages.** Radial IgPhyML trees of V2 apex bNAb lineage heavy chains showing (**A**) late and (**B**) intermediate phylogenetic divergence. All trees are rooted on the lineage's inferred UCA. Gray, yellow, orange, red, and black dots indicate mature bNAb lineage members, colored by their neutralization breadth on our 18-virus panel reported in Roark et al. (*28*). Blue dots indicate the most recent common ancestor (MRCA) of all broadly neutralizing members of the lineage. Different clades branching off from the MRCA and independently acquiring breadth are colored indigo and lilac. A CDRH3 insertion in the V031-a lineage is indicated by an arrow. Scale bars are represented internally, with each concentric circle indicating 10 nucleotide mutations from the UCA.

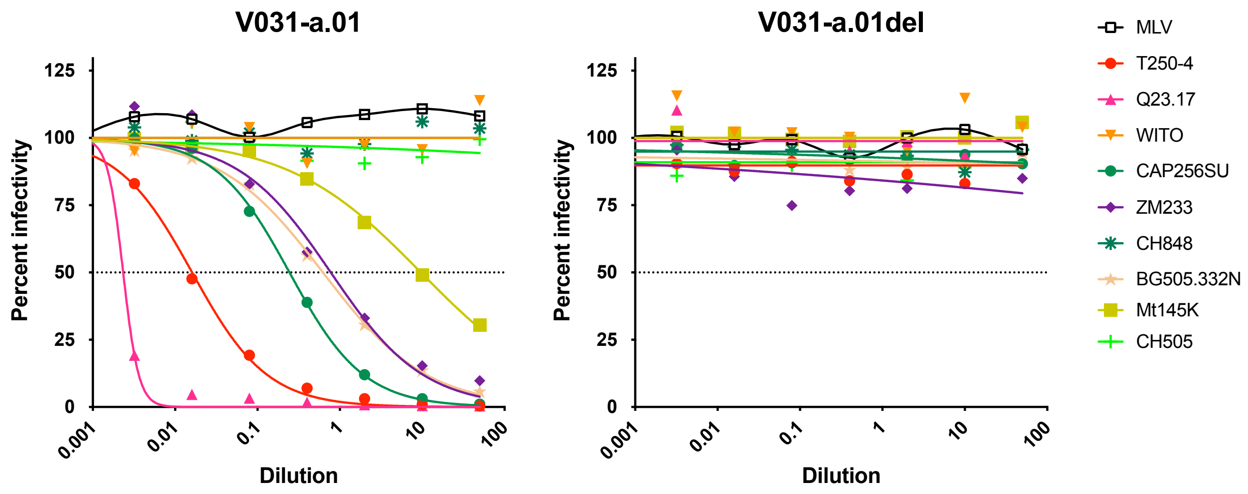


**Fig. S10. A two-residue insertion in necessary for heterologous neutralization by the V031-a.01 mAb.** Neutralization of a pseudovirus panel by the bNAb from RM V031 (V031-a.01) and a V031-a.01 mutant lacking a two residue CDRH3 insertion (V031-a.01del).

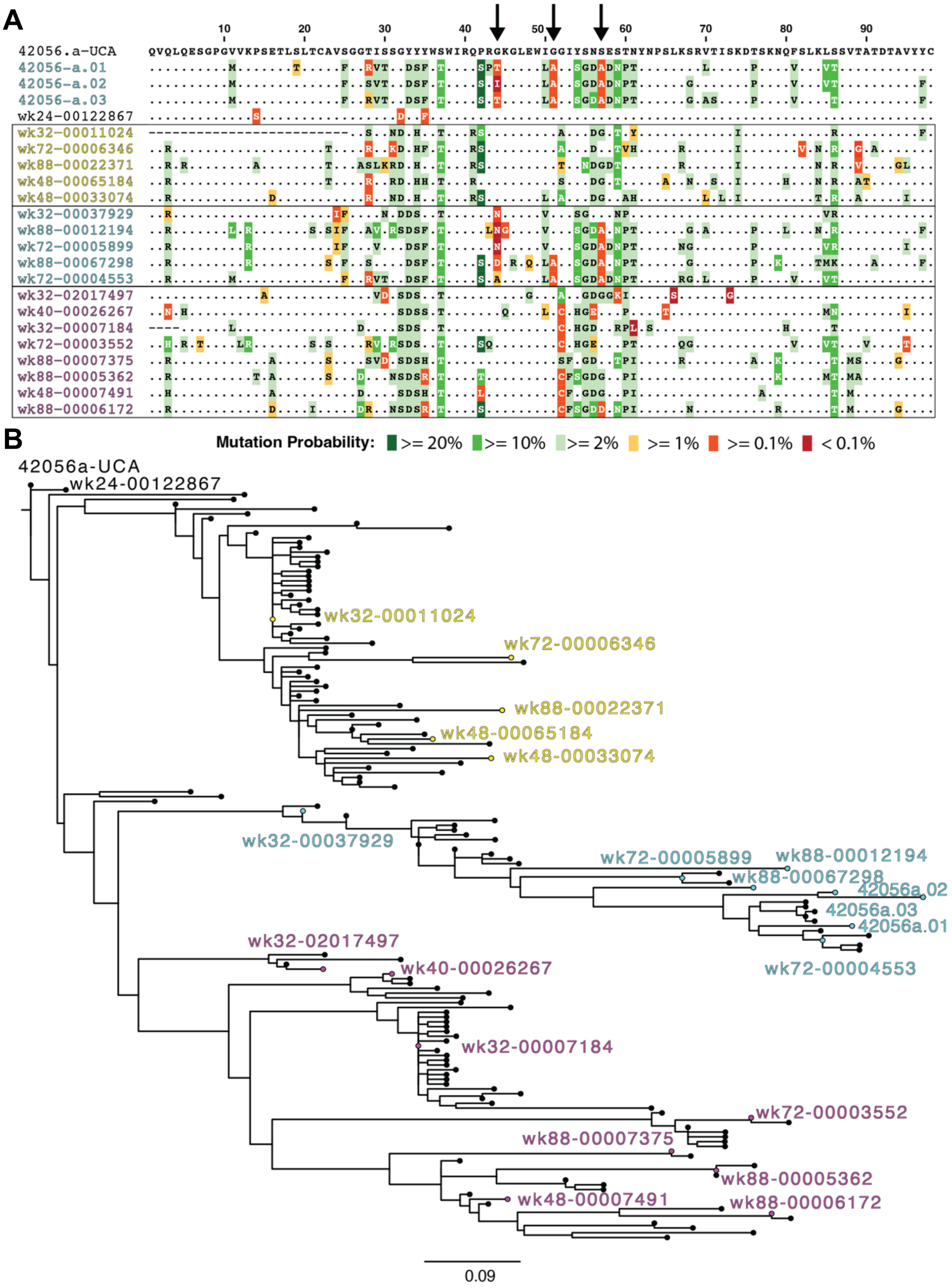
A
10
20
30
40
50
60
70
80
90
42056.a-UCA QVQLQESGPGVVKPSETLSLTCAVSGGTISSGYYYWSWIRQPRGKGLEWIGGIYSNSESTNYNPSLKSRVTISKDTSKNQFSLKLSSVTATDTAVYYC
42056-a.01
42056-a.02
42056-a.03
wk24-00122867
wk32-00011024
wk72-00006346
wk88-00022371
wk48-00065184
wk48-00033074
wk32-00037929
wk88-00012194
wk72-00005899
wk88-00067298
wk72-00004553
wk32-02017497
wk40-00026267
wk32-00007184
wk72-00003552
wk88-00007375
wk88-00005362
wk48-00007491
wk88-00006172
Mutation Probability: >= 20% >= 10% >= 2% >= 1% >= 0.1% < 0.1%
B
42056a-UCA
wk24-00122867
wk32-00011024
wk72-00006346
wk88-00022371
wk48-00065184
wk48-00033074
wk32-00037929
wk72-00005899
wk88-00012194
wk88-00067298
42056a.02
42056a.03
42056a.01
wk72-00004553
wk32-02017497
wk40-00026267
wk32-00007184
wk72-00003552
wk88-00007375
wk88-00005362
wk48-00007491
wk88-00006172
0.09

**Fig S11. Mutational bottleneck in the 42056-a lineage.** (**A**) The mutation probabilities of the amino acid mutations computed with ARMADiLLO for the 42056-a lineage mAbs and 19 representative NGS sequences. Only the V segment region is shown for clarity. Three highly improbable mutations (black arrows) that are shared only by the isolated bNAb mAbs from the 42056-a lineage (42056-a.01, 42056-a.02, 42056-a.03) and some members of their clade (colored blue). (**B**) An IgPhyML tree of the 42056-a lineage heavy chain showing the positions of sequences from **A**.

**A**

| Eliciting SHIV | | T250-4 | Q23.17 | WITO.4160 | ZM233.6 | CAP256SU | CH505 T/F | CH848 T/F | CH848wk36con | CE1176 |
|---|---|---|---|---|---|---|---|---|---|---|
| CAP256SU | 40591-a.UCA | >200 | >200 | >200 | >200 | >200 | >200 | >200 | >200 | >200 |
| | 42056-a.UCA | >200 | >200 | >200 | >200 | >200 | >200 | >200 | >200 | >200 |
| | 42056-b.UCA | >200 | >200 | >200 | >200 | >200 | >200 | >200 | >200 | >200 |
| Ce1176 | 6561-a.UCA | >200 | >200 | >200 | >200 | >200 | >200 | >200 | >200 | >200 |
| Q23.17 | 41328-a.UCA | >200 | >200 | >200 | >200 | >200 | >200 | >200 | >200 | >200 |
| | V031-a.UCA | >200 | 183 | >200 | >200 | >200 | >200 | >200 | >200 | >200 |
| | V033-a.UCA | >200 | 106 | >200 | >200 | >200 | >200 | >200 | >200 | >200 |
| CH848wk36con | 44715-a.UCA | >200 | >200 | >200 | >200 | >200 | >200 | >200 | >200 | >200 |
| CH505 | RHA1.UCA | >200 | >200 | >200 | >200 | >200 | >200 | >200 | >200 | >200 |
| | 5695-b.UCA | >200 | >200 | >200 | >200 | >200 | >200 | >200 | >200 | >200 |
| | 6070-a.UCA | >200 | >200 | >200 | >200 | >200 | >200 | >200 | >200 | >200 |
| | T646-a.UCA | >200 | >200 | >200 | >200 | >200 | >200 | >200 | >200 | >200 |

**B**

| | | CH505 | Q23 | CAP256 SU | T250 | WITO | 246F3 |
|---|---|---|---|---|---|---|---|
| 6070 UCA | WT | >200 | >200 | >200 | >200 | >200 | >200 |
| | N160K | 166 | 0.087 | >200 | 7.8 | 45 | 0.059 |
| T646 UCA | WT | >200 | >200 | >200 | >200 | >200 | >200 |
| | N160K | >200 | 147 | >200 | >200 | 47 | >200 |

**Fig. S12. Neutralization phenotype of inferred V2 apex bNAb UCAs.** (**A**) Neutralization $IC_{50}$ values of all rhesus V2 apex bNAb UCAs against a panel of Envs previously described to be sensitive to neutralization by V2 apex bNAb precursors. (**B**) Neutralization $IC_{50}$ values of the 6070-a.UCA and T646-a.UCA against a panel of autologous and heterologous N160 glycan-deficient viruses.

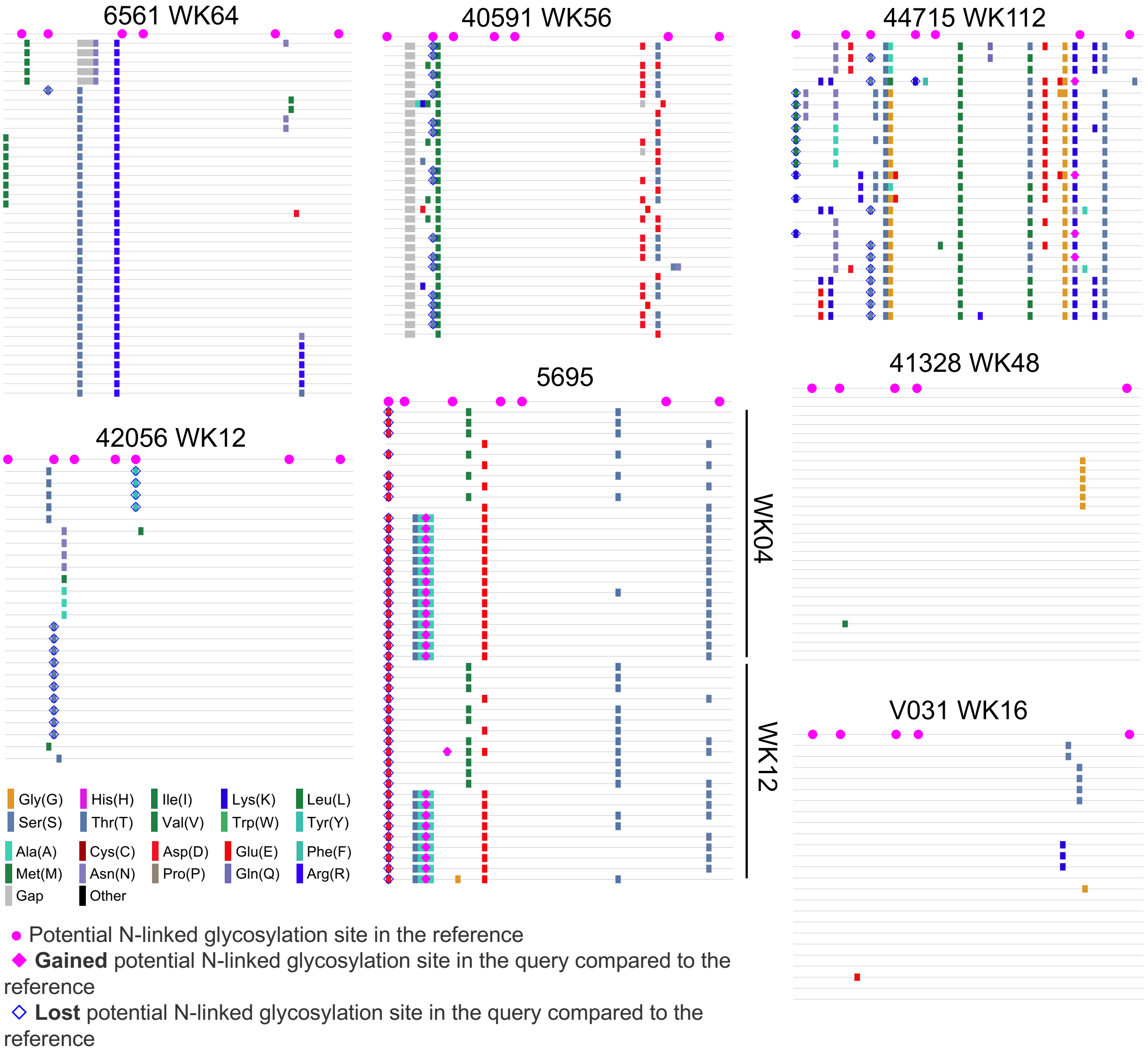


**Fig. S13. T/F V1V2 sequences are usually absent preceding bNAb initiation.** Highlighter plots showing alignments of Env SGS V1V2 sequences (residues 130-199) at the timepoint prior to when V2 apex bNAb lineages first became detectable by B cell NGS. In each plot, the top sequence represents the T/F Env sequence, and mismatches to the T/F Env sequence are highlighted.

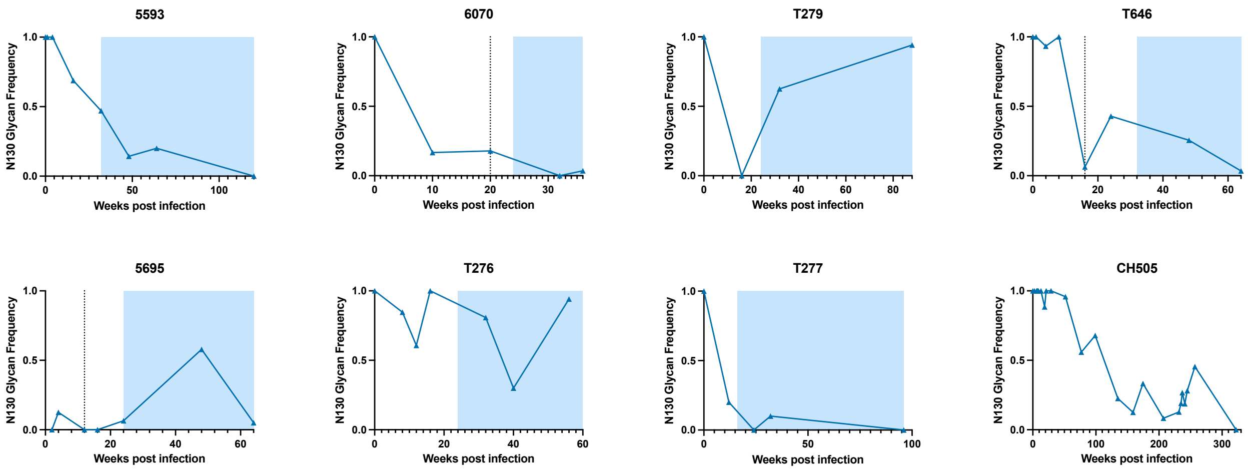


**Fig. S14. N130 glycan deletion precedes V2 apex bNAb elicitation in SHIV-CH505-infected RMs.** Longitudinal glycan sequon frequencies at residue 130 in SHIV-CH505-infected RMs that developed V2 apex bNAbs. Dotted lines indicate the timepoint where the V2 apex bNAb lineage was first detected by B cell NGS in RMs where NGS data is available. Shaded regions indicate timepoints where heterologous tier-2 plasma neutralization are observed. Longitudinal glycan sequon frequencies at residue 130 in human participant CH505 are included in the bottom right panel for comparison.

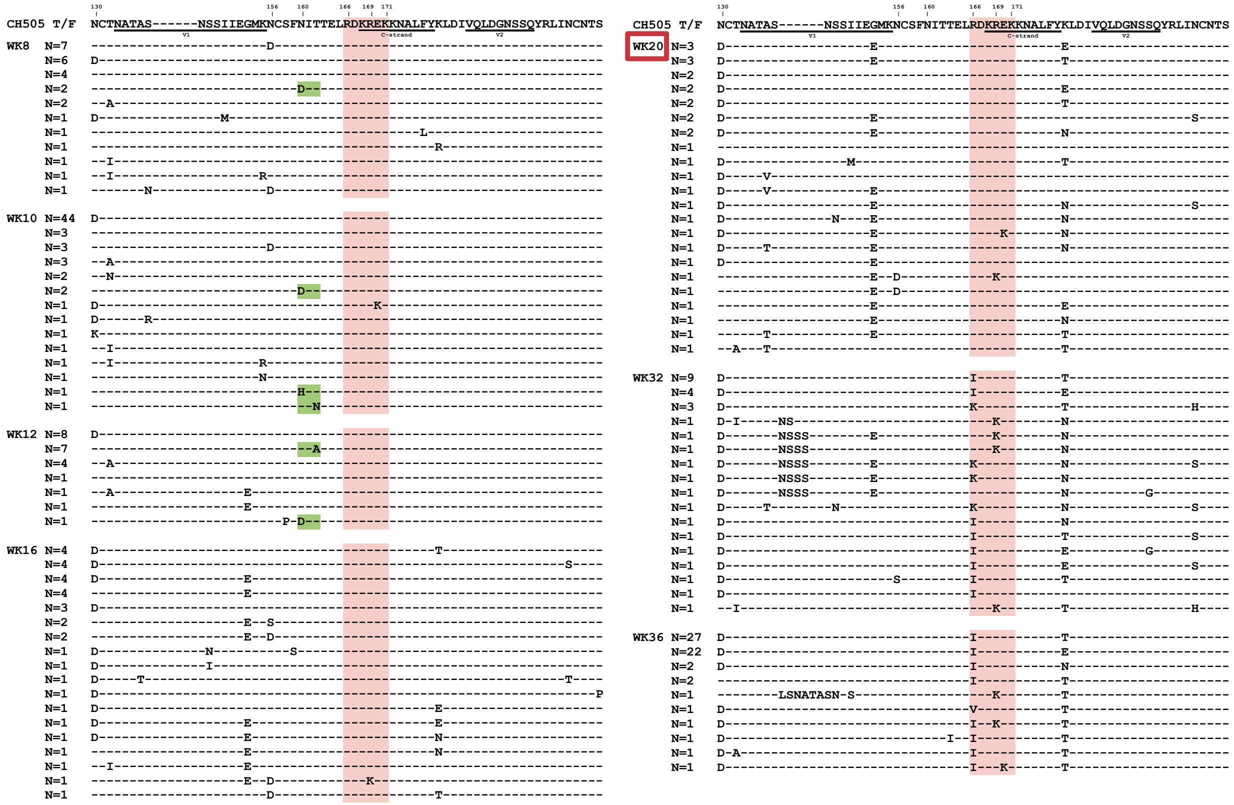


**Fig. S15. V1V2 sequence evolution in RM 6070.** Amino acid alignments of longitudinal Env sequences obtained by SGS of circulating plasma virion RNA from RM 6070. Sequences show residues 130-199 (HXB2 numbering). Sequences are grouped by timepoint indicated on the left in weeks. Dashes indicate identity to the T/F sequence. C-strand residues 166-171 are shaded red. Mutations that delete the glycan at N160 are shaded green. A red box indicates the timepoint when the bNAb lineage was first detected.

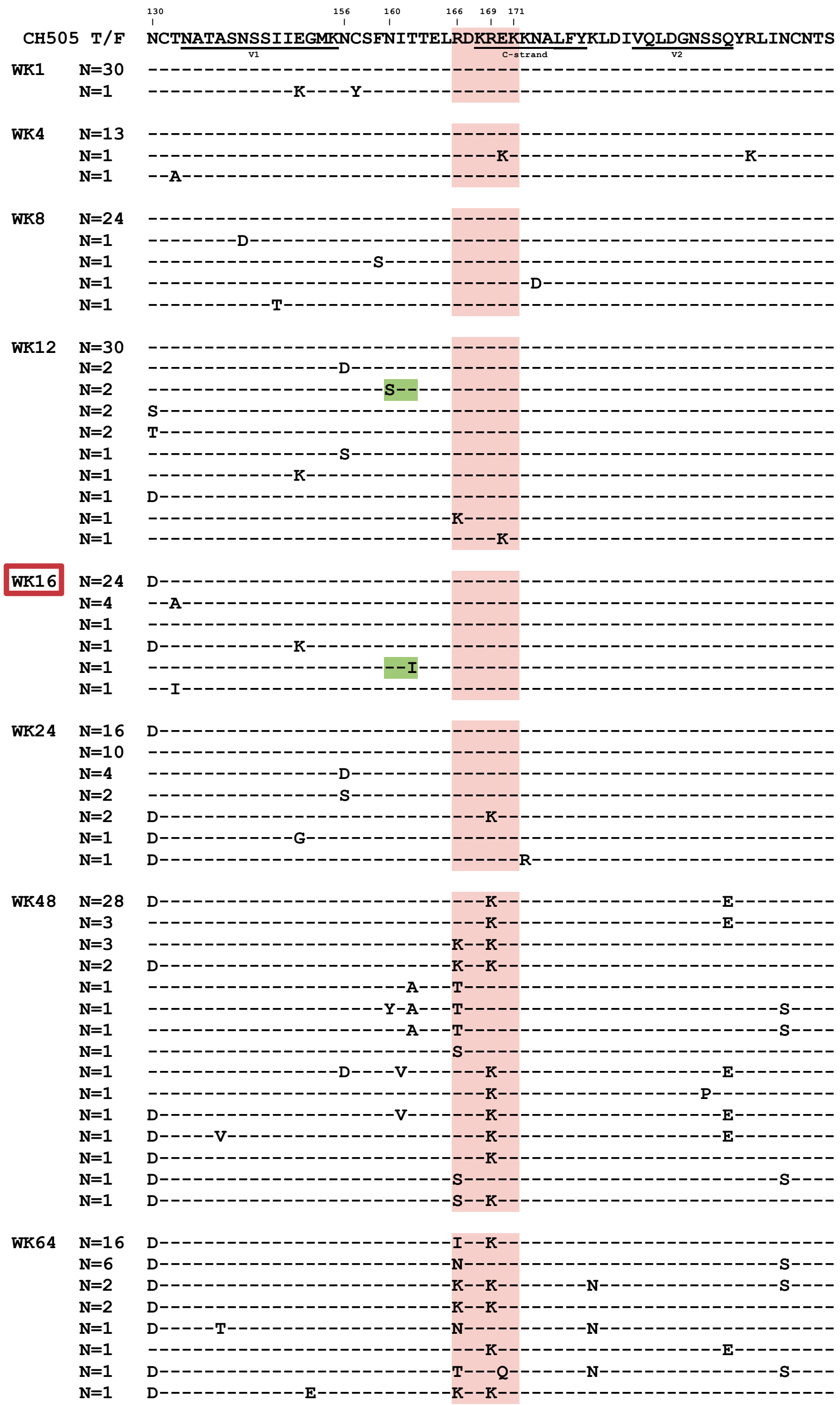


**Fig. S16. V1V2 sequence evolution in RM T646.** Amino acid alignments of longitudinal Env sequences obtained by SGS of circulating plasma virion RNA from RM T646. Sequences show residues 130-199 (HXB2 numbering). Sequences are grouped by timepoint indicated on the left in weeks. Dashes indicate identity to the T/F sequence. C-strand residues 166-171 are shaded red. Mutations that delete the glycan at N160 are shaded green. A red box indicates the timepoint when the bNAb lineage was first detected.

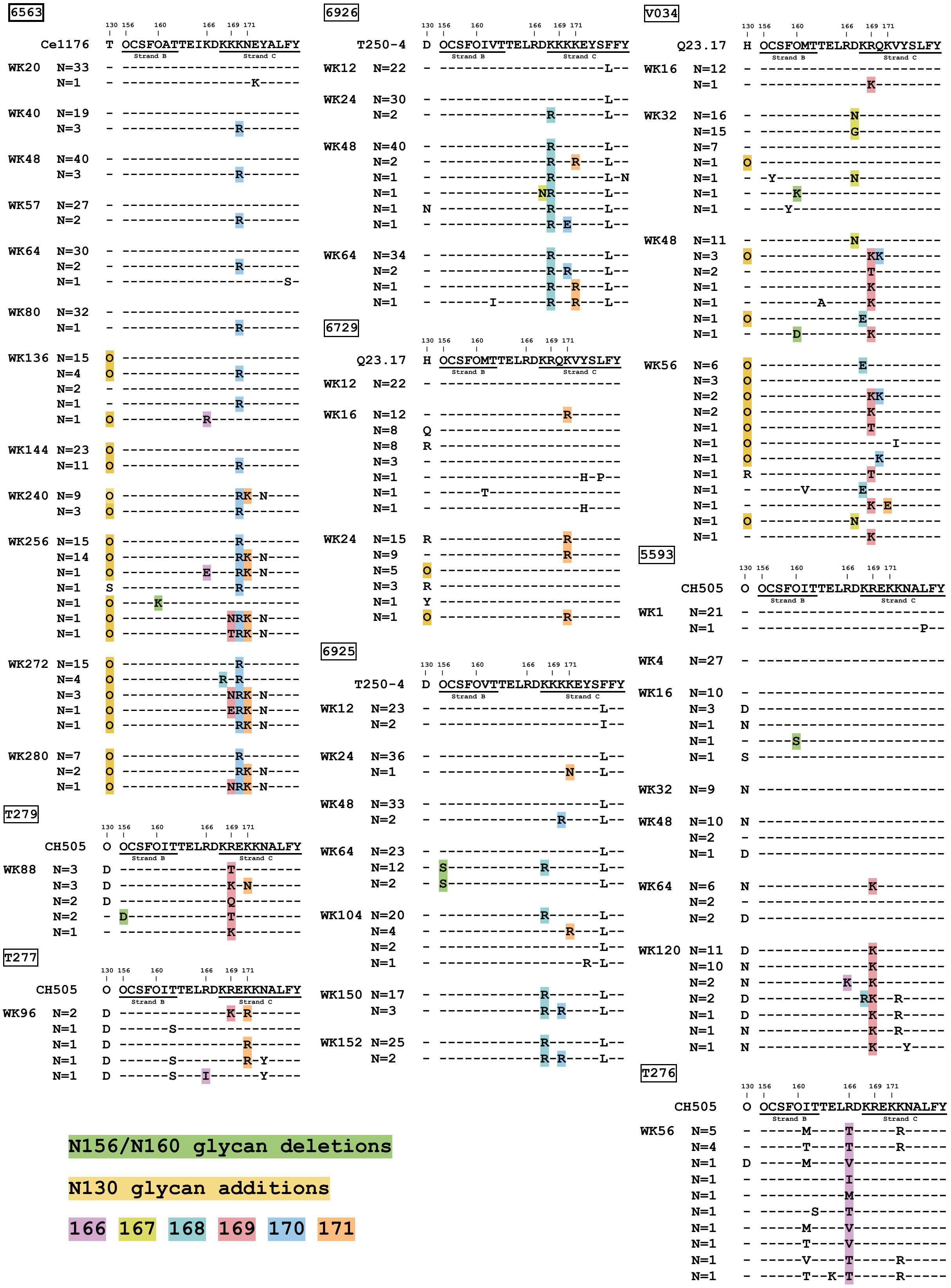
6563
Ce1176 T OCSFOATTEIKDKKKNEYALFY
6926
T250-4 D OCSFOIVTTELRDKKKKEYSFFY
6729
Q23.17 H OCSFOMTTELRDKRQKVYSLFY
6925
T250-4 D OCSFOVTTELRDKKKKEYSFFY
T279
CH505 O OCSFOITTELRDKREKKNALFY
T277
CH505 O OCSFOITTELRDKREKKNALFY
V034
Q23.17 H OCSFOMTTELRDKRQKVYSLFY
5593
CH505 O OCSFOITTELRDKREKKNALFY
T276
CH505 O OCSFOITTELRDKREKKNALFY
Strand B
Strand C
N156/N160 glycan deletions
N130 glycan additions
166 167 168 169 170 171

**Fig. S17. Limited and convergent Env mutations guide rhesus V2 apex bNAb maturation.** Amino acid alignments of longitudinal Env sequences obtained by SGS of circulating plasma virion RNA in RMs with V2 apex bNAbs. Sequences show residues 130 and 156-177 (HXB2 numbering). Sequences are grouped by timepoint indicated on the left in weeks. Dashes indicate identity to the T/F sequence. Mutations at residues 166-171, mutations that delete glycans at N156 or N160, or mutations that introduce a glycan to residue 130 are highlighted. Glycan sequons are denoted by "O."

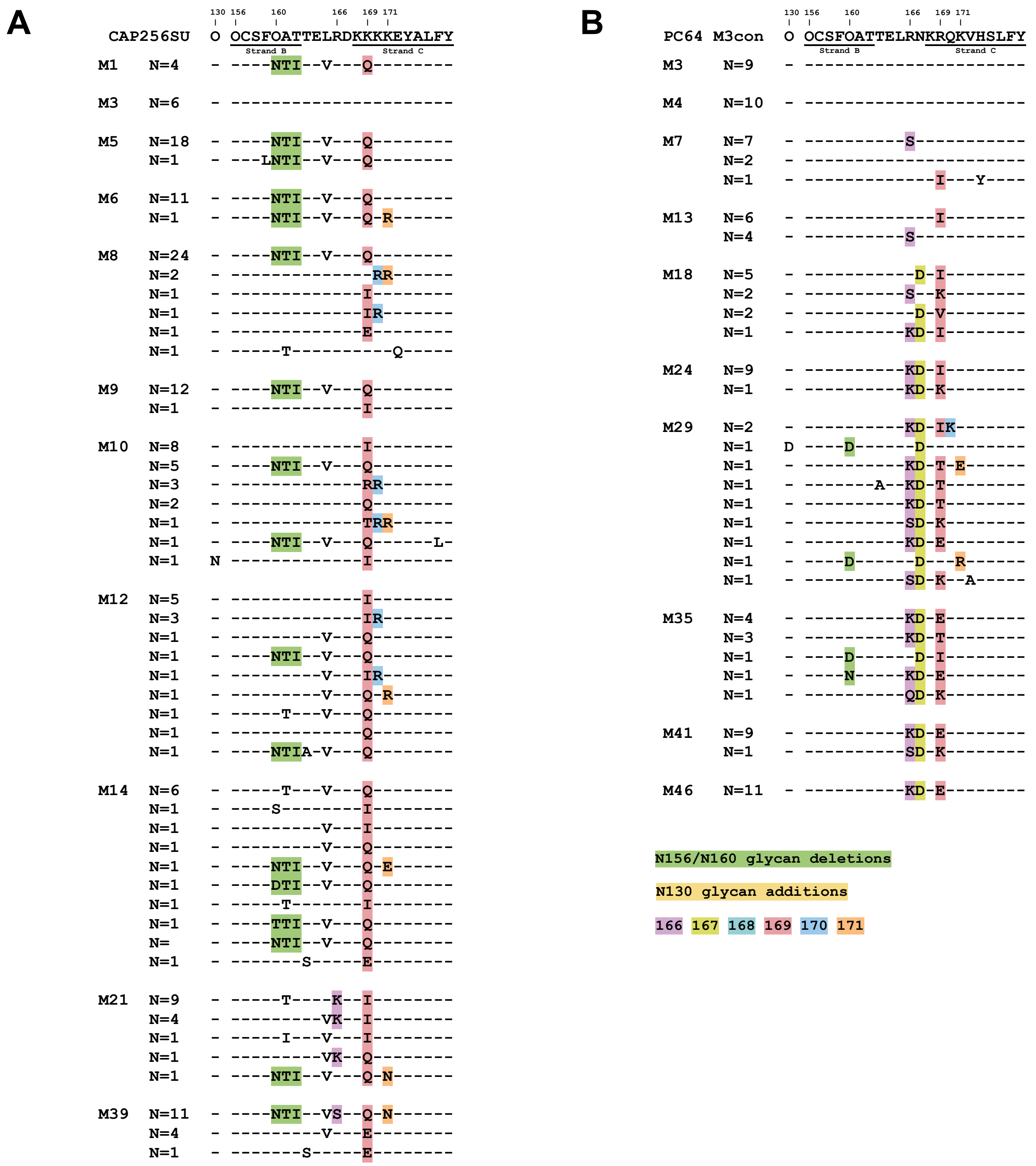

**Fig. S18. Limited and convergent Env mutations guide human V2 apex bNAb maturation.** Longitudinal Env SGS in human participants (**A**) CAP256 and (**B**) and PC64 who made V2 apex bNAbs. Mismatches in (**A**) are shown against the superinfecting strain CAP256SU. Dashes indicate sequence identity. Mutations at residues 166-171, mutations that delete glycans at N156 or N160, or mutations that insert glycans at residue 130 are highlighted. Sequences are grouped by timepoint indicated on the left in months. Data in this figure are re-analyzed from Doria-Rose et al. Nature 2014 and Landais et al. Immunity 2017 (*23, 46*).

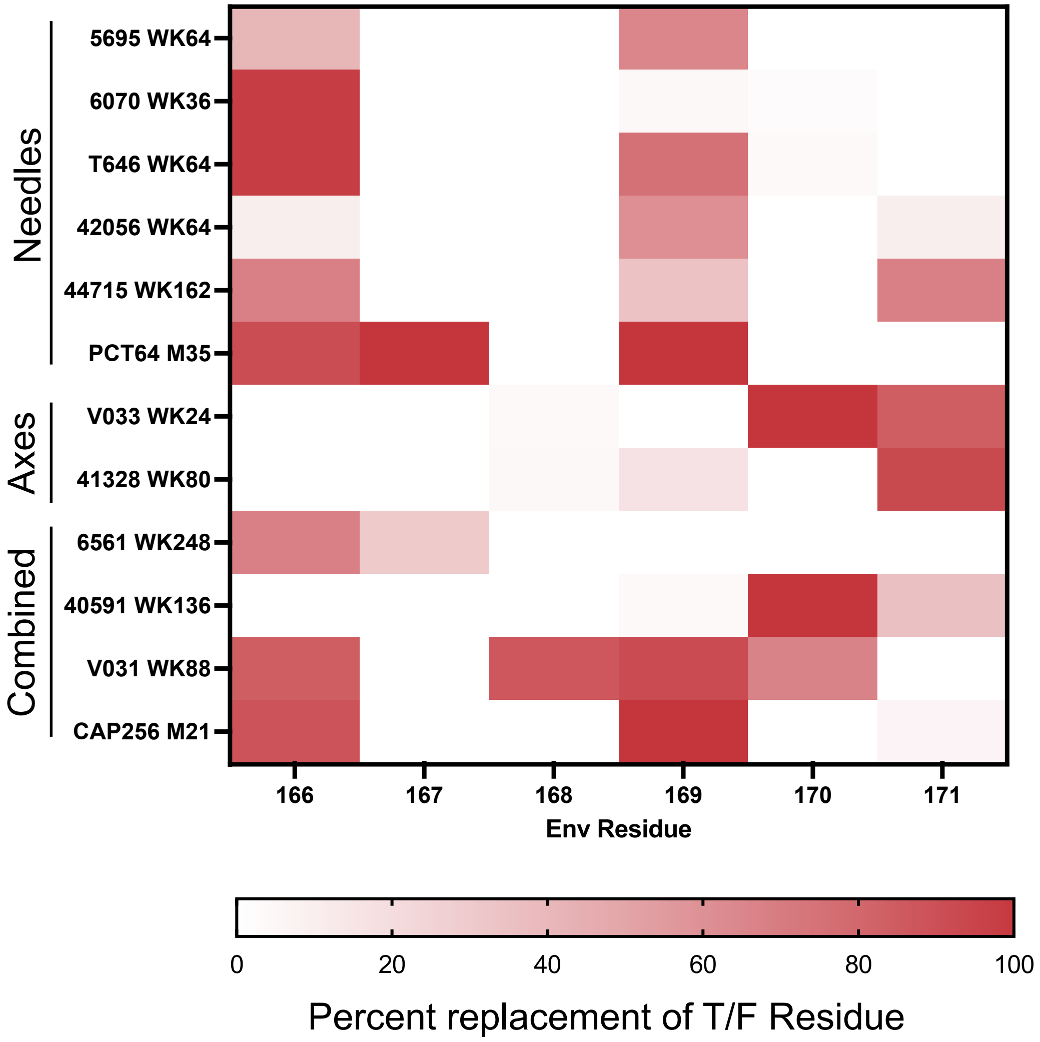


**Fig. S19. Env selection correlates with bNAb structural footprints.** Heatmap of the percent replacement of Env C-strand residues 166-171 in SGS sequences from humans and RMs with V2 apex bNAbs at the timepoint indicated. The structural class of each V2 apex bNAb is indicated on the left.

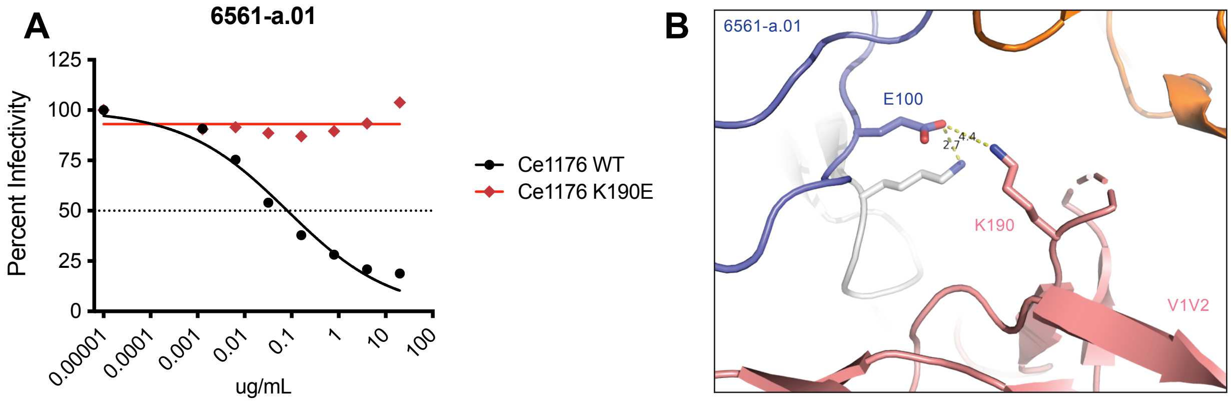


**Fig. S20. A K190E V2 hypervariable loop mutation mediates initial escape from the 6561-a bNAb lineage.** (**A**) Neutralization of Ce1176 wild-type and K190E mutant viruses by the 6561-a.01 mAb. (**B**) Inset from cryo-EM structure of 6561-a.01 in complex with an autologous Ce1176-SOSIP (PDB: 9BTJ) to highlight electrostatic contact between HCDR3 residue E100 and Env hypervariable V2 residue K190.

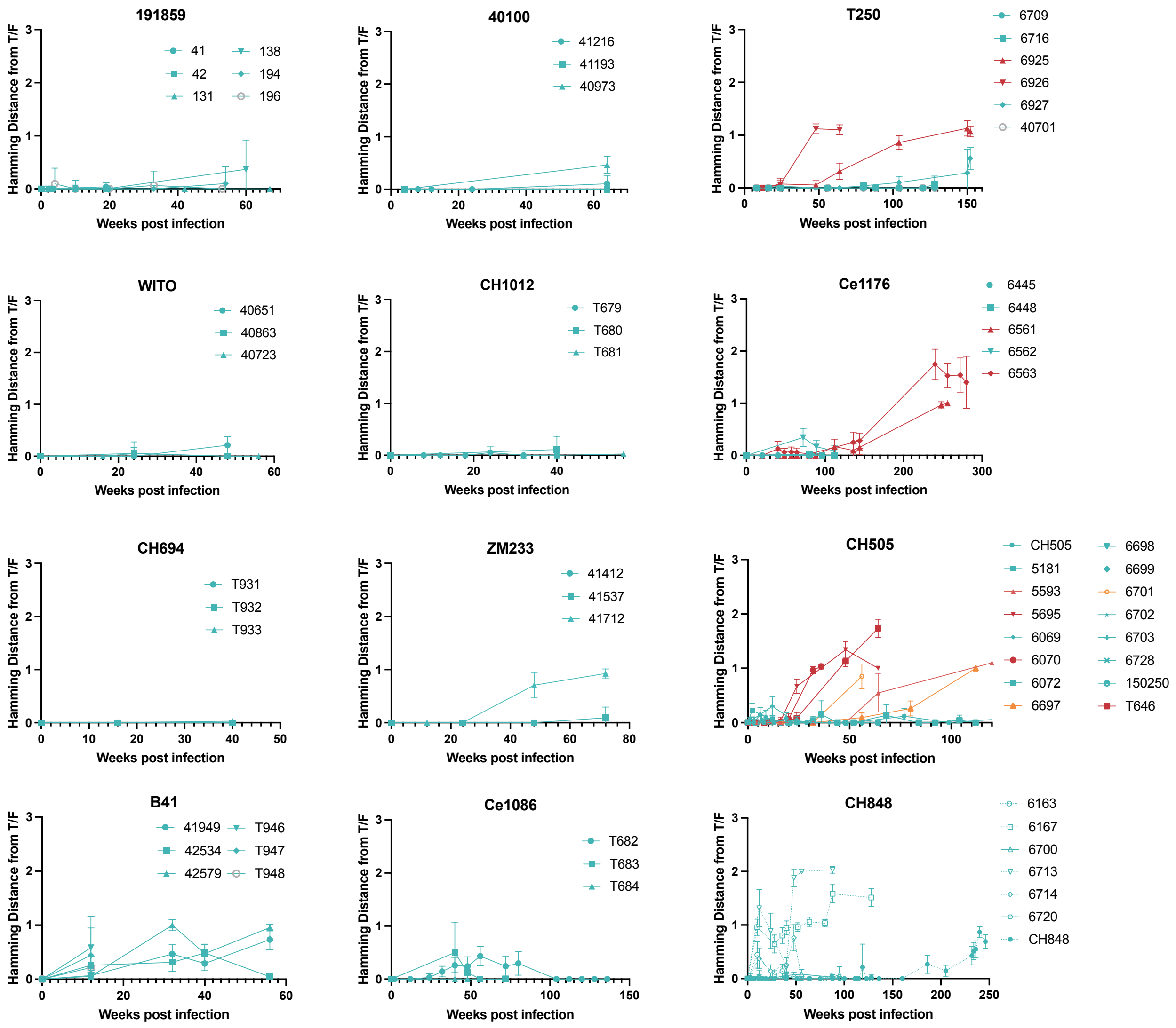


**Fig. S21. Longitudinal C-strand Hamming distances in SHIV-infected RMs.** Mean longitudinal Hamming distances of Env residues 166-171 from the T/F sequence in 101 SHIV-infected rhesus macaques. Data from human trial participants from which the SHIV strains were isolated are included where available. For all panels, RMs with V2 apex bNAbs are colored red and RMs without are colored teal. Data for the remaining SHIVs are shown in Figure 6B. RMs with C-strand targeted antibody responses that did not meet our criteria for breadth are colored orange. All error bars indicate 95% confidence intervals.

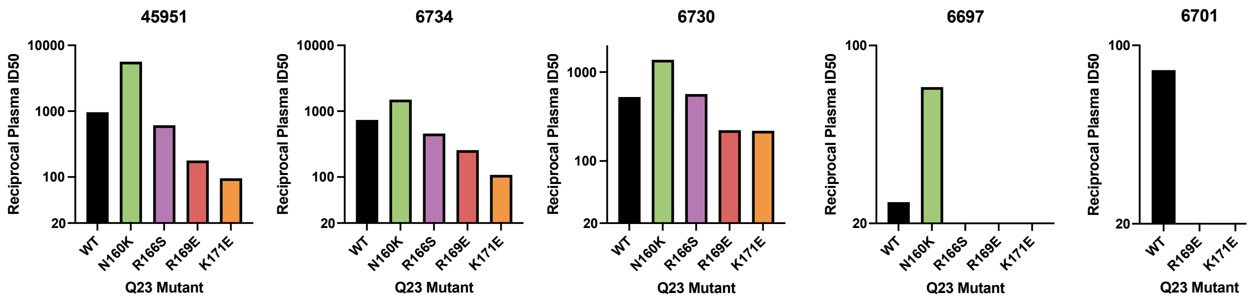


**Fig. S22. Evidence of C-strand targeted NAbs in five RMs.** Neutralization mapping against Q23.17 wild-type and V2 apex-epitope mutant pseudoviruses by plasma from SHIV-Q23.17 infected RMs 45951, 6734, 6730, and SHIV-CH505 infected RMs 6697 and 6701.

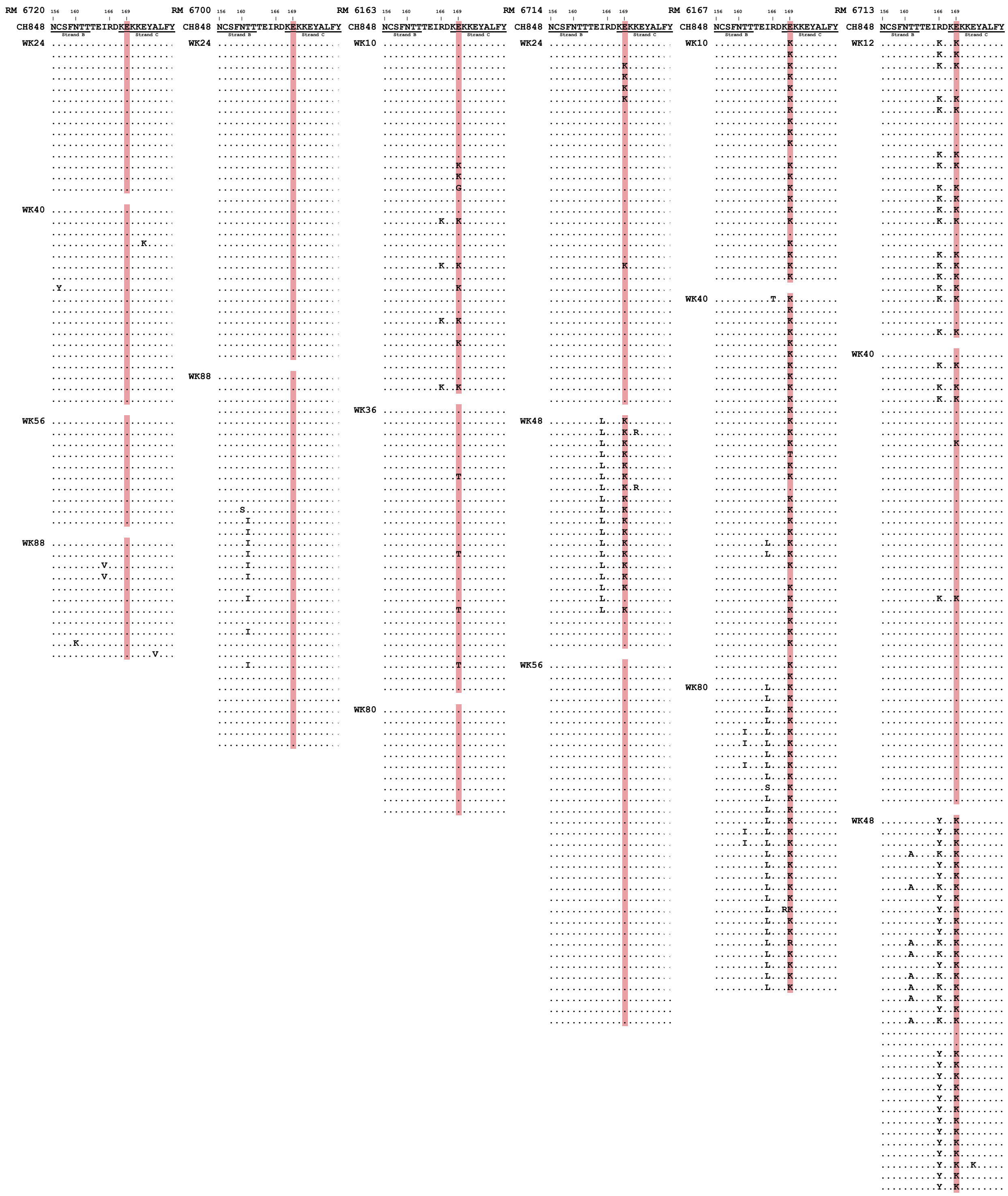


**Fig. S23. C-strand selection in SHIV CH848-infected RMs usually consists of an E169K reversion to group M consensus.** Longitudinal Env SGS alignments of V2 apex strands B and C from SHIV-CH848-infected RMs. Dots indicate identical residues. Residue 169 is highlighted red.

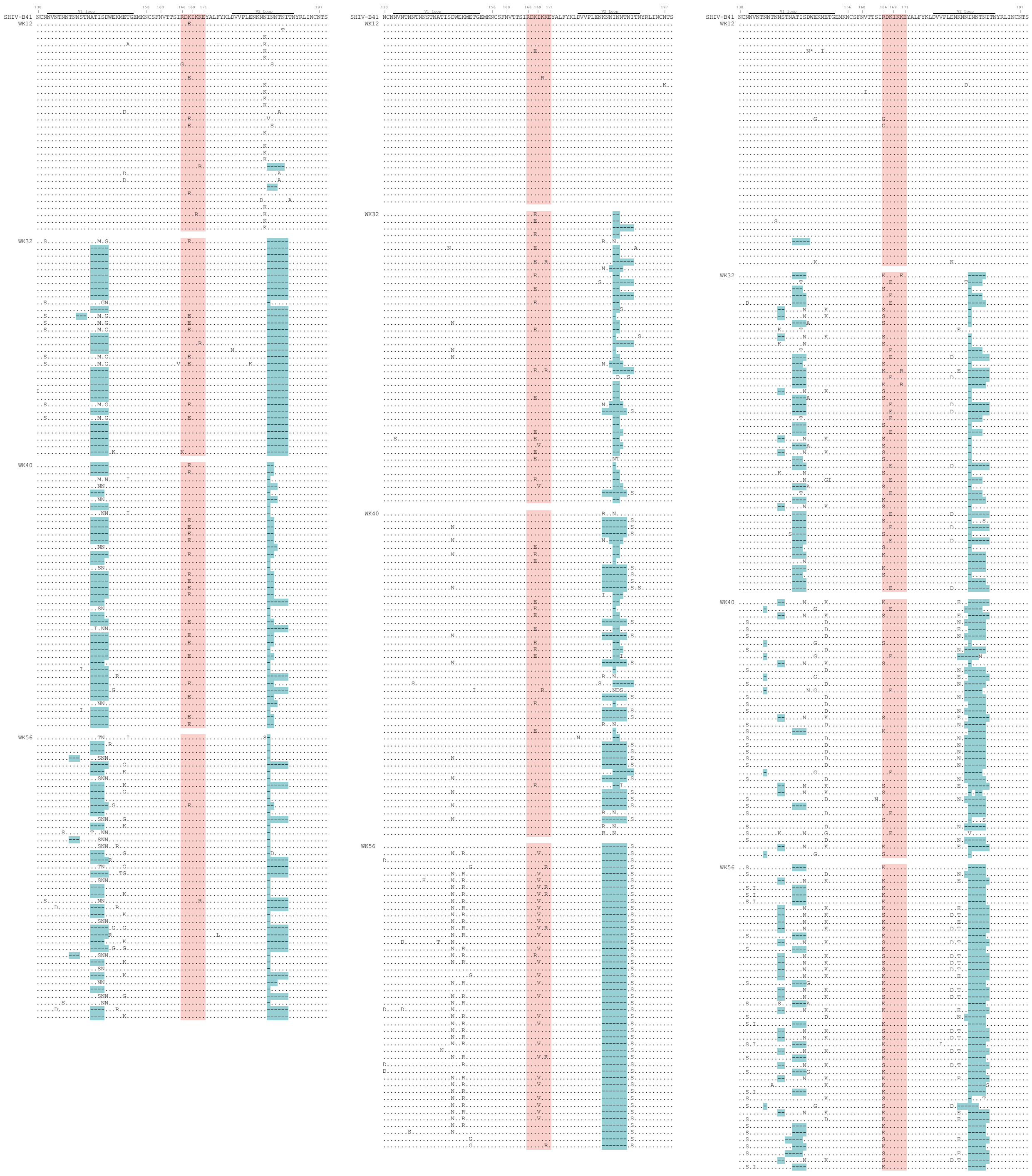

**Fig. S24. C-strand selection in SHIV B41-infected RMs.** Longitudinal Env SGS alignments of the V1V2 region from SHIV-B41-infected RMs 41949 (left), 42534 (middle), and 42579 (right). Dots indicate identical residues, while dashes represent gaps in the alignment. C-strand residues 166-171 are highlighted red, while V1/V2 hypervariable loop deletions are highlighted teal.

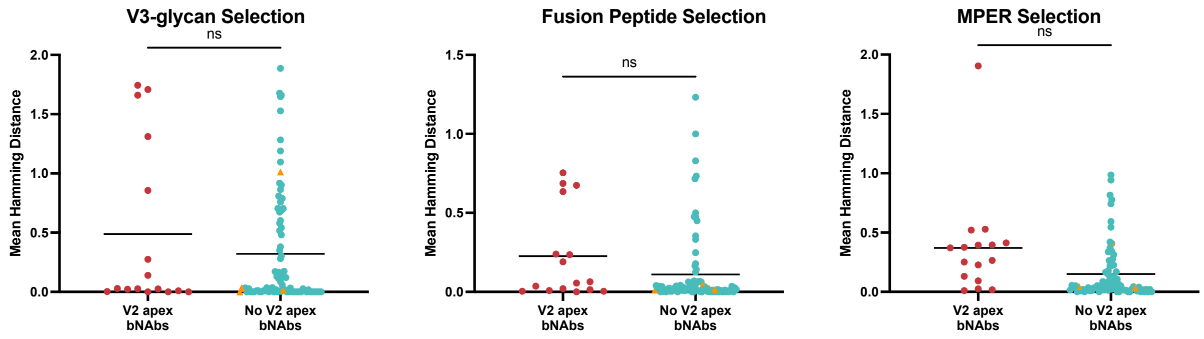


**Fig. S25. Selection at other bNAb epitopes is not significantly associated with the development of V2 apex bNAbs.** Comparison of the mean C-strand Hamming distances between RMs with and without V2 apex bNAbs at the bNAb epitopes indicated. Epitopes analyzed include the V3-glycan (residues 324-334), fusion peptide (residues 512-525) and MPER (residues 660-678). Significance was determined by Welch's t-test. ns = not significant.

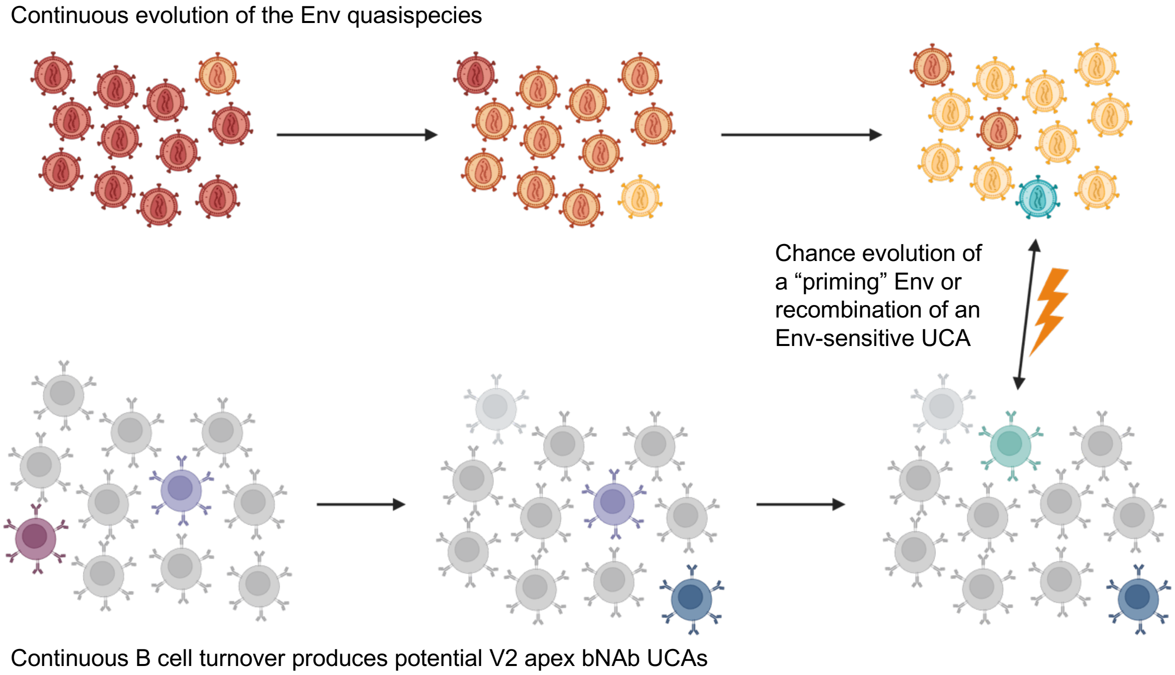


**Fig. S26. Model of V2 apex bNAb priming.**

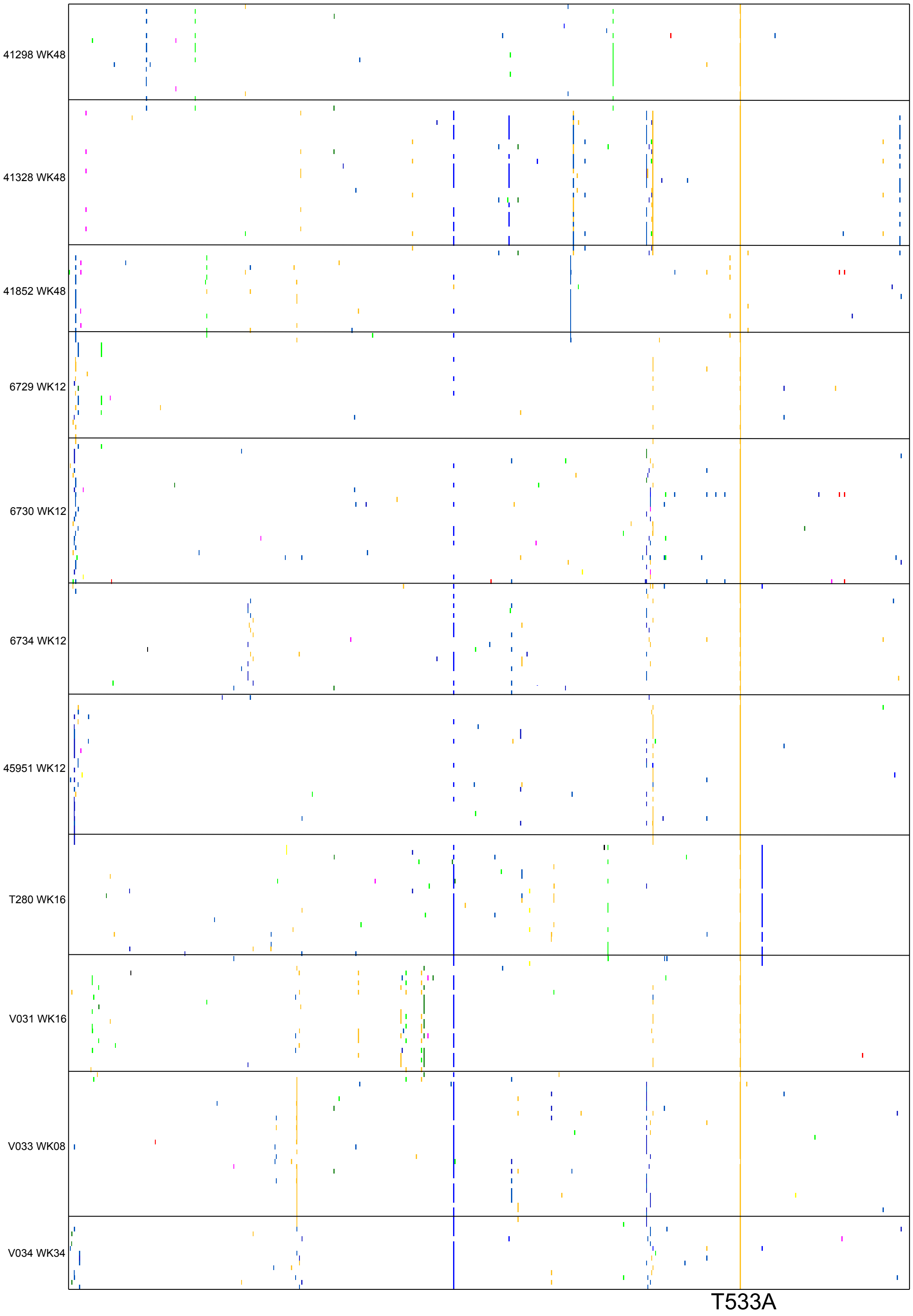


**Fig. S27. An early fitness reversion in SHIV-Q23.17 infected RMs.** A pixel plot of Env sequences from the first sequenced timepoint from each SHIV-Q23.17 infected RM. Mismatches to the T/F sequence are colored. A universally detected T533A reversion to HIV-1 group M consensus is indicated below.

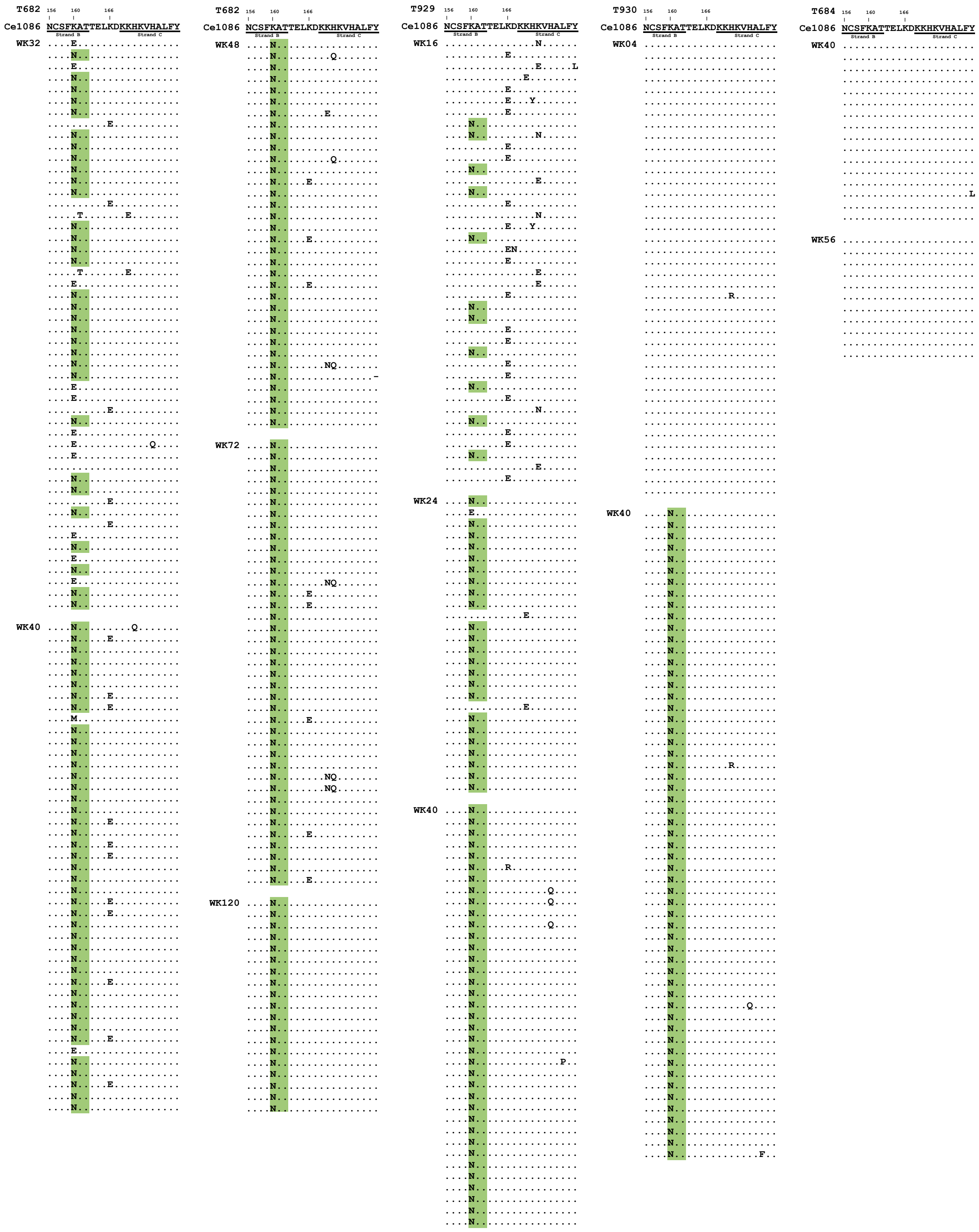


**Fig. S28. Transient C-strand selection in SHIV-Ce1086 infected RMs reverts after replacement of the N160 glycan.** Longitudinal Env SGS alignments of V2 apex strands B and C from SHIV-CH1086-infected RMs that exhibited C-strand selection. Dots indicate identical residues. N160 glycan additions are highlighted green.

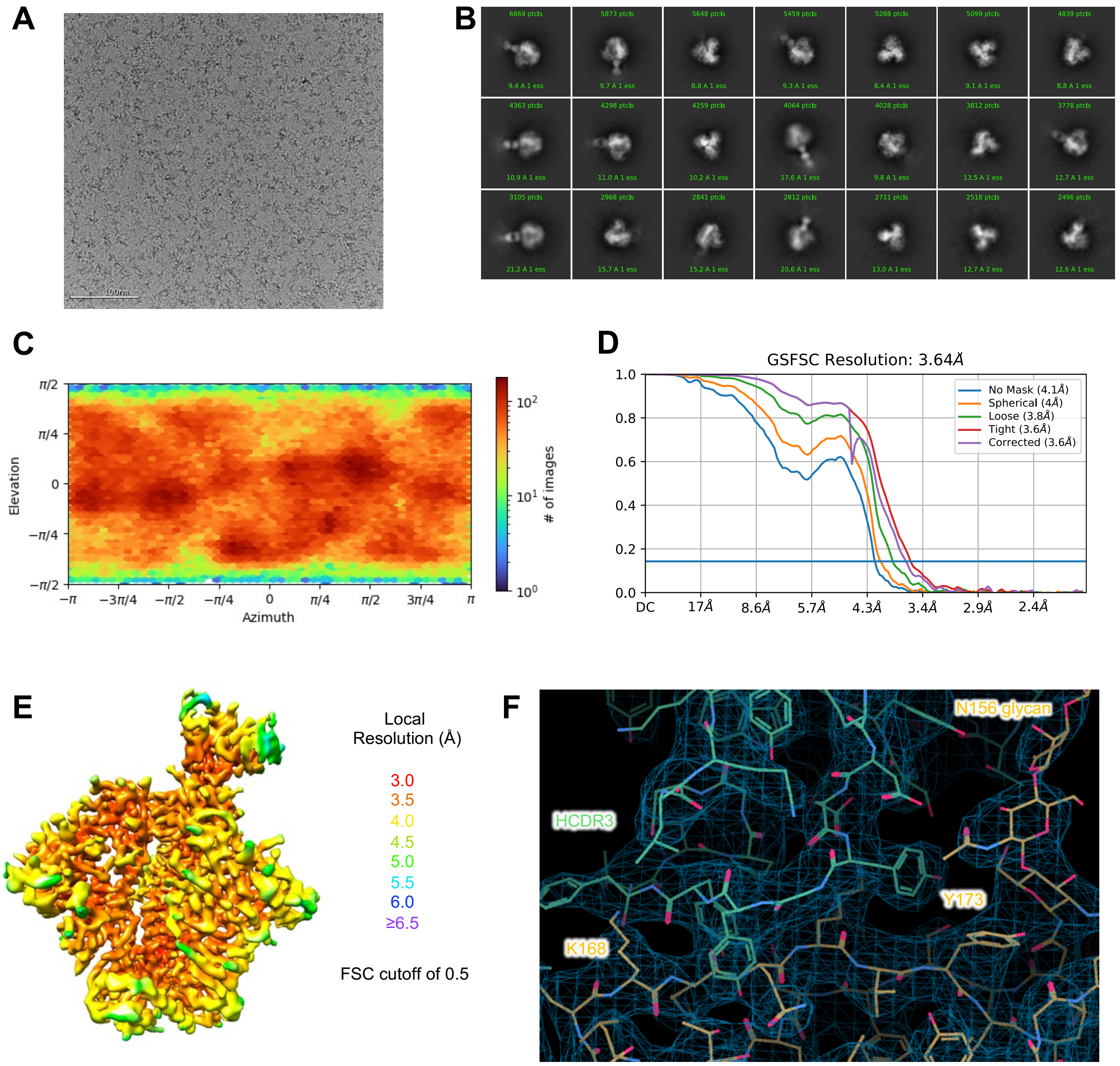


**Fig. S29. Cryo-EM details of V033-a.I1 in complex with Q23.17 MD39 envelope trimer.** (**A**) Representative raw micrograph is shown. (**B**) Representative 2D class averages of picked particles are shown. (**C**) The orientations of all particles used in the final refinement are shown as a heatmap. (**D**) The gold-standard fourier shell correlation (FSC) at threshold of 0.143 resulted in a resolution of 3.64 Å using non-uniform refinement with C1 symmetry. (**E**) The local resolution of the full map is shown as generated through cryoSPARC using an FSC cutoff of 0.5. (**F**) Cryo-EM 3D reconstruction density to highlight the Fab-trimer interactive surface.

| | V033-a.I1 in complex with Q23.17 MD39 Env |
|---|---|
| **PDB** | **9OMG** |
| **EMDB** | **70613** |
| **Data collection & processing** | |
| Microscope | FEI Titan Krios |
| Camera | Gatan K3 |
| Magnification | 105,000x |
| Voltage (kV) | 300 |
| Electron dose ($e^-/Å^2$) | 58 |
| Defocus range (µm) | 0.8 - 2.0 |
| Pixel size (Å) | 0.83 |
| Micrographs collected | 6,268 |
| Software | cryoSPARC v4.1 |
| Micrographs used | 5,382 |
| Final refined particles | 155,379 |
| Symmetry imposed | C1 |
| Map Resolution (Å) | 3.64 |
| FSC threshold | 0.143 |
| **Refinement & validation** | |
| Initial model used | 9BNP |
| Software | Phenix 1.21 |
| Number of residues | |
| Protein | 1,923 |
| Ligand | 122 |
| Map CC | 0.81 |
| R.m.s. deviations | |
| Bond lengths (Å) | 0.004 |
| Bond angles (˚) | 0.865 |
| EMRinger score | 2.26 |
| MolProbity score | 1.44 |
| Clashscore | 3.34 |
| Rotamer outliers (%) | 0 |
| Ramachandran plot | |
| Favored (%) | 95.4 |
| Allowed (%) | 4.6 |
| Outliers (%) | 0 |

**Table S3. Cryo-EM data collection, processing, and refinement validation statistics**
